# Prefrontal 5α-reductase 2 mediates male-specific acute stress response

**DOI:** 10.1101/2024.05.07.593076

**Authors:** Roberto Cadeddu, Giulia Braccagni, Gabriele Floris, Caterina Branca, Eleonora Corridori, Sara Salviati, Pilar Sánchez, Luca Spiro Santovito, Jesus M Torres, Esperanza Ortega, Graziano Pinna, Philip J. Moos, Simona Scheggi, Marco Bortolato

## Abstract

A key response to acute stress is the increased brain synthesis of the neurosteroid allopregnanolone (AP). While the rate-limiting step of this reaction is catalyzed by 5α-reductase (5αR), the role of its two primary isoenzymes, 5αR1 and 5αR2, in stress reactivity remains unclear. Here, we found that acute stress led to increased levels of 5αR2, but not 5αR1, in the medial prefrontal cortex (mPFC) of male, but not female, rats. Downregulation of 5αR2 in the mPFC significantly reduced stress response in males, and similar sexual dimorphic effects were observed in a novel line of 5αR2 knockout rats. Notably, 5αR1 regulated baseline AP synthesis, while 5αR2 enabled AP production under stress. Acute AP administration restored stress response in 5αR2 knockdown rats. Single-nucleus transcriptomics showed that 5αR2 enabled stress-induced protein translation in neurons and glia. These results highlight the crucial role of 5αR2 in mediating sex-specific differences in acute stress reactivity.

## INTRODUCTION

The acute stress response is a transient physiological and psychological reaction elicited by an immediate perceived threat or demand. It manifests as a heightened activation of the sympatho-adrenomedullary system and the hypothalamic-pituitary-adrenal (HPA) axis, aimed at mobilizing the organism’s energy resources to cope with immediate situational demands (*1*). Despite its short duration, acute stress can have significant physiological and psychological impacts, insofar as it influences cognitive function and emotional regulation (*2*, *3*). Notably, while stress-induced signals are initially adaptive, they can exacerbate health risks and contribute to the development of stress-related disorders when excessively activated or dysregulated.

Glucocorticoids, catecholamines, and neurotrophins have long been recognized for their pivotal roles in orchestrating the physiological and behavioral aspects of adaptation to acute stress; additionally, emerging research has shed light on the significance of allopregnanolone (3α-hydroxy-5α-pregnan-20-one; AP), a neurosteroid derivative of progesterone, which exhibits potent anxiolytic and neuroprotective properties (*4*). In male rodents, exposure to acute stressors increases AP levels in the prefrontal cortex (PFC) (*5–7*). Given the primary role of this brain region in coping and resilience (*8*), this phenomenon has been extensively interpreted as an adaptive mechanism to reinstate homeostasis by mitigating the excessive activation of other systems implicated in the acute stress response, such as the HPA axis (*4*). Indeed, AP acts as a positive allosteric modulator of GABA-A channels (*9*), the most prominent inhibitory neurotransmitter receptors in the central nervous system.

Several details remain elusive with respect to the mechanisms that enable AP biosynthesis in response to acute stress. The conversion of progesterone to AP occurs through a two-step process. Initially, progesterone is transformed into dihydroprogesterone (DHP) by the enzyme 5α-reductase (5αR), followed by the conversion of DHP into AP by the enzyme 3α hydroxysteroid oxidoreductase (3αHSOR) (Fig. S1A) (*10*). Unlike 3αHSOR, 5αR catalyzes a unidirectional reaction, regarded as the key rate-limiting step in AP synthesis (*11*). In line with this evidence, 5αR inhibitors produce a profound reduction in AP synthesis (*12*).

Notably, recent evidence indicates that acute administration of finasteride, a prototypical 5αR inhibitor, hinders the capacity to mount an effective adaptive response to both stressful and incentive stimuli (*13*), suggesting that AP may be relevant in promoting coping and shaping the broader behavioral response to acute and salient challenges. The two most extensively characterized 5αR isoenzymes, 5αR1 and 5αR2, share similar structures and genetic ancestry (*11*). Despite this, their substrate affinity and distribution differences throughout the body suggest distinct physiological roles. For instance, 5αR2 exhibits a markedly higher affinity for testosterone and progesterone than 5αR1 (*11, 14*).

Accordingly, 5αR2 is implicated in the peripheral conversion of testosterone into its more active metabolite, 5α-androstan-17β-ol-3-one (dihydrotestosterone; DHT), a potent androgen hormone responsible for developing numerous secondary sexual characteristics in males. While both isoenzymes are expressed in the brain (*11,15*), 5αR2 shows significantly lower levels in the PFC of male and female rats, along with a sexually dimorphic regulation exerted by androgens (*16,17*). Collectively, the functional role of this enzyme remains less comprehensively understood. Our study exposed male and female rats to various acute stressors to elucidate the specific roles of these 5αR isozymes in stress-induced AP synthesis and behavioral reactivity. Additionally, we specifically suppressed the expression of each isoenzyme in the medial (m) PFC and nucleus accumbens (NAc), two crucial regions in the regulation of reward, motivation, and mood (*18,19*). Then, we assessed the impact of this manipulation on behavior, AP synthesis, and cell-specific transcriptomic alterations.

In contrast to previous assumptions (*20*), our results identify 5αR2, rather than 5αR1, as the primary contributor to the acute synthesis of AP in the mPFC of male rats. Specifically, our observations reveal that acute stress rapidly upregulates 5αR2 expression in the mPFC, and this increase is instrumental for the production of AP and the enactment of several behavioral responses to salient stimuli. This distinctive response is accompanied by marked transcriptomic alterations in PFC pyramidal neurons, with 5αR2 deficiency resulting in a generalized reduction of the acute stress response across most cell types. These effects were observed exclusively in males under stressful conditions, indicating a significant sexual dimorphism in the role of 5αR2 in behavioral regulation. This distinction provides valuable insights into the potential contribution of this enzyme to sex-specific stress susceptibility and may also yield valuable insights into the mechanisms underlying sex differences in the vulnerability to psychiatric disorders characterized by aberrant stress responses.

## RESULTS

Below is a summary of the main results; a comprehensive description of all findings, including statistical details, can be found in the supplementary results.

### Acute stress upregulates 5**α**R2 in males

To assess the impact of acute stress on 5αR levels, we employed forced swim and footshock, two of the best-established acute stressors in laboratory rats. These procedures have been thoroughly validated for their capacity to produce consistent and measurable stress responses across various behavioral, physiological, and neurobiological parameters, including the elevation of AP in the mPFC. We confirmed that both stressors elevated the levels of AP and its immediate precursor, DHP, within 30 minutes of their conclusion (Fig. S1). The DHP/progesterone ratio (signifying 5αR activity) was also significantly elevated, indicating increased 5αR activity (Fig. S1).

We then examined the effects of acute stress on the levels of 5αR1 and 5αR2 in different brain regions before, immediately after, and 30 minutes following forced swim. Figure S2 shows that this stressor increased 5αR2 transcript levels in the mPFC and hippocampus of male rats, whereas 5αR1 levels remained unchanged. No significant changes were observed in the amygdala or hypothalamus. Evaluation of protein expression through Western blot analyses (representative images are provided in Fig. S3) revealed that forced swim resulted in a significant upregulation of 5αR1 in the NAc and hypothalamus (Fig. 1A) at 30 minutes after the completion of forced swim. In contrast, footshock reduced 5αR1 levels in the NAc and hippocampus (Fig. 1B). Forced swim rapidly increased the concentrations of 5αR2 in the mPFC and hypothalamus, coupled with a modest yet significant reduction in amygdala (Fig. 1C). Notably, footshock led to a similar elevation in 5αR2 expression in the mPFC (Fig. 1D), but not in other brain regions. In contrast with males, females showed no alterations in the 5αR isoenzyme expression levels in response to either stressor (Fig. 1E-H).

**Figure 1:**
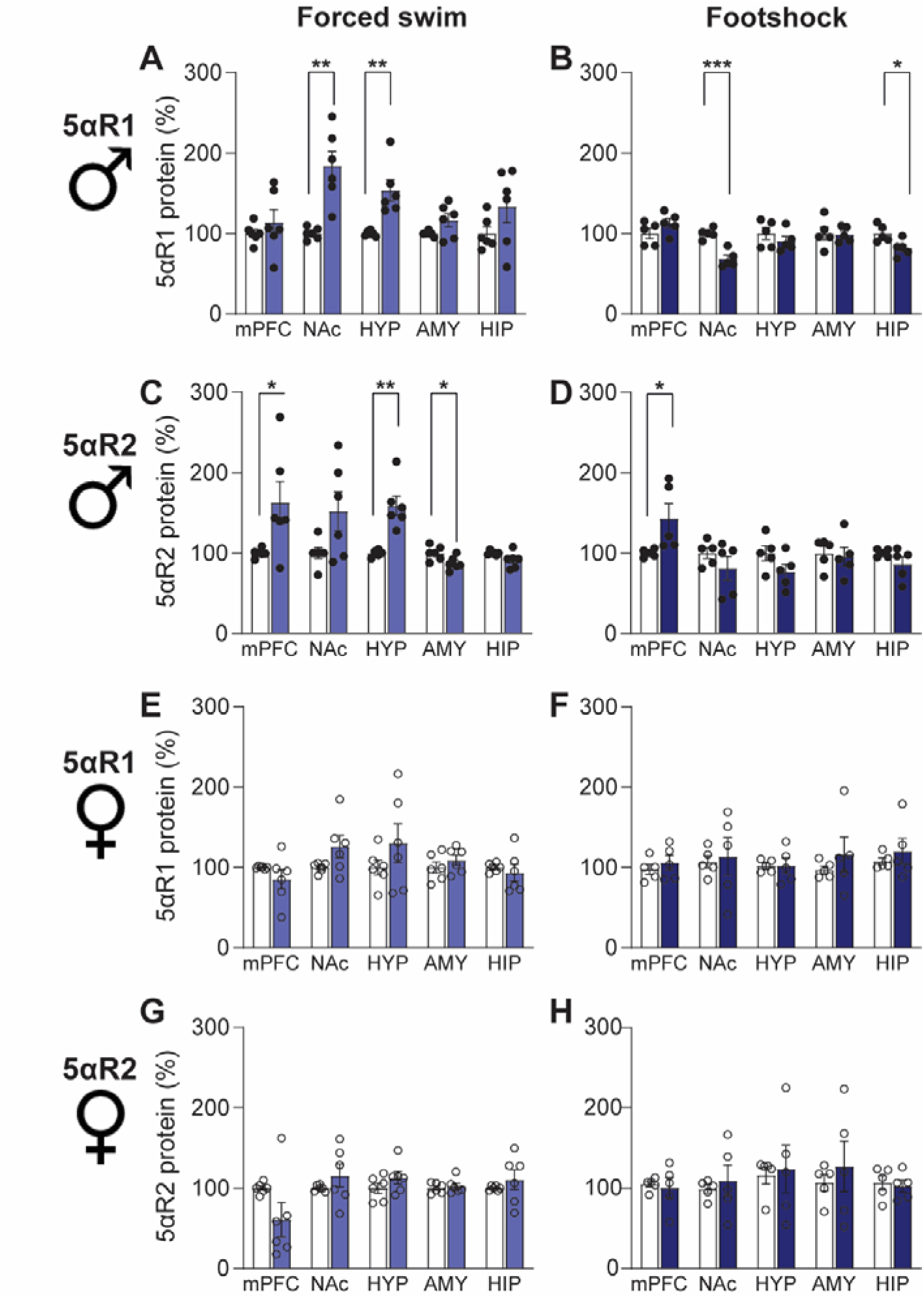
Stress upregulates 5αR2 in males. Stressed rats (represented by colored columns) were exposed to either the forced swim test (left panels) or footshock (right panels) and were sacrificed 30 minutes afterward. Western blot was used to evaluate the protein levels of 5αR1 and 5αR2 in the medial prefrontal cortex (mPFC), nucleus accumbens (NAc), hypothalamus (HYP), amygdala (AMY), and hippocampus (HIP). Control rats (represented by white columns) remained undisturbed in their home cages. In male rats, forced swim exposure significantly increased 5αR1 levels in the NAc and HYP, while footshock led to a decrease in 5αR1 levels in the NAc and HIP. Additionally, 5αR2 levels were elevated by forced swim in the PFC and HYP but decreased in the AMY. Footshock exposure increased 5αR2 levels in the mPFC of male rats. For female rats, neither forced swim nor footshock significantly affected 5αR1 or 5αR2 protein levels. For further details, refer to Supplementary Results. Data are presented as mean ± SEM (n=6/group). Light blue indicates forced swim, dark blue indicates footshock, full dots represent males, and empty dots represent females. *, p<0.05; **, p<0.01 for comparisons indicated by brackets.

To further confirm and examine anatomical details of these findings, we conducted immunofluorescence analyses of 5αR1 and 5αR2 distribution in the mPFC (Fig. S4). 5αR1 was homogeneously distributed across all compartments of the PFC and associated with both neurons and non-neuronal cells. In line with previous findings (*15*), 5αR2 was primarily found in the somata and neurites of pyramidal cells of the prelimbic and infralimbic cortex (Fig. S4). The quantification of immunofluorescent staining confirmed that the highest expression of 5αR2 was in the prelimbic cortex, but forced swim increased the expression in both pre- and infralimbic cortex of male rats (Fig. S5).

### 5αR2 is necessary to mount a response to stress in males

Building on these findings, we next characterized the specific effects of 5αR1 or 5αR2 in the mPFC and NAc, given that these two brain regions exhibited the most prominent changes in the expression of these enzymes in response to acute stress. To this end, we knocked down the expression of either isoenzyme by subjecting male and female rats to stereotaxic bilateral injections of adeno-associated serotype 5 viral vectors (AAV5) carrying shRNA constructs directed against 5αR1 or 5αR2, along with the GFP reporter gene. Scrambled RNA was used as control. This procedure effectively and selectively reduced the protein levels of the enzyme of interest, as validated by the measurement of protein levels (Fig. S6). Furthermore, the distribution of the spread of the construct diffusion was confirmed by assessment of the expression of the GFP reporter used for monitoring virus distribution (Fig. S6). Two weeks after surgery, we performed a comprehensive behavioral assessment in these animals, which included evaluating locomotor activity, forced swim immobility (representing susceptibility to acute stress), defensive withdrawal (to examine avoidant behaviors), olfactory arousal (a novel test developed by our group to assess exploration of new odorant stimuli; see validation and details in Fig. S7), social interaction (to assess natural sociability), and sucrose preference (to evaluate the response to palatable stimuli).

Examination of the phenotypes induced by 5αR1 knockdown in either mPFC or NAc of male rats showed that this manipulation yielded minimal impact (Fig. 2A-L), except for an increase in locomotor activity following mPFC targeting (Fig. 2A) and decreased social interaction following targeting of either the mPFC (Fig. 2E) or the NAc (Fig. 2K). Conversely, 5αR2 knockdown in the mPFC of male rats compromised their ability to appropriately respond to acute exposure to salient (either arousing or stressful) stimuli (Fig. 2M-R). Notably, these animals did not exhibit noticeable alterations in locomotion (Fig. 2M), yet they exhibited prolonged immobility during the forced swim test (Fig. 2N) and a reduced tendency to exit the dark chamber in the defensive withdrawal paradigm (Fig. 2O). Additionally, they displayed diminished responsiveness to novel olfactory stimuli, as evidenced by decreased exploration towards a new scent (Fig. 2P), and reduced interaction time with unfamiliar counterparts (Fig. 2Q). Furthermore, male rats did not show sucrose preference over time (Fig. 2R). Notably, most of these phenotypes were specific to males with 5αR2 knockdown in the mPFC; targeting the NAc with the same RNA interference construct did not produce this range of effects (Fig. 2S-X), except for reduced social interaction (Fig. 2W) and sucrose preference (Fig. 2X). Additionally, the same manipulation in females resulted in no significant alterations (Fig. 3), except for a significant increase in immobility in the forced swim test following 5αR1 knockdown in the mPFC (Fig. 3A).

**Figure 2:**
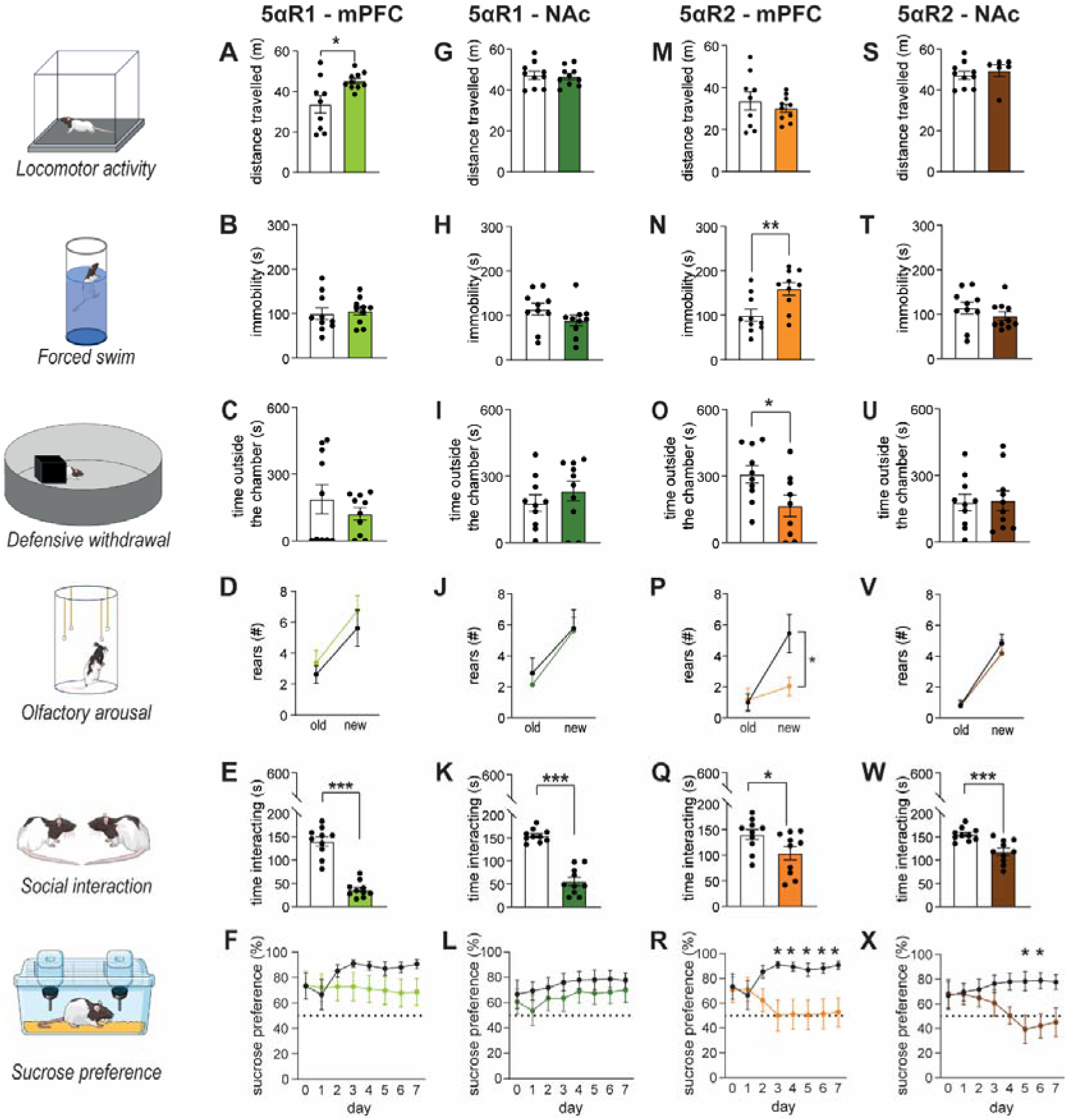
Behavioral characterization of male animals. Male rats received stereotaxic injections of either scrambled-shRNA or shRNA targeting 5αR1 (A-L) or 5αR2 (M-X) in the medial prefrontal cortex (mPFC, light colors) or nucleus accumbens (NAc, dark colors), and their behavior was evaluated two weeks post-surgery. (A-F) For 5αR1 knockdown (KD) in the mPFC, locomotor activity increased, but no differences were observed in the forced swim test, defensive withdrawal, new odor exposure, or sucrose preference tests. However, KD rats interacted less with unfamiliar animals compared to controls. (G-L) When 5αR1 was knocked down in the NAc, no changes were observed in locomotor activity, forced swim, defensive withdrawal, or in odor arousal. Similar deficits were found in social interaction with unfamiliar animals, but there were no differences in sucrose preference. (M-R) For 5αR2 KD in the mPFC, KD rats showed increased immobility in the forced swim test and spent less time outside the chamber during defensive withdrawal. They also showed reduced arousal to new odor exposure and reduced interaction with unfamiliar animals, but no changes in locomotor activity. In the sucrose preference test, KD rats showed no preference over time. (S-X) When 5αR2 was knocked down in the NAc, no significant changes were found in locomotor activity, forced swim, or defensive withdrawal. However, KD rats interacted less with unfamiliar animals and showed a lack of sucrose preference at specific time points. For further details, refer to Supplementary Results. Data are presented as mean ± SEM (n=6-10/group). *, p<0.05, **, p<0.01, and ***, p<0.001 for comparisons indicated by brackets or against controls at the same time point.

**Figure 3:**
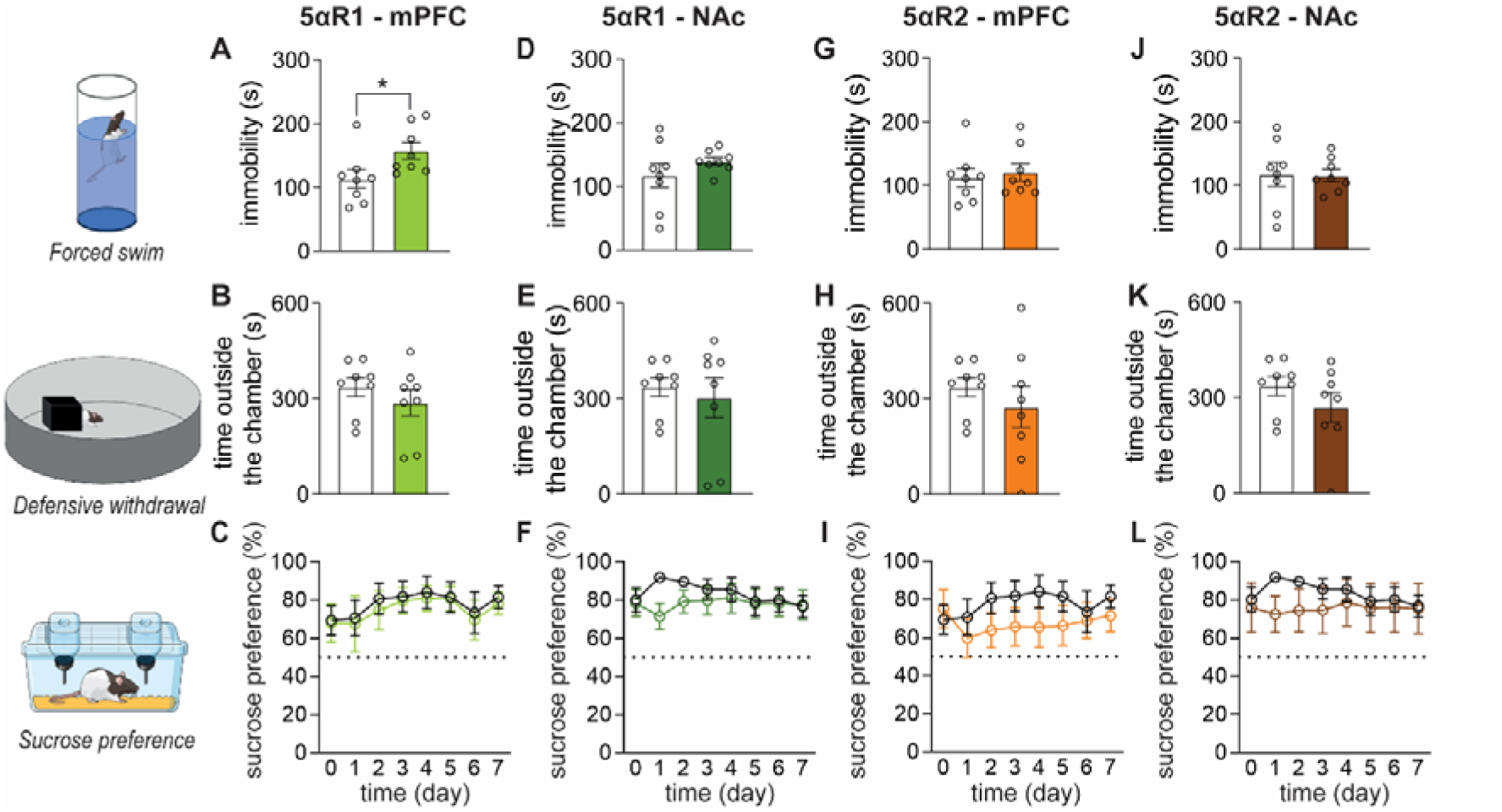
Behavioral characterization of female rats. Female rats received stereotaxic injections into the prefrontal cortex (mPFC) or nucleus accumbens (NAc) with constructs carrying scrambled-shRNA, or shRNA targeting 5αR1 or 5αR2, and their behavior was evaluated two weeks post-surgery. (A-C) Knockdown (KD) of 5αR1 in the PFC increased immobility time in the forced swim test, while no effects were observed in the defensive withdrawal paradigm or sucrose preference test. (D-F) KD of 5αR1 in the NAc did not lead to any significant changes in the tested behaviors. (G-L) Similarly, KD of 5αR2 did not induce any significant differences, regardless of whether the mPFC or NAc was targeted. For further details, refer to Supplementary Results. Data are presented as mean ± SEM (n=6-8/group). *p<0.05 for comparisons indicated by brackets.

To confirm the contribution of 5αR2 to the regulation of environmental reactivity, we generated 5αR2 full-body knockout (KO) rats by CRISPR/Cas9 (see Fig. 4A-B for generation and validation). The loss-of-function mutation of 5αR2 in humans has been shown to impair the normal development of male external genitalia (*21*). In line with this finding, we documented that male KO rats displayed a significant reduction in the anogenital distance (Fig. S8) – a reliable index of abnormal masculinization of the reproductive tract - and underdeveloped vesicular glands and testicles. Interestingly, behavioral analyses showed that male KO rats showed a significant increase in immobility time during the forced swim test (Fig. 4C) and a decrease in time spent outside the chamber in the defensive withdrawal paradigm (Fig. 4D), as well as sucrose preference (Fig. 4E). In line with our findings in knockdown rats, these abnormal phenotypes were not detected in females (Fig. 4F-H).

**Figure 4:**
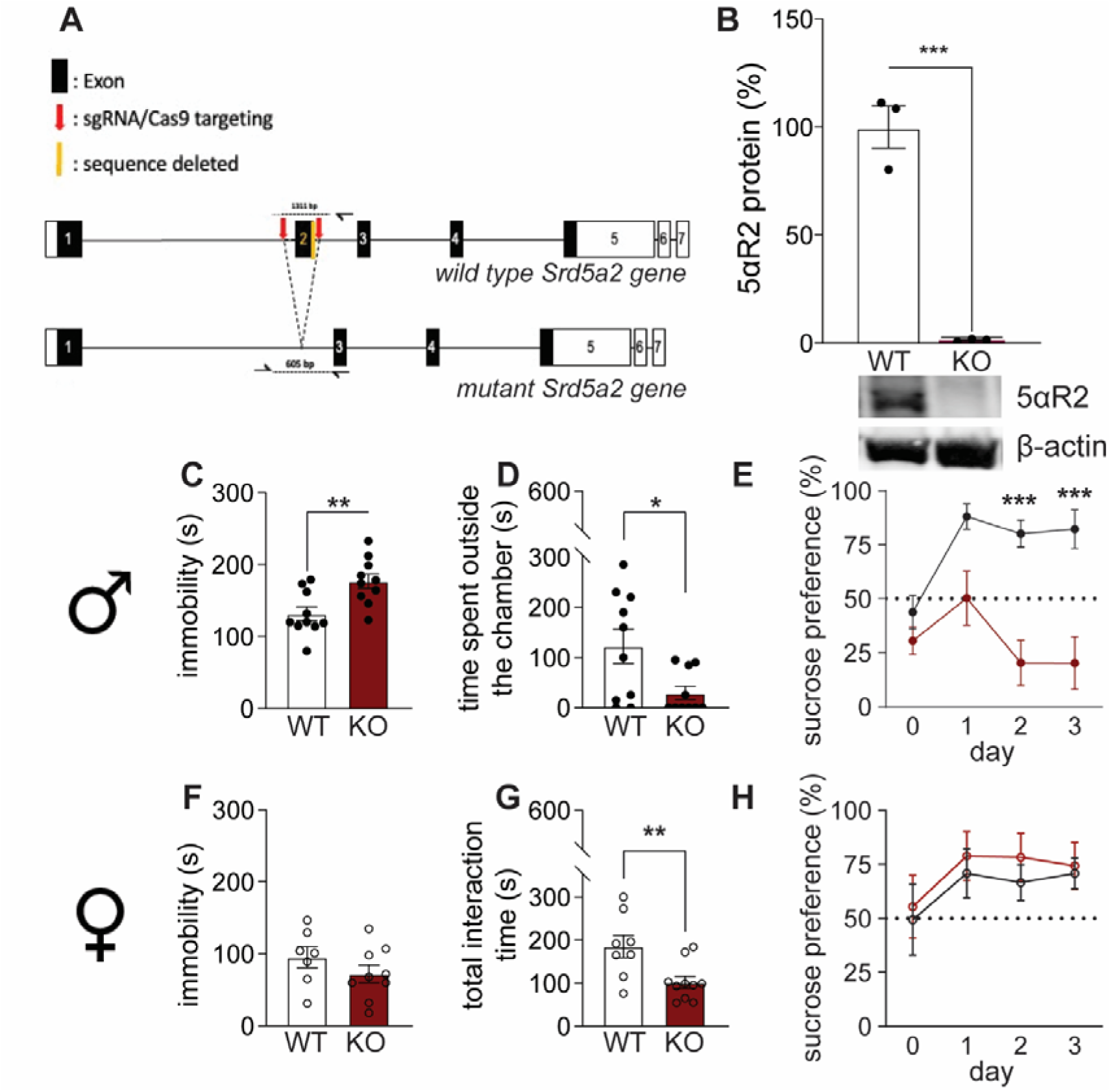
Generation and behavioral characterization of 5αR2 knockout (KO) rats. (A-B) The 5αR2 gene was targeted using CRISPR/Cas9 genome editing to remove exon 2. Validation of the KO model was performed by evaluating 5αR2 levels in prostate tissue using Western blot (representative image shown below panel B), confirming the efficient deletion and absence of 5αR2 expression. (C-E) KO male rats exhibited increased floating time in the forced swim test, reduced time spent outside the chamber in the defensive withdrawal paradigm, and decreased sucrose preference. (F-H) KO female rats showed fewer changes, with no significant differences in the forced swim and sucrose preference tests but a decrease in interaction time with unfamiliar counterparts. For further details, see Supplementary Results. Data are presented as mean ± SEM (n=10/group). *, p<0.05; **, p<0.01; ***, p<0.001 for comparisons indicated by brackets or against controls at the same time point.

### 5αR2 enables stress-induced AP synthesis

To understand whether the difference in behavioral regulation and stress sensitivity of the two 5αR isoenzymes were paralleled by different enzymatic functions, we measured the levels of progesterone and AP (as well as their relative ratio) in the PFC in basal conditions and after stress exposure. Male rats knockdown for either 5αR1 or 5αR2 were sacrificed 30 minutes after exposure to forced swim stress, in comparison with no-stress conditions. At baseline, knocking down 5αR1 increased progesterone levels and decreased AP levels, while knocking down 5αR2 reduced the content of both progesterone and AP (Fig. 5A-B). Notably, the AP/progesterone ratio was significantly reduced by the knockdown of 5αR1, but not 5αR2 (Fig. 5C). After stress exposure, we detected significant upregulation of progesterone and significant downregulation of AP levels in 5αR2 knockdown animals (Fig. 5D-E), leading to a significant downregulation of the ratio AP/progesterone (Fig. 5F). Taken together these data suggest that while 5αR1 is crucial for maintaining the proper levels of progesterone/AP at baseline (tonic phase), 5αR2 presence is instrumental to increase AP during stress exposure (inducible phase).

**Figure 5:**
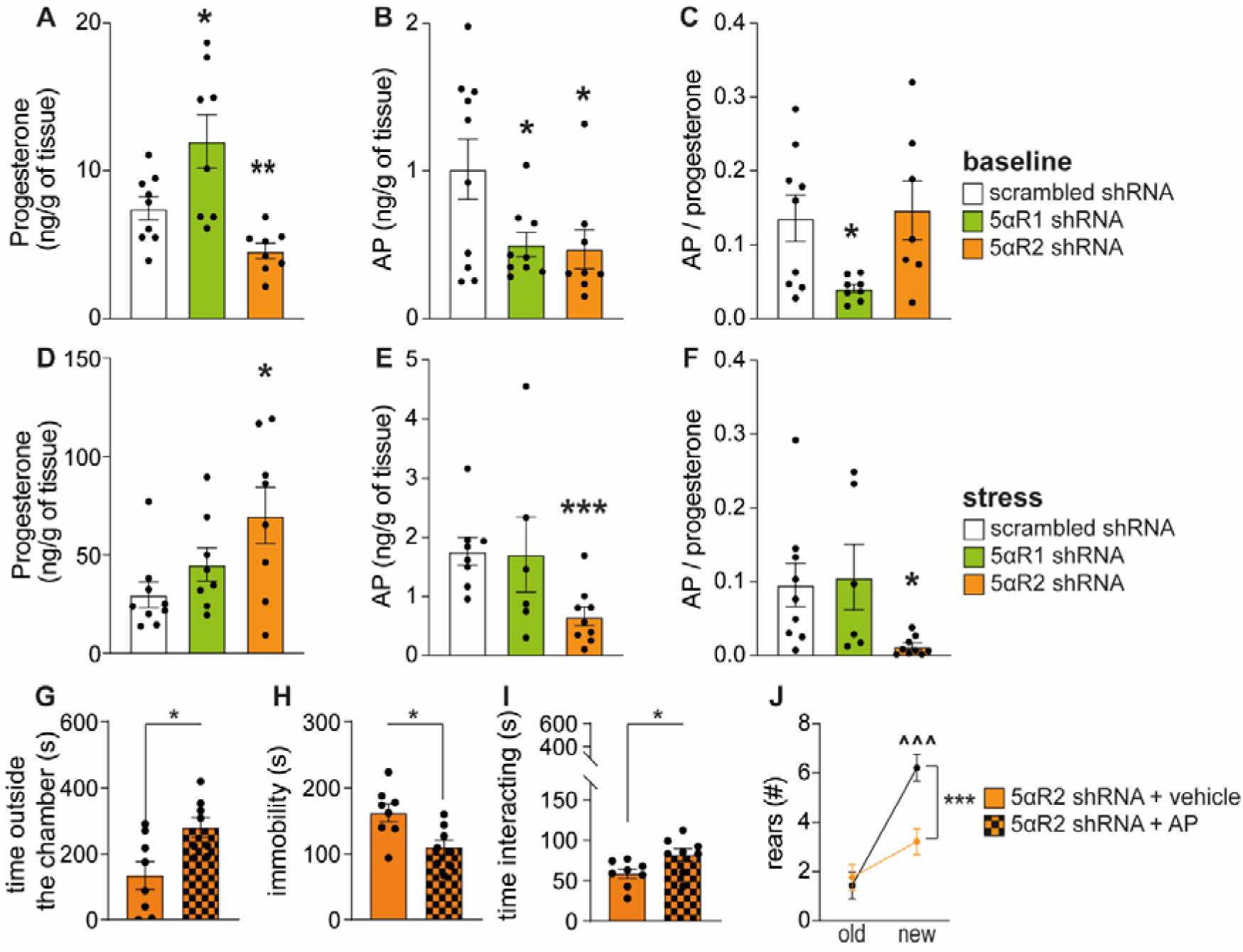
Levels of neuroactive steroids in rat prefrontal cortex and effects of systemic administration of allopregnanolone in 5αR2 deficient rats. (A-F) Analysis of progesterone and allopregnanolone (AP) levels in the prefrontal cortex of experimental animals (scrambled-shRNA, shRNA targeting 5αR1, and shRNA targeting 5αR2) was conducted under basal conditions (A-C) and 30 minutes post-stress (forced swim) (D-F). (A-C) Under basal conditions, shRNA targeting 5αR1 significantly increased progesterone levels, whereas shRNA targeting 5αR2 significantly decreased them. AP levels were reduced by the knockdown of either 5αR1 or 5αR2. Notably, only 5αR1 downregulation reduced the ratio of AP to progesterone. (D-F) After stress exposure, both progesterone and AP levels increased, but the AP/progesterone ratio remained unchanged. In animals with 5αR2 downregulation, stress significantly elevated progesterone levels while reducing AP levels, affecting the AP/progesterone ratio. No significant changes were observed in animals with 5αR1 downregulation. Data are presented as mean ± SEM (n=6-10/group) *, p<0.05; **, p<0.01; ***, p<0.001 for comparisons vs the scrambled-shRNA samples. (G-J) To explore the effects of 5αR2 downregulation, we tested whether AP treatment could rescue the observed phenotype. Treatment with AP (6 mg/kg, IP, 15 minutes before testing) increased the time spent outside the chamber in the defensive withdrawal test, decreased immobility in the forced swim test, enhanced social interaction (measured as total interaction time), and increased odor arousal. For further details, refer to Supplementary Results. Data are presented as mean ± SEM (n=8/group). *, p<0.05; **, p<0.01; ***, p<0.001 for comparisons indicated by brackets. ^^^, p<0.001 for comparison between new and old odor for the AP treated animals.

These data led us to hypothesize that the inability to synthesize AP in response to acute stress may underlie the impaired stress response in male rats with 5αR2 knockdown in the mPFC. To test this, we reevaluated these animals’ behavior following AP treatment (6 mg/kg, IP, 15 minutes before testing). Notably, AP treatment improved their behavioral abnormalities (Fig. 5G-5J, Fig. S9). Specifically, male rats spent more time outside the dark chamber in the defensive withdrawal paradigm (Fig. 5G, Fig. S9), reduced immobility in the forced swim test (Fig. 5H, Fig. S9), increased social interaction (Fig. 5I, Fig. S9), and new odor arousal (Fig. 5J).

### 5αR2 enables RNA translation processes in the mPFC in response to acute stress

To understand the cellular mechanisms underlying the regulation of stress reactivity by 5αR2 in the mPFC, we conducted snRNA-seq of this region in male rats subjected to either scrambled-RNA or 5αR2-shRNA administration. Specifically, to elucidate the selective impact of this enzyme on molecular changes induced by acute stress, different groups of animals were either exposed to forced swim stress or maintained under control, no-stress conditions. Gamma-Poisson modeling identified 13 distinct clusters of cells, which were recognized via cell markers as: pyramidal neurons (PN) from various cortical layers; interneurons (VIP positive and VIP negative); oligodendrocyte progenitor cells (OPCs); oligodendrocytes; astrocytes; microglia; stromal cells; and endothelial cells (Fig. 6A-B, S10). To elucidate the effects of 5αR2 downregulation in the mPFC under both baseline conditions and in the context of acute stress response, we compared snRNA-seq data between unstressed 5αR2 knockdown and control rats. Notably, no discernible differences in gene expression were observed in the interneuron, stromal, and endothelial cell clusters, as well as most PN clusters (Data S1-S3). In contrast, layer-V PNs, microglia, astrocytes, OPCs, and oligodendrocytes exhibited substantial transcriptomic variations between groups, suggesting their heightened susceptibility to the effects of 5αR2 deficiency.

**Figure 6:**
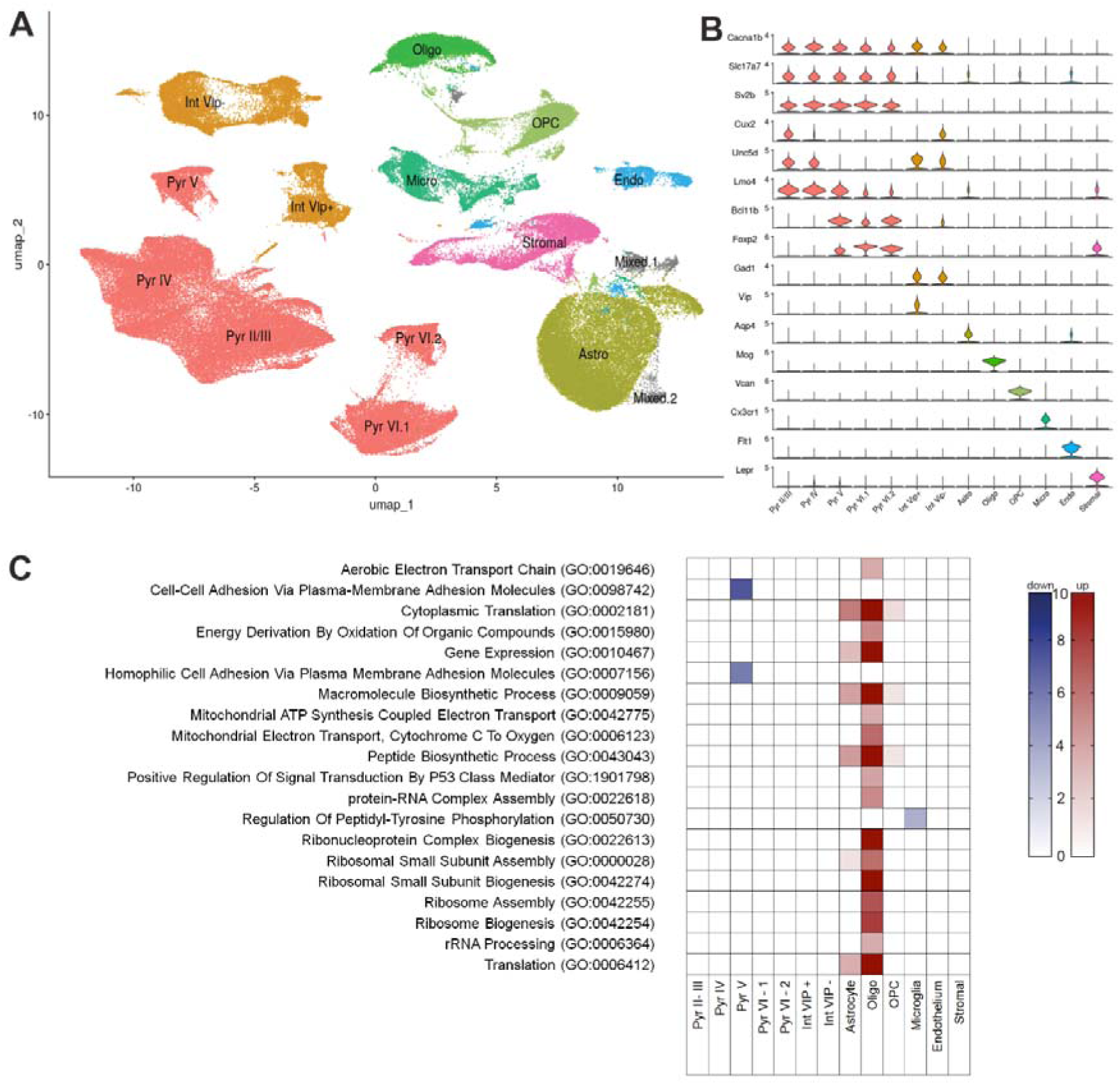
Single-nucleus transcriptomic analyses of cortical tissues of 5αR2-shRNA and scrambled-shRNA rats. (A) Uniform Manifold Approximation and Projection (UMAP) following SCTransformation with the generalized linear model, Gamma Poisson method, annotated by major cell types, including astrocytes, oligodendrocytes, microglia, pyramidal neurons, interneurons, oligodendrocyte progenitor cells (OPC), and others. (B) Violin plots of marker genes used to distinguish the clusters differentiating known cell types. (C) Gene ontology analysis for biological processes highlights the most differential biological processes among these cell clusters. Abbreviations: Pyr, pyramidal neurons; Int, interneurons; Astro, astrocytes; Oligo, oligodendrocytes; OPC, oligodendrocyte precursor cell; Micro, microglia; Endo, endothelial cells.

Within the oligodendrocyte and astrocyte clusters, 164 and 48 differentially expressed genes (DEGs) were identified, respectively. Gene Ontology (GO) analysis revealed marked upregulations in multiple genes associated with cytoplasmic translation and peptide biosynthetic processes (Fig. 6C), along with increased expression of transcripts associated with ribosomal subunits and RNA binding (Fig. S11). Astrocytes also exhibited an upregulation of 48 DEGs, with significant enrichment in cytoplasmic translation, macromolecule biosynthesis (Fig. 6C), and ribosomal subunits (Fig. S11). Conversely, the 76 DEGs identified in layer V pyramidal neurons featured a notable downregulation in pathways related to “Cell-Cell Adhesion Via Plasma-Membrane Adhesion Molecules”, “Homophilic Cell Adhesion” (Fig. 6C), “Neuron projection”, and “Cell Adhesion mediator activity” (Fig. S11). Coexpression analyses by high-dimensional Weighted Gene Co-expression Network Analysis (hdWGCNA) across all these cell types pointed to significant changes in modules enriched in DEGs associated with bioenergetics, such as ATP synthesis, and cytoplasmic translation (Figs. S12-14 and Data S4-S6). In addition, astrocytes of knockdown rats featured upregulations in modules associated with aspartate transmembrane transport, implying enhanced activity in these cells (Fig. S13 and Data S5) Finally, layer V pyramidal cells of knockdown rats exhibited upregulations in modules enriched for dendrite morphogenesis and glutamate receptor signaling, again indicating modifications in dendrite arborization, calcium transport into cytosol and dendrite morphogenesis were found in layer V pyramidal neurons, likely indicating increased excitation (Fig. S14 and Data S6).

Given that our analyses revealed that most effects of 5αR2 are best captured in relation to the response to acute stress, we conducted a detailed analysis of snRNA-seq data with respect to the effects of forced swim stress in rats previously subjected to mPFC 5αR2 knockdown or scrambled RNA injections. To initially determine the transcriptomic effects of acute stress in control rats, we ranked the overall GO biological processes in the scrambled RNA groups (either unstressed or 30 minutes post-stress exposure). Next, we analyzed the differences in transcriptomic profiles in unstressed and stressed 5αR2-shRNA groups to evaluate the specific impact of this enzyme on acute stress response. While stress exposure significantly upregulated several processes related to translation and gene expression in both neurons and glial cells (Fig. 7A, left panel), a notable downregulation of these pathways was observed in tissues with 5αR2 knockdown (Fig. 7A, right panel). Specifically, in the mPFC neuron clusters, the effects were driven by 262 DEGs in the control group (Data S7-S9) and 1191 DEGs in the 5αR2-shRNA group (Data S10-S12). Additionally, the downregulation of pathways such as “Chemical Synaptic Transmission” and “Nervous System Development” in interneurons was entirely abolished by 5αR2 deficiency. Similar patterns were found in the GO cellular component analysis (Fig. 7B) and GO molecular function analysis (Fig. S15). Notably, coexpression analyses across most neuronal clusters revealed that stress produced similar changes in mitochondrial ATP synthesis and oxidative phosphorylation in the top modules of both 5αR2-knockdown and controls, likely suggesting that acute stress affected the bioenergetics of ATP across most cell populations, irrespective of 5αR2 deficiency (Fig. S16-S27). The same changes were observed in oligodendrocytes and OPC (Fig. S28-S31).

**Figure 7:**
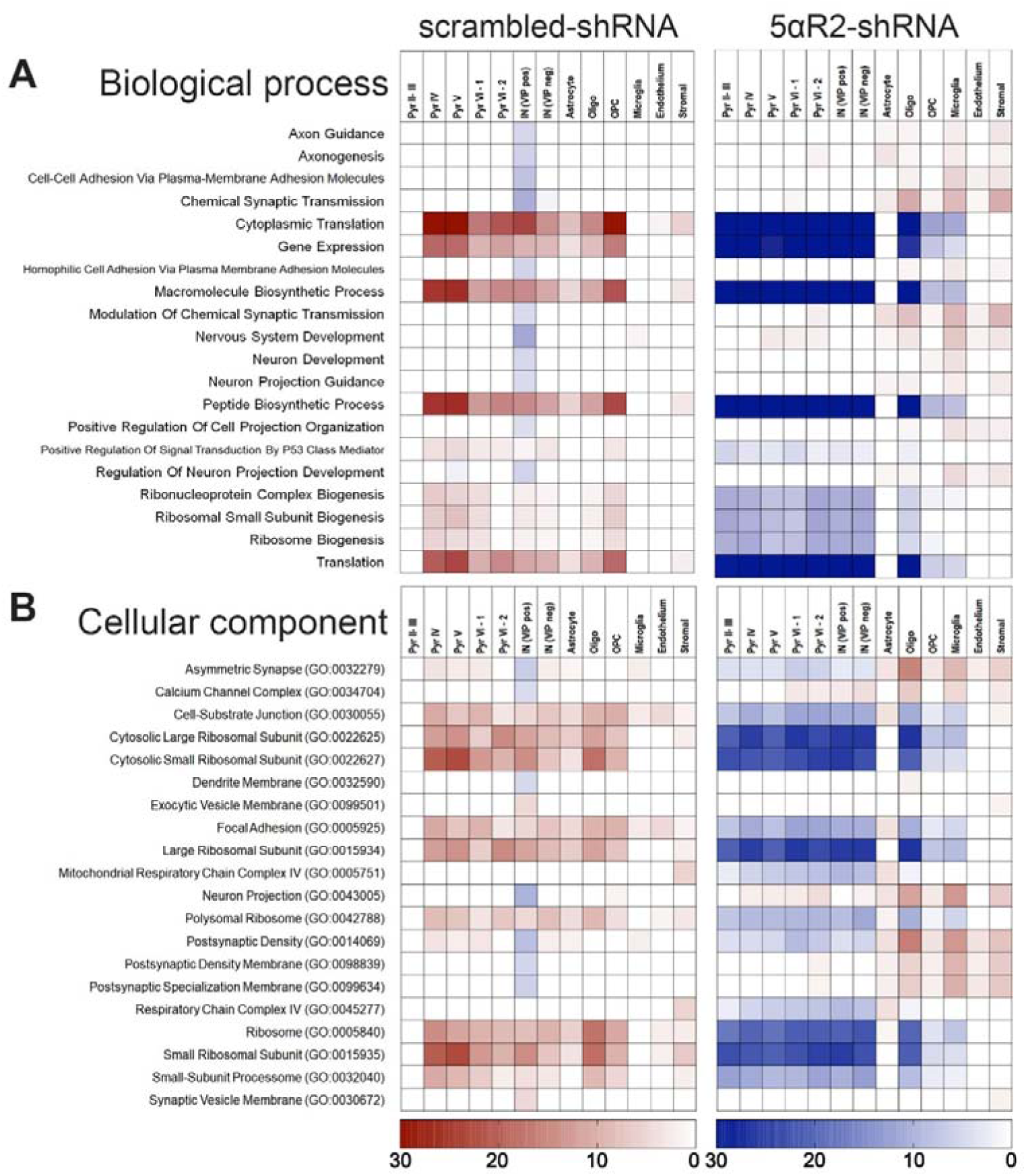
Single-nucleus transcriptomic analyses of cortical tissues of 5αR2-shRNA and scrambled-shRNA rats exposed to acute stress. (A-B) Gene ontology analysis for biological process (A) and cellular component (B) highlighting the most differential ontologies among these cell clusters. The two comparisons are shown side by side; on the left, differences induced by stress exposure in the prefrontal cortex (mPFC) of control rats; on the right, the same behavioral paradigm applied to rats knocked down for 5αR2 in the mPFC. Abbreviations: Pyr, pyramidal neurons; Int, interneurons; Oligo, oligodendrocytes; OPC, oligodendrocyte precursor cell.

## DISCUSSION

The results of our study point to a paramount role of prefrontal 5αR2, rather than 5αR1, in enabling acute stress response exclusively in male subjects. This conclusion is supported by several key findings: first, 5αR2 expression was rapidly upregulated in the mPFC in response to two distinct, well-established experimental protocols producing inescapable acute stress; however, this response was exclusively observed in males, in sharp contrast with the lack of changes in females; second, in contrast with 5αR1, targeted genetic downregulation of 5αR2 expression in this region led to a profound reduction in the ability of male, but not female, rats to mount an adaptive behavioral response to both stressful and arousing stimuli; third, a selective reduction of 5αR2, but not 5αR1, abolished stress-induced elevation of AP levels in males, identifying this enzyme as a pivotal factor for the biosynthesis of this neurosteroid in response to acute stress. In support of this idea, AP administration reversed the behavioral deficits observed in male rats subjected to 5αR2 knockdown in the mPFC. Finally, 5αR2 suppression in this region led to a complete reversal of the cellular response to stress in males, as evidenced by single-nucleus transcriptomic analyses. Overall, these results underscore 5αR2’s central role in regulating behavioral reactivity in males but not in females and point to the implication of this enzyme in sexually dimorphic regulation of the response to acute stress. These findings align with our prior research indicating a diminished reactivity to environmental stimuli following finasteride administration (*14*) by pointing to the role of 5αR2, rather than 5αR1, inhibition as critical to blunt the ability to react to salient stimuli. Most importantly, these results challenge established beliefs regarding the functions of 5αR1 and 5αR2 in behavioral regulation. While both isoforms are expressed in the brain, 5αR1 has frequently been considered the predominant functional variant, proposed to act as a constitutive enzyme with catabolic and neuroprotective functions. In contrast, 5αR2 has generally been regarded as playing an ancillary role in the brain, possibly restricted to early developmental stages (*20*).

Our findings on the effects of acute stress on the expression of both 5αR isoenzymes showed that, while both footshock and forced swim elicited an increased 5αR2 synthesis in the mPFC, these two stressors led to a range of region- and sex-dependent responses in the expression of these isoenzymes. For example, forced swim stress led to a significant upregulation of 5αR1 in the NAc and hypothalamus of males. Conversely, footshock decreased 5αR1 expression in the NAc and hippocampus. These changes were not observed in females. In addition, forced swim stress also upregulated 5αR2 expression in the hypothalamus of males. This complex array of sex-specific changes suggests a complex regulatory mechanism governing the expression of 5αR isoenzymes under different forms of stress, which might involve both transcriptional and post-transcriptional controls. The differential regulation of 5αR1 and 5αR2 in the mPFC, NAc, hypothalamus, hippocampus, and amygdala may reflect region-specific roles in the modulation of neurosteroid synthesis and, consequently, stress reactivity. Notably, our results contrast with the findings of previous studies indicating elevated levels of both 5αR isoenzymes in the mPFC following forced swim stress in Wistar rats (*22*). Furthermore, in another study on Wistar rats, restraint induced a similar elevation in both isoenzymes in males; however, stressed females showed 5αR1 down-regulation and 5αR2 up-regulation (*23*). While the causes of these discrepancies remain unknown, they may reflect differences in genetic background, time, and conditions of stress exposure. Preliminary results in our lab suggest that strain and circadian differences may influence the expression of 5αR enzymes. Future studies are needed to explore these potential factors further.

As previously delineated, AP biosynthesis proceeds from progesterone through a two-step enzymatic pathway, initially involving the transformation of progesterone into DHP via the action of 5αR, followed by the subsequent conversion of DHP into AP by 3αHSOR. Our findings indicate that, in rats subjected to knockdown of 5αR1 in the mPFC, the basal concentrations of progesterone were significantly elevated, while AP levels were reduced. However, no behavioral differences were observed between these rats and their counterparts treated with scrambled shRNA. These data suggest that 5αR1 functions as a constitutive enzyme responsible for the continuous conversion of progesterone to DHP and the sustained production of basal levels of AP. In contrast, 5αR2 knockdown in the mPFC actively abolished stress-induced augmentation of AP levels.

These findings collectively suggest that 5αR1 and 5αR2 exhibit distinct functional roles in regulating AP synthesis. These data indicate that 5αR2 orchestrates a transient, stress-adaptive response by enabling the rapid synthesis of AP in response to swift environmental variations. The characteristics of 5αR2, including its synthesis within 30 minutes from acute stress exposure, align with those of an adaptive or inducible enzyme responsive to sudden environmental changes. Consistent with this distinction, 5αR2 exhibits a markedly higher affinity for progesterone and other ketosteroids than 5αR1 (*11,14*). Furthermore, the lower and more localized distribution of 5αR2 within the brain aligns with the specific expression pattern characteristic of inducible enzymes. Steroidogenic cells do not store preformed hormones but synthesize steroids *de novo* in response to physiological demands (*24*); thus, the functional bifurcation between 5αR1 and 5αR2 appears perfectly tailored to enable either slow and sustained or rapid and transient adaptive responses to environmental contingencies. Taken together, these data highlight a distinction between the dynamics of 5αR1 and 5αR2 regarding AP production.

Notably, 5αR2 knockdown caused a reduction in baseline progesterone and AP levels, while their ratio remained unchanged. These results suggest that 5αR2 may play a role in the synthesis of progesterone from its precursors, cholesterol and pregnenolone, potentially through indirect effects mediated by other steroid products of 5αR2. Although the precise mechanisms need further investigation, the data indicate that 5αR2 affects baseline AP levels, likely influencing its precursors’ synthesis or metabolism. Additionally, baseline AP levels in 5αR2 knockdown animals were lower than in the control group. However, as the control animals were also handled and sacrificed, we cannot rule out the possibility that stress-induced changes in progesterone and AP levels occurred despite the absence of forced swim tests.

The findings regarding 5αR2 downregulation in the PFC (including its sexual dimorphism) were largely corroborated in our newly generated constitutive, full-body KO rat line. These results indicate that the loss of 5αR2 function leads to widespread alterations in stress responsiveness and that these effects are not fully compensated by neurodevelopmental processes. The null-function mutation in the *SRD5A2* gene (encoding 5αR2 in humans) is associated with a rare genetic disorder characterized by male pseudohermaphroditism (*21*). Consistent with these findings, our male KO rats exhibited gonadal hypotrophy. Interestingly, however, psychiatric disturbances have not been thoroughly investigated in individuals with this syndrome, raising the question of whether more in-depth analyses should be performed to determine whether and how 5αR2 deficiency affects emotional reactivity in humans.

One of the most significant findings of this study was that the role of 5αR2 in the acute stress response was observed only in males, highlighting a pronounced sexual dimorphism in the involvement of 5αR2 in behavioral regulation. Consistent with these results, previous research has demonstrated that acute stress does not result in an increase in AP levels in the brains of female rodents (*25*). Our identification of 5αR2 as a key player in the acute stress response in male rats underscores the specific role of this isoform in mediating such responses and suggests potential therapeutic implications for modulating AP levels in stress-related disorders. These findings offer a potential mechanistic explanation for the differences in stress vulnerability and resilience between sexes. Furthermore, they enhance our understanding of the neurochemical mechanisms underlying stress adaptation and provide potential targets for therapeutic intervention in stress-related neuropsychiatric disorders.

Although the mechanisms driving the differing responses in males and females regarding 5αR2 are not yet understood, this divergence may be influenced by the enzyme’s well-established function in converting testosterone into the more potent androgen, DHT. Complementary to this observation, it is possible that changes in 5αR2 function may affect sexual dimorphism by influencing dopaminergic responses. Previous studies from our group have documented that the 5αR2 inhibitor finasteride reduces the behavioral effects of D1 dopamine receptor stimulation in male rodents (*26,27*), which plays a key role in the role of mPFC in acute stress response (*28*). Notably, emerging evidence indicates that sex differences in stress response may be influenced by prefrontal dopamine (*29*).

Acute environmental stress triggers adaptive survival mechanisms, including changes in gene expression and physiological functions. In our single-nucleus transcriptomic analysis of mPFC tissues with 5αR2 knockdown, baseline changes were minimal and mostly observed in oligodendrocytes and deep-layer pyramidal neurons. However, 5αR2 downregulation led to significant alterations in the acute stress response. In control rats, acute stress-induced extensive transcriptional changes across neurons and glia, enhancing translational activities and activating the endoplasmic reticulum. By contrast, 5αR2 knockdown rats exhibited dramatic reductions in these biological processes following acute stress, suggesting that 5αR2 facilitates neuroprotective actions that buffer against stress-induced damage in the PFC, likely through the production of AP or other 5α-reduced neurosteroids. Insufficient 5αR2 levels, coupled with overwhelming stress exposure, result in a substantial decrease in translation and energy production. Consistent with this notion, coexpression analyses revealed that 5αR2 knockdown led to the downregulation of mitochondrial processes across most cell types, pointing to a likely impairment in bioenergetics, while compensatory upregulations occurred in cytoskeletal reorganization pathways, synaptic organization, and cellular adhesion. Taken together, these results suggest that, when exposed to acute stress, 5αR2 knockdown rats failed to upregulate pathways related to translation, gene expression, and synaptic regulation, unlike control rats. This resulted in widespread mitochondrial dysfunction and impaired synaptic plasticity in neurons, indicating that 5αR2 deficiency impairs cellular resilience and stress adaptation. In line with these ideas, AP exerts neurogenic and neuroprotective effects, which likely support homeostasis and mitigate the adverse impacts of acute stress.

Notably, knocking down 5αR2 in the PFC did not exclusively affect acute stress response, but also led to intrinsic changes in specific populations of mPFC cells, including oligodendrocytes, astrocytes, and layer V pyramidal neurons. It remains unclear how these cell types were affected by 5αR2 knockdown, given that our immunofluorescence staining indicated that this enzyme is primarily present in pyramidal neurons. Regardless of the specific contribution of 5αR2 to AP synthesis, these findings suggest that changes in cells lacking 5αR2 expression might still be influenced by its downregulation, potentially due to a deficiency in local allopregnanolone synthesized by pyramidal cells. Moreover, other neurosteroids synthesized by brain 5αR2 may contribute to the functional roles of oligodendrocytes, astrocytes, and other cell types. Future research is needed to evaluate the full range of phenotypic changes attributable to 5αR2 and to explore the roles of other products of this enzyme beyond AP.

Several limitations in this study should be acknowledged. Firstly, while our investigation provides valuable insights into the roles of 5αR1 and 5αR2 within the mPFC and NAc, it is essential to recognize that our analysis was confined to these specific brain regions and isoenzymes. We did not comprehensively explore the involvement of other critical brain regions implicated in the acute stress response, such as the hippocampus, amygdala, and hypothalamus, nor did we assess the potential contributions of other 5αR isoenzymes, such as 5αR3, or other neurosteroidogenic enzymes. Future research should investigate the involvement of these additional factors in relation to the acute stress response to provide a more comprehensive understanding of the neurobiological mechanisms underlying stress adaptation.

Secondly, while our study primarily focuses on AP, it is crucial to acknowledge that other 5α-reduced neurosteroids, such as tetrahydrodeoxycorticosterone (THDOC), may also be synthesized concurrently. These neurosteroids exert diverse effects on receptors, including but not limited to GABA-A, NMDA, and progesterone membrane receptors. Given the interconnected nature of neurosteroid signaling pathways, it is plausible that changes in 5αR2 synthesis may impact the activation of these receptors. Therefore, future studies are warranted to investigate whether alterations in 5αR2 expression are accompanied by corresponding changes in the signaling of these neurosteroid-interacting receptors, thereby providing further insight into the intricate mechanisms underlying stress response and neurological function.

Thirdly, the behavioral profile of rats subjected to 5αR2 knockdown in the mPFC aligns with several characteristics associated with chronic stress and depression-related responses, including a low preference for palatable stimuli, reductions in exploratory activity and social interaction, and blunted response to acute stressors (*30–34*). In line with this observation, it is important to note that AP plays a key role in the pathophysiology of several neuropsychiatric disorders, including anxiety and depression (*35*). In particular, AP exerts potent, rapid antidepressant properties, which have led to its recent approval as a new therapy for post-partum depression (*36*). Future studies are warranted to verify the changes in 5αR2 related to chronic stress. Prior research has shown a reduction in 5αR2 levels in rats subjected to isolation rearing (*37*), indicating a possible vulnerability of this enzyme to chronic stress. Nevertheless, caution is warranted when interpreting our present findings in terms of sex differences in susceptibility to anxiety and depressive disorders, as our study did not specifically investigate the role of 5αR2 in models related to the pathophysiology of these conditions. Clarifying how alterations in 5αR2 are involved in depression and other stress-related disorders could contribute to the development of new therapeutic strategies.

Even with these limitations, our findings have revealed that 5αR2 is a key contributor to the intricate cascade of molecular events governing the adaptive stress response, shedding light on its potential importance in neurological and psychiatric contexts. Possibly due to its low baseline expression in the brain, this enzyme has received limited attention in neurobiological research. These findings pave the way for further investigations into the molecular mechanisms underlying stress resilience and vulnerability. Further investigation into the regulatory mechanisms and functional implications of 5αR2 in the brain is warranted to elucidate its precise role in stress adaptation and related neurological processes. Moreover, translational studies in humans are needed to validate the relevance of these findings in clinical settings. Understanding the interplay between 5αR isoenzymes, neurosteroid synthesis, and stress-related disorders could lead to the development of novel pharmacological interventions targeting these pathways.

## METHODS

### Animals

Experiments were performed using Long-Evans (Charles River Laboratories, Raleigh, NC) and Holtzman Sprague Dawley (HSD, Envigo, Indianapolis, IN) male and female rats, weighing 250-350 g, and housed in groups of 3-4 per cage. Animals were kept in a dark-light cycle, with lights off at 6:00 AM and on at 6:00 PM. Unless otherwise stated for specific experimental manipulations, rats had *ad libitum* access to food and water. For all tests, animals were used only once. Separate groups of animals were used for each behavioral paradigm to avoid stress carry-over effects. Experimental manipulations were carried out in the animals’ dark cycle between 10:00 AM and 06:00 PM (with the only exception of the sucrose preference test, which was continuously assessed over several days). All handling and experimental procedures complied with the National Institute of Health guidelines and were approved by the local Institutional Animal Care and Use Committees of the University of Utah (Protocol 19-05005).

### Drugs

Allopregnanolone (AP, Tocris Bio-Techne, Minneapolis, MN) was dissolved in 2.5% DMSO, 2.5% Tween 80, and 0.9% NaCl, and administered IP. The injection dose for AP systemic administrations was 6□mg/kg, while the injection volume was kept at 2 ml/kg.

### RNA interference

Adeno-associated viral (AAV) constructs were obtained from Vector Biolabs (Malvern, PA). All the constructs carried the eGFP reporter under the CMV promoter and the shRNA sequence (against 5αR1, 5αR2, or scrambled) under the U6 promoter. Constructs were packaged into AAV5 capsid, with a titer of 1.5×10^12^ GC/ml for 5αR1, 2.1×10^12^ GC/ml for 5αR2, and 1.2×10^12^ GC/ml for scrambled. A mix of 3 different sequences was used to efficiently downregulated the expression of the target of interest, specifically:

5αR1:

5‘-ACC GCTATGTACAGAGCAGATACTCTCGAGAGTATCTGCTCTGTACATAGCTTTTT-3‘

5‘-ACC GCACCATCAGTGGTACCATGACTCGAGTCATGGTACCACTGATGGTGCTTTTT-3‘

5‘-ACC GGGAAACTGGATACAAGATACCTCGAGGTATCTTGTATCCAGTTTCCCTTTTT-3‘

5αR2:

5‘-ACCG TACTTCCACAGGACATTTATT CTCGAG AATAAATGTCCTGTGGAAGTATTTTT-3‘

5‘-ACCGGTACACAGATGTGCGGTTTACTCGAGTAAACCGCACATCTGTGTACCTTTTT-3‘

5’-ACCGCAGGAGTTGCCTTCCTTTGTCTCGAGACAAAGGAAGGCAACTCCTGCTTTTT-3‘

Long Evans rats were anesthetized with xylazine/ketamine (20/80 mg/kg, IP) and then placed onto a stereotaxic frame (David Kopf Instruments, Tujunga, CA). Blunt ear bars were used to avoid damage to the tympanic membrane. Under aseptic conditions, rats were shaved, and their scalp was retracted. Bilateral craniotomies were made above the target site and a 5μl Hamilton Neuros syringe (Reno, NV) was positioned above bregma. The target locations for brain infusions were: mPFC: AP + 3.0 mm, ML ± 0.5 mm, DV − 3.0 mm from the dura mater; NAc: AP + 1.7 mm, ML ± 0.8 mm, DV = − 7.8 mm from the dura mater. Coordinates were taken from bregma, according to the stereotaxic brain atlas (*38*). The needle was slowly lowered into the injection site, and 1 μl of adeno-associated virus (AAV) was infused into one hemisphere. The syringe was left in place for at least 5 minutes after completion of the infusion to eliminate back-flow, and then slowly withdrawn. This process was repeated on the other side to produce bilateral viral injections. After completing injections into both hemispheres, rats were placed back on a warming pad until the body temperature returned to normal and the recovery of normal movement. Rats were given antibiotic therapy for two days (enrofloxacin, Bayer HealthCare, Shawnee Mission, KS) and were allowed to recover in their home cages (single-housed) with food and water available. Behavioral testing started 14 days after surgery.

### Generation of targeted *Srd5a2* null mutations

HSD rats were maintained in standard housing in an Association for Assessment and Accreditation of Laboratory Animal Care-accredited animal facility. Animal protocols were approved by the Institutional Animal Care and Use Committee at the University of Kansas. Targeted mutations were generated using CRISPR/Cas9-based genome editing strategies. Single guide RNAs (gRNA) specific for the rat *Srd5a2* gene were designed, assembled, and validated by the Genome Engineering and IPSC Center at Washington University in St Louis, MO. Selected gRNAs were targeted to Exon 2 of the *Srd5a2* gene (target sequence: ACTCCTTCAAGCCTACTACCNGG; corresponding to nucleotides 416-435, NM_022711). To obtain viable zygotes four-to five-week-old donor rats were intraperitoneally injected with 30 units of pregnant male serum gonadotropin (PMSG, Sigma-Aldrich, St. Louis, MO), followed by an intraperitoneal injection of 30 units of human chorionic gonadotropin (hCG, Sigma-Aldrich) approximately 46 h later, and immediately mated with stud males. Zygotes were recovered from oviducts the next morning (0.5 days post conception) and then maintained in warm M2 medium supplemented with bovine serum albumin (Millipore) at 37°C, 5% CO_2_ for 2 h. Zygotes were microinjected with a mixture of Cas9 mRNA (30 ng/ml) and *Srd5a2* sgRNAs (20 ng/ml) prepared in TE buffer (pH 7.4) and microinjected into single-cell rat embryos. Microinjections were performed using a Leica microscope and Eppendorf microinjection system. Pseudopregnant females were generated by mating with vasectomized males. On day 0.5 of pseudopregnancy, rats were anesthetized by isoflurane followed by transfer of injected zygotes in oviducts (20–30 zygotes per pseudopregnant rat). Offspring were screened for mutations at specific target sites within the *Srd5a2* gene. For initial screening, genomic DNA was purified from tail-tip biopsies using the Extract-N- Amp™ Tissue PCR Kit (Sigma Aldrich). Potential mutations within the target locus were screened by PCR and accurate boundaries of deletions determined by DNA sequencing (Genewiz Inc, South Plainfield, NJ). PCR primers used for the genotyping: forward primer: 5’-ACCTCTGGCTCGTACCATCT-3’; reverse primer: 5’-TTCACTTTGATCCTGGCCCC-3’. Germline transmission of the mutations was determined in F1 rats by backcrossing founder rats with a wild-type rat. Detection of mutation from F1 rats identical to the mutation from F0 parents was considered as evidence for germline transmission. To validate the efficiency of 5αR2 deletion, rats (n=3/group) were sacrificed, and the prostate tissue was collected.

### Behavioral Manipulations

#### 1) Locomotor activity

Locomotion was evaluated in a square force plate actometer. The apparatus consisted of a white load plate (42 × 42 cm) surrounded on all four sides and covered by a clear Plexiglas box (30 cm tall). Four force transducers placed at the corners of each load plate were sampled 100 times/s, yielding a 0.01 s temporal resolution, a 0.2 g force resolution, and a 2 mm spatial resolution. Custom software directed the timing and data-logging processes via a LabMaster interface (Scientific Solutions Inc., Mentor, OH). Total distance traveled was calculated as the sum of the distances between coordinates of the location of center of force, recorded every 0.5 s over the recording session. Animals were placed in the center of the arena and their behavior was monitored for 15 min. The test was performed in complete darkness to avoid potential anxiety-related confounds.

#### 2) Defensive withdrawal

The defensive withdrawal test was performed as previously described (*13*). Adult rats were placed in a round arena (diameter: 124 cm; height: 56 cm) under bright light (200 lux) and video-recorded for 10 min. A black bucket (height: 20 cm; length: 8 cm; depth: 25 cm) was located inside the arena to provide a hiding place for the animal. At the beginning of the test, the rat was placed inside the bucket, and its behavior was recorded for 10 minutes to evaluate the latency to leave the chamber and the time spent outside the chamber. Behavioral analyses were performed by blinded observers.

#### 3) Olfactory arousal

This test was designed to assess the exploration of novel olfactory stimuli (see Fig. S7A). Rats were individually placed in a vertical plexiglass cylinder (30 cm diameter, 45 cm height) with four scented cotton swabs beneath the lid to encourage rearing towards the scents. Lighting was maintained at 50 lux throughout. The protocol consisted of six trials, each lasting 120 seconds, with a 30- second rest interval between trials. During each interval, the rat was gently removed from the cylinder, the swabs were replaced, and the rat was placed back in the cylinder. In the first five trials, the swabs were soaked with either almond or lemon scent (250 μl, counterbalanced across treatment groups to avoid bias). In the sixth trial, swabs with a novel scent were introduced. Olfactory arousal was assessed by comparing the number of exploratory rears toward the new odor with those from the last trial of the previous scent. To validate the protocol (Fig. 7B), a preliminary study was conducted with rats treated with low-dose reserpine (1 mg/kg/day SC for six days), which is known to alter exploratory drive without affecting locomotor activity (*39*).

#### 4) Social interaction

Social interaction was tested as previously described (*13*). Social interaction was tested against foreign counterparts in a round aluminum chamber (diameter: 124 cm; height: 56 cm). Before the behavioral test, both the experimental and the counterpart animals were acclimated to the aluminum chamber. Social interaction was videotaped and the latency to the first interaction, the duration of total interaction, and sniffing in the genital, mid-sectional, and facial areas were measured by a blinded observer. Environmental lights were maintained at 50 lux.

#### 5) Sucrose Preference Test (SPT)

Rats were kept in isolation for 4 days, and then habituated to consuming a sucrose solution to avoid potential avoidance due to neophobia. They were deprived of water for 15 hours before the test, starting 1 hour before the onset of the dark phase. Each animal was given access to two bottles containing a 2% sucrose solution in tap water, placed on both the right and left sides of the cage, for 3 hours. The following day, rats were again deprived of water for 12 hours, after which they were provided with two bottles: one containing a 2% sucrose solution in tap water, placed on the non-preferred side (i.e., the side from which they had consumed less sucrose the previous day), and the other containing tap water on the opposite side. Sucrose or water consumption was measured every 24 hours for at least 3 consecutive days. To prevent spills, consumption calculations were made by observing the bottle meniscus and estimating the amount consumed using calibration curves that relate the meniscus level (observed at a fixed angle by the same operator) to the weight of the remaining liquid. We previously confirmed that this method provides >99% accuracy in estimating the volume of liquid consumed while avoiding potential spills associated with repeated bottle weighing. Sucrose preference was determined as the ratio of sucrose solution to total liquid consumed by each rat.

#### 6) Forced Swim

Rats were plunged individually into a vertical plexiglass cylinder (diameter: 30 cm; height: 45.7 cm) containing 30 cm of water maintained at 25°C. After 10 min in the cylinder, they were removed and allowed to dry for 15 min in a heated enclosure (32°C) before being returned to their cages. Environmental light was kept at 300 lux. Animals were video recorded, and the latency to immobility (s) and the duration of immobility (s) during the first 5 minutes were measured. When the FST was used as an acute stress paradigm, animals were sacrificed immediately or 30 minutes after the test.

#### 7) Footshock

Footshock stress was delivered through the Potentiated Startle Kit (SR-LAB, San Diego Instruments, San Diego, CA). Adult rats were placed in plexiglass enclosures with shock grids within sound-attenuating startle chambers. After a 1-min acclimation, animals were subjected to nine brief (1 s) electric shocks (0.2-0.5 mA), administered at random intervals of 30-60 s over 5 min. Rats were sacrificed 30 minutes after the footshock.

### RNA qPCR

Tissue from the medial prefrontal cortex, amygdala, hypothalamus, and hippocampus of experimental animals was homogenized in phenol–guanidine isothiocyanate (Trizol) (ThermoFisher Scientific, Waltham, MA). RNA samples were then treated with Turbo DNase (ThermoFisher Scientific) to remove contamination with genomic DNA. Total RNA extracted was evaluated with a NanoDrop 1000 spectrophotometer (ThermoFisher Scientific) to determine its concentration and purity. Three replicates of each RNA sample were measured, averaging concentrations. Only samples with an OD260/280 between 1.8 and 2.0 were used for the following analyses. Electrophoresis with ethidium bromide staining was carried out to evaluate total RNA integrity. RNA samples were stored at −80°C until analysis. For real time RT-PCR, first strand cDNA was generated from 1μg total RNA by reverse transcription using MuLV reverse transcriptase (ThermoFisher Scientific) as previously described (*40*). Absolute quantification of the mRNA of 5αR1 and 5αR2 was performed using real-time RT-PCR on the Techne Quantica™ Real-time PCR system (Burlington, NJ) with SYBR Green PCR Master Mix (Promega, Madison, WI). The standard curves were generated following the procedure described by (*41*). The cRNA was purified with Turbo-DNAse (ThermoFisher Scientific), and its purity and concentration were measured spectrophotometrically. The amount of mRNA was expressed as number of mRNA copies per microgram of total RNA. The standard cRNA was serially diluted to 1×102–1×109□copies/µL. The PCR profile was as follows: denaturation at 94°C for 30 s, annealing at 55°C, and extension at 72°C for 30 s. The number of cycles was always 40. At the end of the amplification phase, melting curve analysis was performed on the products formed to confirm that a single PCR product was detected with the SYBR Green dye. All reactions were run in triplicate, and no cDNA was added to negative reactions.

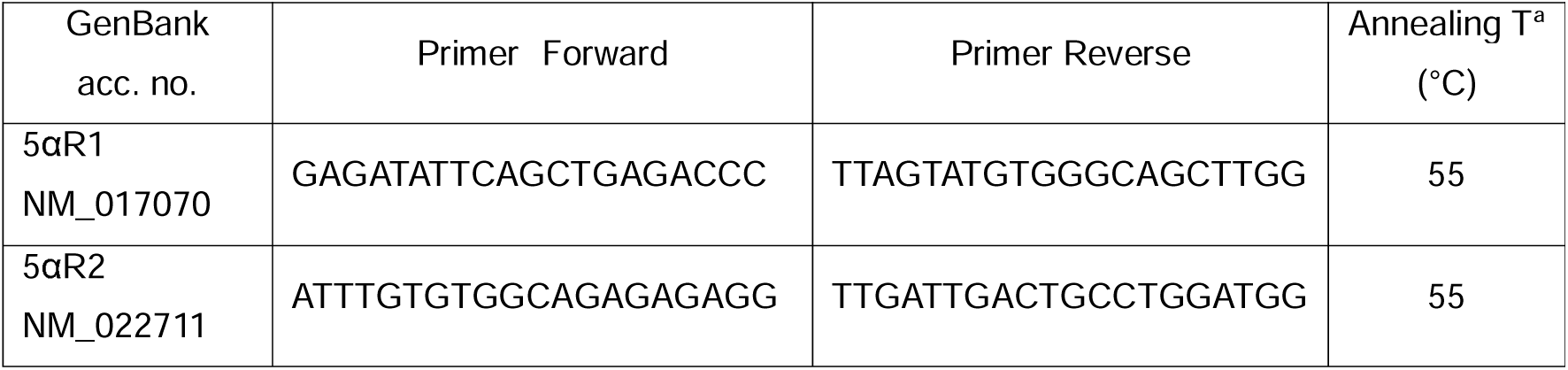

### Immunoblotting

Rats were euthanized by isoflurane and immediately decapitated; brain areas were dissected and flash-frozen for western blot analysis. Tissues were weighted and diluted (100 µg/10 µl) in RIPA buffer [20mM Tris-HCl (pH 7.5), 150 mM NaCl, 1 mM Na2EDTA, 1mM EGTA, 1% NP-40, 1% sodium deoxycholate, 2.5 mM sodium pyrophosphate, 1mM beta-glycerophosphate, 1mM Na_3_VO_4_, 1 µg/ml leupeptin and protease inhibitor cocktail]. Samples containing 15 µg of total proteins were run in duplicate onto 4-15% Criterion™ TGX Stain-free™ precast gels or 12% Criterion™ TGX precast gels (Bio-Rad) and transferred into nitrocellulose membranes (Bio-Rad). Primary antibodies (the complete list is reported in Table 1) were incubated in TBS-T containing 3% (w/v) BSA. Next, blots were washed in TBS- T and then incubated in TBS-T containing secondary antibodies for 90 minutes at room temperature. Chemiluminescence was detected with the ChemiDocTM XRS+ Imaging System (Bio-Rad) using the Clarity Western ECL substrate (Bio-Rad). For further details on the antibodies and dilutions, see Table 1.

**Table 1:**
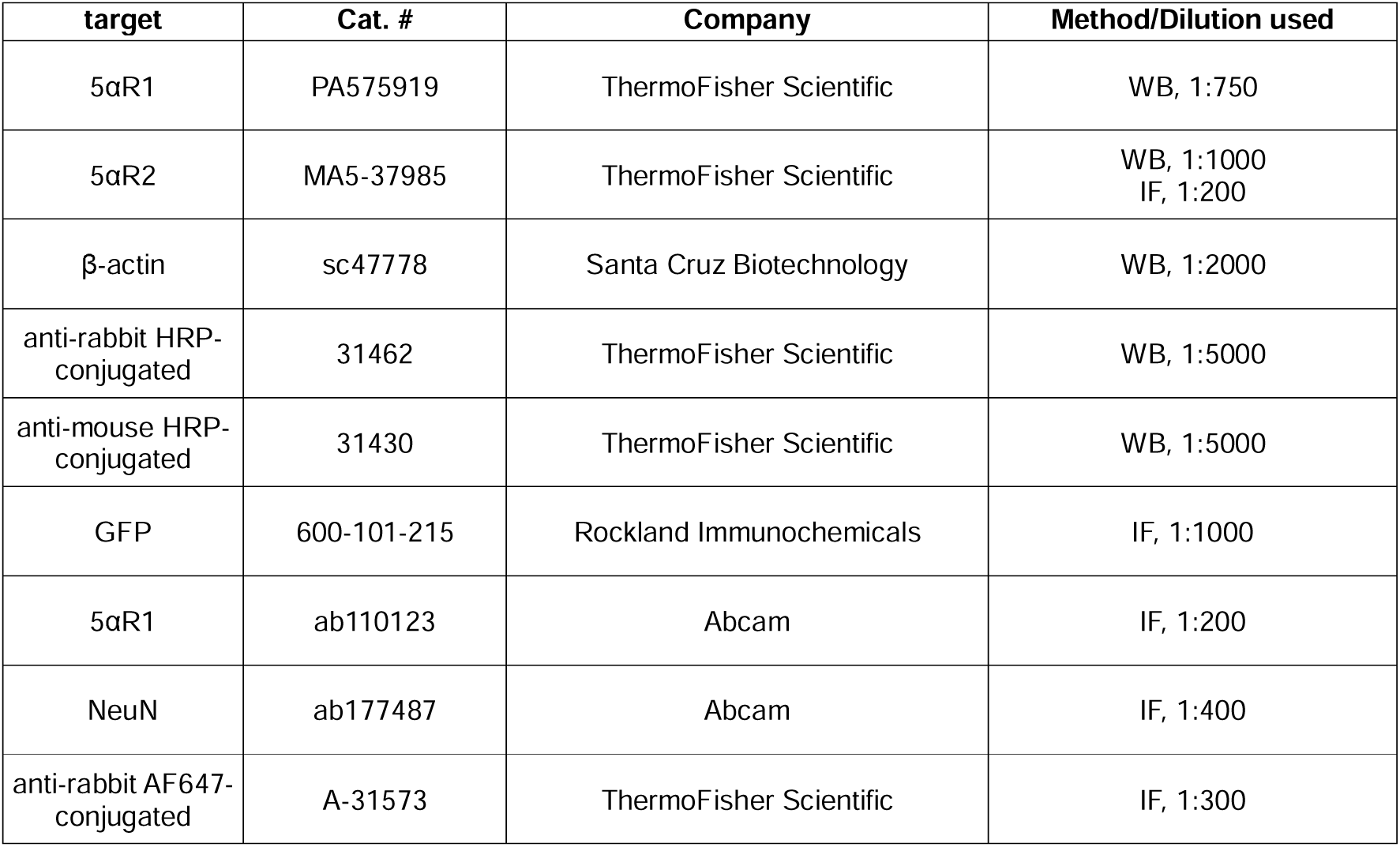
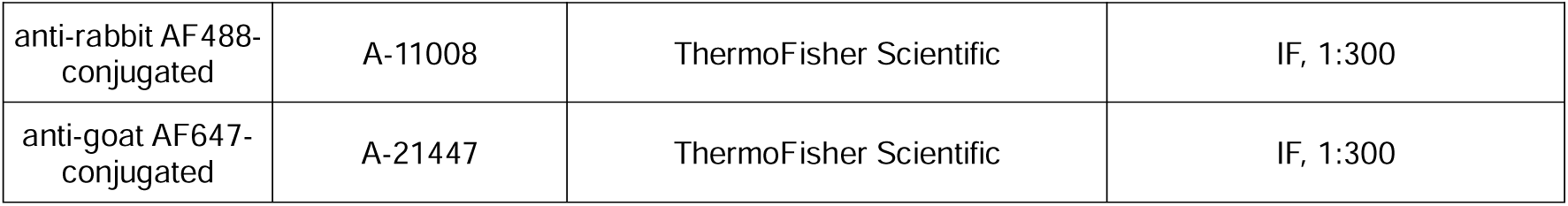
antibodies used in this study.

### Immunofluorescence (IF)

Rats were euthanized with xylazine/ketamine (20/80 mg/kg, IP) and intracardially perfused. Brains were initially fixed in 4% paraformaldehyde (PFA, #J19943, ThermoFisher Scientific) at 4°C for 48 hours, followed by washing in 30% sucrose in PBS. Afterward, the samples were transferred into 70% ethanol and subsequently embedded in formalin. Ten-micron-thick slices were obtained using a microtome and directly mounted onto slides. The slides were deparaffinized through three changes of xylene, then rehydrated with a series of descending ethanol concentrations, concluding with deionized water. Antigen retrieval was performed by submerging the slides in Tris-EDTA Antigen Retrieval Buffer (PR30002, Proteintech, Rosemont, IL) and heating them to 95°C for 20 minutes. After cooling to room temperature, slides were washed twice with TBS (50 mM Tris-HCl, pH 7.4, and 150 mM NaCl), once with TBS-A (TBS + 0.1% Triton X-100), and once with TBS-B (TBS-A + 2% BSA). Primary antibodies (see Table 1 for the complete list) were diluted in TBS-B and incubated overnight at 4°C. The following day, excess primary antibody was removed by two washes with TBS-A and one with TBS-B, and the slides were then incubated for 1 hour at room temperature with secondary antibodies and DAPI diluted in TBS-B (D9542, SigmaAldrich). After incubation, slides were washed three times with TBS and coverslipped using Fluoroshield mounting medium (F6182, SigmaAldrich). For each staining, two negative controls were prepared: one without the primary antibody and one without the secondary antibody, to check for non-specific binding. Images were acquired using a Leica SP8 confocal microscope (Leica, Germany). To quantify 5αR2 levels, two images per experimental animal were captured for each region (prelimbic, PL, and infralimbic, IL). A custom macro in ImageJ (reported below) was then used to measure the mean intensity signal of the target protein and the number of nuclei in each image, which were subsequently used for normalization of the results.

~~~
run("8-bit");
run("Auto Threshold", "method=Shanbhag ignore_black white");
run("Measure");
close;
run("8-bit");
run("Auto Threshold", "method=Default ignore_black white");
run("Analyze Particles…", "size=10-Infinity show=Outlines exclude summarize");
close;
close;
~~~

### Steroid analyses

Extraction, derivatization, and GC-MS analyses of progesterone and AP levels were performed as described in (6). The PFCs of male rats belonging to all the experimental groups were harvested, snap-frozen, and homogenized in distilled water. Supernatants were extracted with ethyl acetate, and after lyophilization, neurosteroids were purified and separated by HPLC (*42*). Tritiated neurosteroids (American Radiolabeled Chemicals, St. Louis, MO) were added to monitor retention time through HPLC (*43*), while deuterated internal standards (CDN Isotopes, Pointe-Claire, QC, and Steraloids, Newport, RI) were added to allow quantification of the compound of interest. Mass fragmentography analysis of derivatized hormones was performed in the standard electron impact mode with a detection limit of ≈ 10 fmol and intra-assay coefficients of variation less than 5%. Neurosteroids were identified based on their GC/MS retention time characteristics; the definitive structural identification of each neurosteroid was provided by its unique mass fragmentation pattern. To calculate the quantity of the neurosteroid of interest in each fraction, the area under the peak of the neurosteroid in the sample was divided by the area under the peak of the respective deuterated internal standard. Only peaks with a signal-to-noise ratio greater or equal to 5:1 were integrated.

### Single Nuclei Preparation and Transcriptomic Analyses

Freshly dissected PFC were immediately flash-frozen using liquid nitrogen and stored at −80°C until further processing for nuclei extraction. Upon processing, specimens were thawed in ice-cold phosphate-buffered saline (PBS, devoid of Ca^2+^ and Mg^2+^) supplemented with 0.54 μM Necrostatin-1 (#J65341, ThermoFisher Scientific), 1 μM HPN-07 (SML2163, Sigma-Aldrich, St. Louis, MO), 0.32 μM sodium hydroxybutyrate (A1161314, ThermoFisher Scientific), 78 nM Q-VD-Oph (SML0063, Sigma-Aldrich), and 0.2 U/ml of RiboLock RNase inhibitor (EO0381, Life Technologies Corporation, Carlsbad, CA). The tissue was then minced and transferred into a 2 ml Dounce homogenizer containing nuclei extraction buffer (130-128-024, Miltenyi Biotec, Bergisch Gladbach, Germany) along with 0.2 U/ml Ribolock RNase inhibitor. After homogenization, samples were centrifuged at 600×g for 5 min, resuspended 2% BSA in PBS, and strained sequentially with 70-μm FLOWMI and 40-μm FLOWMI. Library construction and sample sequencing were performed by the Huntsman Cancer Institute High Throughput Genomics Core. The 10X Genomics Cell Ranger Single Cell pipeline generated alignments and counts using the recommended parameters. Cell ranger’s default R_norvegicus_Mar_2012 dataset was used for alignment. Seurat (4.1.0) was used for quality control and downstream analysis. Cells were filtered out when having less than 750 genes and more than 15% mitochondrial genes. Genes were filtered out including a) mitochondrial genes; b) hemoglobin genes; and c) bias-producing genes, including Gm42418, AY036118, Gm47283, Rpl26, Gstp1, Rpl35a, Erh, Slc25a5, Pgk1, Eno1,Tubb2a, Emc4, Scg5, Ehd2, Espl1, Jarid1d, Pnpla4, Rps4y1, Xist, Tsix, Eif2s3y, Ddx3y, Uty, Kdm5d, Cmss1, AY036118, and Gm47283. Data were processed, including dimension reduction and clustering by SCTransformation (0.3.5) using the Gamma-Poisson generalized linear model (glmGamPoi, 1.4.0) method. Cell subpopulations were identified for clusters using differential gene expression, and all manual annotations were compared to those produced through automated classification using SingleR (1.6.1) and enrichR (3.1, CellMarker Augmented 2021). Pathway and Gene Ontology analyses were performed with enrichR (3.1). Differentially expressed genes (DEGs) were defined as genes that had a 1.5-fold expression over the background with an FDR less than 0.05. To generate the heatmap, differentially expressed genes (DEGs) with a p-value of less than 0.0125 were selected for Gene Ontology (GO) analysis. The resulting GO terms were sorted based on their adjusted p-values, from lowest to highest. The top 20 unique pathways were identified, and their signals were analyzed across all clusters, providing the data for the heatmap visualization. High-dimensional weighted gene co-expression network analysis [hdWGCNA (*44*)] was used to evaluate gene co-expression in single-nuclei sequencing. For this analysis, we subset the data by the Seurat cluster-identified cell types, and then genes were selected if at least 5% of the nuclei in the cell type expressed that gene. The nearest neighbor parameter, k was set to 20 for each subset analysis with at most ten shared cells between metacell groups. For each subgroup, the soft power threshold was identified and the topological overlap matrix was generated. Module eigengenes were identified and the module connectivity was computed. Pathway analysis on the module genes was performed with enrichR.

### Statistical analyses

Statistical analyses were conducted utilizing either one- or multiway ANOVA, as appropriate, with Brown-Forsythe transformation employed when necessary to correct heteroskedasticity. Post-hoc comparisons were executed using Tukey’s test. Statistical significance was determined at the 0.05 threshold. While only the primary outcomes of statistical analyses are delineated in the main text, comprehensive details are provided in the Figure Legends and in the Supplementary Results. Direct comparisons of snRNA-seq data used Wilcoxon rank sum tests for significance testing.

## Author Contributions

Conceptualization, S.S. and M.B.; Methodology, P.S, G.F., J.M.T., E.O., G.P., P.J.M, and M.B; Investigation, R.C., G.B., G.F., E.C., S.S., P.S., L.S.S., J.M.Y., E.O., G.P., and P.J.M.; Writing, R.C., C.B., S.S., and M.B; Funding Acquisition, M.B.; Resources, P.J.M. and M.B.; Supervision, S.S. and M.B.

## Acknowledgments

We thank Michael J. Soares and his group at KUMC for their support in generating the full-body *Srd5a2* knockout rat model. We acknowledge the Huntsman Cancer Institute High-throughput genomics and Bioinformatics Center, as well as the HSC Cell Imaging Core, for enabling sh-RNA sequencing and IF microscopy analyses. We thank Laura Mosher, Sean Godar, Hunter Strathman, Claudia Collu, Karen Odeh, Marco Orrù, Eva Vigato, Stefanos Loizou, and Easton Van Luik for their technical assistance with animal testing. We thank Anton Classen for his invaluable support in image acquisition and analysis.

## Data and Material Availability

All data needed to evaluate the conclusions in the paper are present in the paper and/or the Supplementary Materials. The data generated for sn-RNA seq are available on Gene Expression Omnibus, GSE267559.

## References

1. G. Russell, S. Lightman, The human stress response. Nat. Rev. Endocrinol. 15, 525–534 (2019).

2. B. S. McEwen, J. Gray, C. Nasca, Recognizing Resilience: Learning from the Effects of Stress on the Brain. Neurobiol. Stress 1, 1–11 (2015).

3. B. S. McEwen, R. M. Sapolsky, Stress and cognitive function. Curr. Opin. Neurobiol. 5, 205–216 (1995).

4. B. G. Gunn, L. Cunningham, S. G. Mitchell, J. D. Swinny, J. J. Lambert, D. Belelli, GABAA receptor-acting neurosteroids: a role in the development and regulation of the stress response. Front. Neuroendocrinol. 36, 28–48 (2015).

5. M. L. Barbaccia, G. Roscetti, M. Trabucchi, M. C. Mostallino, A. Concas, R. H. Purdy, G. Biggio, Time-dependent changes in rat brain neuroactive steroid concentrations and GABAA receptor function after acute stress. Neuroendocrinology 63, 166–172 (1996).

6. R. Cadeddu, L. J. Mosher, P. Nordkild, N. Gaikwad, G. M. Ratto, S. Scheggi, M. Bortolato, Acute stress impairs sensorimotor gating via the neurosteroid allopregnanolone in the prefrontal cortex. Neurobiol. Stress 21, 100489 (2022).

7. R. H. Purdy, A. L. Morrow, P. H. Moore, S. M. Paul, Stress-induced elevations of gamma-aminobutyric acid type A receptor-active steroids in the rat brain. Proc. Natl. Acad. Sci. U. S. A. 88, 4553–4557 (1991).

8. S. F. Maier, L. R. Watkins, Role of the medial prefrontal cortex in coping and resilience. Brain Res. 1355, 52–60 (2010).

9. D. H. Legesse, C. Fan, J. Teng, Y. Zhuang, R. J. Howard, C. M. Noviello, E. Lindahl, R. E. Hibbs, Structural insights into opposing actions of neurosteroids on GABAA receptors. Nat. Commun. 14, 5091 (2023).

10. S. Diviccaro, L. Cioffi, E. Falvo, S. Giatti, R. C. Melcangi, Allopregnanolone: An overview on its synthesis and effects. J. Neuroendocrinol. 34, e12996 (2022).

11. S. Paba, R. Frau, S. C. Godar, P. Devoto, F. Marrosu, M. Bortolato, Steroid 5α- reductase as a novel therapeutic target for schizophrenia and other neuropsychiatric disorders. Curr. Pharm. Des. 17, 151–167 (2011).

12. A. Concas, M. C. Mostallino, P. Porcu, P. Follesa, M. L. Barbaccia, M. Trabucchi, R. H. Purdy, P. Grisenti, G. Biggio, Role of brain allopregnanolone in the plasticity of gamma-aminobutyric acid type A receptor in rat brain during pregnancy and after delivery. Proc. Natl. Acad. Sci. U. S. A. 95, 13284–13289 (1998).

13. S. C. Godar, R. Cadeddu, G. Floris, L. J. Mosher, Z. Mi, D. P. Jarmolowicz, S. Scheggi, A. A. Walf, C. J. Koonce, C. A. Frye, N. A. Muma, M. Bortolato, The Steroidogenesis Inhibitor Finasteride Reduces the Response to Both Stressful and Rewarding Stimuli. Biomolecules 9, 749 (2019).

14. D.W. Russell, J.D. Wilson, Steroid 5α-reductase: two genes/two enzymes. Annu. Rev. Biochem. 63, 25–61 (1994).

15. M. P. Castelli, A. Casti, A. Casu, R. Frau, M. Bortolato, S. Spiga, M. G. Ennas, Regional distribution of 5α-reductase type 2 in the adult rat brain: an immunohistochemical analysis. Psychoneuroendocrinology 38, 281–293 (2013).

16. J. M. Torres, E. Ortega, Differential regulation of steroid 5alpha-reductase isozymes expression by androgens in the adult rat brain. FASEB J. Off. Publ. Fed. Am. Soc. Exp. Biol. 17, 1428–1433 (2003).

17. J. M. Torres, E. Ortega, Steroid 5alpha-reductase isozymes in the adult female rat brain: central role of dihydrotestosterone. J. Mol. Endocrinol. 36, 239–245 (2006).

18. E. J. Nestler, W. A. Carlezon, The mesolimbic dopamine reward circuit in depression. Biol. Psychiatry 59, 1151–1159 (2006).

19. S. J. Russo, E. J. Nestler, The brain reward circuitry in mood disorders. Nat. Rev. Neurosci. 14, 609–625 (2013)

20. A. Poletti, A. Coscarella, P. Negri-Cesi, A. Colciago, F. Celotti, L. Martini. 5 alpha-reductase isozymes in the central nervous system. Steroids 63, 246–251 (1998).

21. J. Imperato-McGinley, L. Guerrero, T. Gautier, R.E. Peterson. Steroid 5alpha-reductase deficiency in man: an inherited form of male pseudohermaphroditism. Science. 186, 1213–1215 (1974).

22. P. Sánchez, J.M. Torres, P. Gavete, E. Ortega. Effects of swim stress on mRNA and protein levels of steroid 5alpha-reductase isozymes in prefrontal cortex of adult male rats. Neurochem Int. 52, 426–431 (2008).

23. P. Sánchez, J. M. Torres, A. Olmo, F. O’Valle, E. Ortega, Effects of environmental stress on mRNA and protein expression levels of steroid 5alpha-Reductase isozymes in adult rat brain. Horm. Behav. 56, 348–353 (2009).

24. F.B. Kraemer, W.J. Shen, S. Azhar. SNAREs and cholesterol movement for steroidogenesis. Mol Cell Endocrinol. 441,17–21 (2017).

25. M. G. Pisu, A. Garau, G. Boero, F. Biggio, V. Pibiri, R. Dore, V. Locci, E. Paci, P. Porcu, M. Serra, Sex differences in the outcome of juvenile social isolation on HPA axis function in rats. Neuroscience 320, 172–182 (2016).

26. R. Frau, G. Pillolla, V. Bini, S. Tambaro, P. Devoto, M. Bortolato Inhibition of 5α- reductase attenuates behavioral effects of D1-, but not D2-like receptor agonists in C57BL/6 mice. Psychoneuroendocrinology. 38, 2013 Apr;(4):542–551 (2013).

27. R. Frau, L.J. Mosher, V. Bini, G. Pillolla, R. Pes, P. Saba, S. Fanni, P. Devoto, M. Bortolato. The neurosteroidogenic enzyme 5α-reductase modulates the role of D1 dopamine receptors in rat sensorimotor gating. Psychoneuroendocrinology. 63, 59–67 (2016).

28. R. Shinohara, M. Taniguchi, A.T. Ehrlich, K. Yokogawa, Y. Deguchi, Y. Cherasse, M. Lazarus, Y. Urade, A. Ogawa, S. Kitaoka, A. Sawa, S. Narumiya, T. Furuyashiki. Dopamine D1 receptor subtype mediates acute stress-induced dendritic growth in excitatory neurons of the medial prefrontal cortex and contributes to suppression of stress susceptibility in mice. Mol Psychiatry. 23, 1717–1730 (2018).

29. C.J. McNulty, I.P. Fallon, J. Amat, R.J. Sanchez, N.R. Leslie, D.H. Root, S.F. Maier, M.V. Baratta. Elevated prefrontal dopamine interferes with the stress-buffering properties of behavioral control in female rats. Neuropsychopharmacology. 48, 498–507 (2023).

30. D.D. Markov, Sucrose Preference Test as a Measure of Anhedonic Behavior in a Chronic Unpredictable Mild Stress Model of Depression: Outstanding Issues. Brain Sci. 12, 1287 (2022).

31. C. Belzung, M. Lemoine, Criteria of validity for animal models of psychiatric disorders: focus on anxiety disorders and depression. Biol Mood Anxiety Disord. 1, 9 (2011).

32. P. Willner, Validity, reliability and utility of the chronic mild stress model of depression: a 10-year review and evaluation. Psychopharmacology (Berl).134, 319–329 (1997).

33. M. Becker, A. Pinhasov, A. Ornoy. Animal Models of Depression: What Can They Teach Us about the Human Disease?. Diagnostics (Basel). 11, 123 (2021).

34. C. Bielajew, A.T. Konkle, A.C. Kentner, et al. Strain and gender specific effects in the forced swim test: effects of previous stress exposure. Stress. 6, 269–280 (2003).

35. C. Schüle, C. Nothdurfter, R. Rupprecht, The role of allopregnanolone in depression and anxiety. Prog. Neurobiol. 113, 79–87 (2014).

36. G. Pinna, Allopregnanolone, the Neuromodulator Turned Therapeutic Agent: Thank You, Next? Front. Endocrinol. 11, 236 (2020).

37. M. Bortolato, P. Devoto, P. Roncada, R. Frau, G. Flore, P. Saba, G. Pistritto, A. Soggiu, S. Pisanu, A. Zappala, M. S. Ristaldi, M. Tattoli, V. Cuomo, F. Marrosu, M. L. Barbaccia, Isolation rearing-induced reduction of brain 5α-reductase expression: relevance to dopaminergic impairments. Neuropharmacology 60, 1301–1308 (2011).

38. G. Paxinos, C. Watson, The Rat Brain in Stereotaxic Coordinates (Academic Pr, Sydney, 1982).

39. R. Pes, S.C. Godar, A.T. Fox, L.M. Burgeno, H.J. Strathman, D.P. Jarmolowicz, P. Devoto, B. Levant, P.E. Phillips, S.C. Fowler, M. Bortolato. Pramipexole enhances disadvantageous decision-making: Lack of relation to changes in phasic dopamine release. Neuropharmacology 114, 77–87 (2017).

40. P. Sánchez, B. Castro, S. Martínez-Rodríguez, R. Ríos-Pelegrina, R. G. Del Moral, J. M. Torres, E. Ortega, Impact of chronic exposure of rats to bisphenol A from perinatal period to adulthood on intraprostatic levels of 5α-reductase isozymes, aromatase, and genes implicated in prostate cancer development. Environ. Res. 212, 113142 (2022).

41. S. Fronhoffs, G. Totzke, S. Stier, N. Wernert, M. Rothe, T. Brüning, B. Koch, A. Sachinidis, H. Vetter, Y. Ko, A method for the rapid construction of cRNA standard curves in quantitative real-time reverse transcription polymerase chain reaction. Mol. Cell. Probes 16, 99–110 (2002).

42. D. P. Uzunov, T. B. Cooper, E. Costa, A. Guidotti, Fluoxetine-elicited changes in brain neurosteroid content measured by negative ion mass fragmentography. Proc. Natl. Acad. Sci. U. S. A. 93, 12599–12604 (1996).

43. G. Pinna, E. Dong, K. Matsumoto, E. Costa, A. Guidotti, In socially isolated mice, the reversal of brain allopregnanolone down-regulation mediates the anti-aggressive action of fluoxetine. Proc. Natl. Acad. Sci. U. S. A. 100, 2035–2040 (2003).

44. S. Morabito, F. Reese, N. Rahimzadeh, E. Miyoshi, V. Swarup. hdWGCNA identifies co-expression networks in high-dimensional transcriptomics data. Cell Rep Methods. 3, 100498 (2023).

