## Supplementary Materials for "Prefrontal 5α-reductase 2 mediates male-specific acute stress response"

Science Advances

AAAS

Cadeddu *et al.*

**This PDF file includes:**

Supplementary Results

Figs. S1 to S31

**Other Supplementary Materials for this manuscript include the following:**

Data S1 to S28

**Supplementary Results**

We first investigated the metabolic pathway leading to allopregnanolone (AP) synthesis (Fig. S1A) in response to acute stress, by measuring the levels of progesterone, dihydroprogesterone (DHP), and AP levels in the medial prefrontal cortex (mPFC) of male rats exposed to either forced swim test or footshock, as compared with unstressed controls. The goal of these initial studies was to confirm that both stressors elevated AP concentrations under our experimental settings. At 30 minutes post-forced swim test, we observed a marginal increase in progesterone concentrations in the PFC [Fig. S1B; one-way ANOVA: $F(1,17)=4.30$, $p=0.06$, partial $\eta^2=0.20$], accompanied by a significant increase in DHP [Fig. S1C; one-way ANOVA: $F(1,17)=21.40$, $p<0.001$, partial $\eta^2=0.56$] and AP levels [Fig. S1E; one-way ANOVA: $F(1,17)=6.80$, $p=0.02$, partial $\eta^2=0.29$] in the same region. Notably, the DHP/progesterone ratio was significantly elevated [Fig. S1D; one-way ANOVA: $F(1,17)=15.62$, $p=0.001$, partial $\eta^2=0.48$], whereas the AP/progesterone ratio showed no significant difference between stressed and control animals (Fig. S1F). Similar effects were observed 30 minutes post-footshock, with marked increases in all steroid levels (Figs. S1G-K) {Progesterone: [one-way ANOVA: $F(1,14)=8.68$, $p=0.01$, partial $\eta^2=0.38$]; DHP: [one-way ANOVA with Brown-Forsythe correction: $F(1,10.14)=34.87$, $p<0.001$, partial $\eta^2=0.71$]; AP [one-way ANOVA with Brown-Forsythe correction: $F(1,7.97)=13.21$, $p=0.007$, partial $\eta^2=0.49$]} and a significantly elevated DHP/progesterone ratio (Fig. S1I) [one-way ANOVA with Brown-Forsythe correction: $F(1,9.25)=8.02$, $p=0.02$, partial $\eta^2=0.36$]. Again, the AP/progesterone ratio was not increased significantly (Fig. S1K). These findings suggest that both stressors enhance progesterone synthesis and its subsequent conversion to DHP, catalyzed by 5αR.

Building on these findings, we measured the effects of the forced-swim stressor on the mRNA expression levels of the two main 5αR isoforms, 5αR1 and 5αR2, in the mPFC and other brain regions implicated in the stress response. Male rats were exposed to the forced swim and sacrificed immediately after the task (FS 0) or 30 minutes later (FS 30). Control rats were kept in their home cage (non-stressed, NS). 5αR1 and 5αR2 mRNA levels were evaluated by qRT-PCR in extracts from mPFC, hypothalamus (HYP), amygdala (AMY), and hippocampus (HIP). Stress exposure did not alter 5αR1 mRNA levels compared to NS male rats [Fig. S2A, one-way ANOVAs of mPFC, $F(2,17)=1.04$, $p=0.37$, partial $\eta^2=0.11$; AMY, $F(2,18)=0.27$, $p=0.77$, partial $\eta^2=0.03$; HYP, $F(2,16)=0.11$, $p=0.90$, partial $\eta^2=0.01$; HIP, $F(2,16)=0.17$, $p=0.85$, partial $\eta^2=0.02$]. Stress exposure induced a significant upregulation of 5αR2 mRNAs in mPFC [Fig. S2B, one-way ANOVA, $F(2,18)=11.13$, $p<0.001$, partial $\eta^2=0.55$] and HIP (one-way ANOVA, $F(2,16)=23.41$, $p<0.001$, partial $\eta^2=0.75$). No differences were observed in AMY and HYP (one-way ANOVAs, AMY, $F(2,17)=0.05$, $p=0.95$, partial $\eta^2=0.01$; HYP, $F(2,16)=0.20$, $p=0.83$, partial $\eta^2=0.02$).

To better delineate the effects of acute stress on the protein levels of 5αR1 and 5αR2, western blot analyses across the mPFC, nucleus accumbens (NAc), HYP, AMY, and HIP were performed (representative images are reported in Fig. S3). Male and female rats were exposed to forced swim or footshock, two procedures that trigger consistent and reproducible behavioral, physiological, and neurobiological responses.

Analyses of protein levels in brain extracts from male rats revealed that 30 min after forced swim exposure (Fig. 1A), 5αR1 levels remained unchanged in the mPFC [one-way ANOVA with Brown-Forsythe correction: $F(1,5.941)=0.69$, $p=0.44$, partial $\eta^2=0.07$], while being significantly upregulated by stress in the NAc [one-way ANOVA with Brown-Forsythe correction, $F(1,5.258)=20.71$, $p=0.005$, partial $\eta^2=0.67$] and HYP [one-way ANOVA with Brown-Forsythe correction, $F(1,5.114)=16.77$, $p=0.009$, partial $\eta^2=0.63$]. No differences were observed in the AMY [one-way ANOVA with Brown-Forsythe correction, $F(1,5.219)=3.67$, $p=0.11$, partial $\eta^2=0.27$] or HIP [one-way ANOVA with Brown-Forsythe correction, $F(1,6.678)=2.40$, $p=0.17$, partial $\eta^2=0.19$].

When footshock was used as the stressor (Fig. 1B), 5αR1 was significantly downregulated in the NAc [one-way ANOVA, $F(1,8)=33.27$, $p<0.001$, partial $\eta^2=0.81$] and HIP [one-way ANOVA, $F(1,8)=7.61$, $p=0.025$, partial $\eta^2=0.49$] of male rats. No changes were detected in the mPFC [one-way ANOVA, $F(1,8)=2.18$, $p=0.18$, partial $\eta^2=0.21$], HYP [one-way ANOVA, $F(1,8)=1.00$, $p=0.35$, partial $\eta^2=0.11$], or AMY [one-way ANOVA, $F(1,8)=0.02$, $p=0.90$, partial $\eta^2=0.002$]. Representative Western blot images are shown in Fig. S3.

Analyses of 5αR2 protein levels in male rats exposed to forced swim (Fig. 1C) indicated an upregulation in the mPFC [one-way ANOVA with Brown-Forsythe correction, $F(1,5.070)=5.63$, $p=0.04$, partial $\eta^2=0.36$] and HYP [one-way ANOVA with Brown-Forsythe correction, $F(1,5.206)=23.53$, $p=0.004$, partial $\eta^2=0.70$], while AMY showed a downregulation [one-way ANOVA, $F(1,10)=6.44$, $p=0.03$, partial $\eta^2=0.39$]. No significant changes were observed in the NAc [one-way ANOVA with Brown-Forsythe correction, $F(1,5.830)=4.33$, $p=0.08$, partial $\eta^2=0.30$] or HIP [one-way ANOVA, $F(1,10)=4.12$, $p=0.07$, partial $\eta^2=0.29$].

Exposure to footshock stress in male rats (Fig. 1D) led to an upregulation of 5αR2 in the mPFC [one-way ANOVA with Brown-Forsythe correction, $F(1,4.123)=5.47$, $p=0.048$, partial $\eta^2=0.41$]. However, 5αR2 protein levels in the NAc [one-way ANOVA with Brown-Forsythe correction, $F(1,5.628)=1.34$, $p=0.29$, partial $\eta^2=0.14$], HYP [one-way ANOVA, $F(1,8)=3.03$, $p=0.12$, partial $\eta^2=0.28$], AMY [one-way ANOVA, $F(1,8)=0.09$, $p=0.78$, partial $\eta^2=0.01$], and HIP [one-way ANOVA with Brown-Forsythe correction, $F(1,4.504)=2.20$, $p=0.21$, partial $\eta^2=0.22$] remained comparable to non-stressed controls.

Neither forced swim nor footshock stress significantly affected 5αR1 or 5αR2 protein levels in female rats (Fig. 1E-H). For 5αR1 in the forced swim paradigm (Fig. 1E), one-way ANOVA analysis revealed no significant differences in the mPFC [one-way ANOVA with Brown-Forsythe correction, $F(1,5.021)=1.45$, $p=0.28$, partial $\eta^2=0.13$], NAc [one-way ANOVA with Brown-Forsythe correction, $F(1,5.300)=3.36$, $p=0.12$, partial $\eta^2=0.25$], HYP [one-way ANOVA, $F(1,10)=1.34$, $p=0.28$, partial $\eta^2=0.12$], AMY (one-way ANOVA, $F(1,10)=0.94$, $p=0.36$, partial $\eta^2=0.09$], or HIP [one-way ANOVA with Brown-Forsythe correction, $F(1,5.448)=0.39$, $p=0.56$, partial $\eta^2=0.04$]. Similarly, footshock stress did not alter 5αR1 protein levels in female rats (Fig. 1F) in the mPFC (one-way ANOVA, $F(1,8)=0.53$, $p=0.49$, partial $\eta^2=0.06$], NAc [one-way ANOVA with Brown-Forsythe correction, $F(1,4.818)=0.08$, $p=0.79$, partial $\eta^2=0.01$], HYP [one-way ANOVA, $F(1,8)=7.53 \times 10^{-4}$, $p=0.98$, partial $\eta^2=9.42 \times 10^{-5}$], AMY [one-way ANOVA, $F(1,8)=0.73$, $p=0.42$, partial $\eta^2=0.08$], or HIP [one-way ANOVA, $F(1,8)=0.60$, $p=0.46$, partial $\eta^2=0.07$].

For 5αR2 in the forced swim paradigm (Fig. 1G), no significant differences were found in female rats in the mPFC [one-way ANOVA, $F(1,10)=3.25$, $p=0.10$, partial $\eta^2=0.25$], NAc [one-way ANOVA with Brown-Forsythe correction, $F(1,5.124)=1.14$, $p=0.33$, partial $\eta^2=0.10$], HYP [one-way ANOVA, $F(1,10)=1.66$, $p=0.23$, partial $\eta^2=0.14$], AMY [one-way ANOVA, $F(1,10)=0.28$, $p=0.61$, partial $\eta^2=0.03$], or HIP [one-way ANOVA with Brown-Forsythe correction, $F(1,5.091)=0.72$, $p=0.43$, partial $\eta^2=0.07$]. Similarly, 5αR2 levels remained unchanged in all brain areas of female rats subjected to footshock stress [Fig. 1H], as shown by analyses of the mPFC [one-way ANOVA, $F(1,8)=0.12$, $p=0.74$, partial $\eta^2=0.01$], NAc [one-way ANOVA with Brown-Forsythe correction, $F(1,4.54)=0.24$, $p=0.64$, partial $\eta^2=0.03$], HYP [one-way ANOVA, $F(1,8)=0.06$, $p=0.81$, partial $\eta^2=0.01$], AMY [one-way ANOVA with Brown-Forsythe correction, $F(1,4.83)=0.38$, $p=0.57$, partial $\eta^2=0.05$], and HIP [one-way ANOVA, $F(1,8)=0.12$, $p=0.74$, partial $\eta^2=0.02$].

We then performed immunofluorescence analyses to detail the anatomical localization of the upregulated 5αR2 in the mPFC of male rats following forced swim stress. As shown in Fig. S4, the distribution of 5αR2 is strikingly divergent from that of 5αR1 in this region. The distribution of 5αR1 is more uniform across all areas and layers of the mPFC, and includes the somata of neurons as well as glial cells, as indicated by NeuN immunofluorescence. In contrast, 5αR2 was predominantly expressed in the somata and neurites of neurons, with more abundant distribution in the prelimbic and infralimbic cortex. As shown in Fig. S5, the stress-induced upregulation of 5αR2 was observed in both superficial (II-III) and deep layers (V-VI) of both the prelimbic and infralimbic cortex [Fig. S5B, main effect of area: $F(1,10)=9.62$, $p=0.01$, partial $\eta^2=0.49$; main effect of stress: $F(1,10)=13.22$, $p=0.005$, partial $\eta^2=0.57$].

We next examined the functional role of 5αRs in behavioral regulation. To this end, we injected constructs harboring scrambled RNA, 5αR1- or 5αR2-targeting shRNA into the PFC or NAc of male and female rats. Successful targeting of the PFC was confirmed by representative images of AAV5 spreading (Fig. S6A, visualized as GFP staining) and successful downregulation of 5αR1 or 5αR2 expression was quantified by Western blot [Fig. S6B-C, one-way ANOVA, $F(1,4)=18.01$, $p=0.013$, partial $\eta^2=0.82$ for 5αR1; Fig. S6D-E, one-way ANOVA, $F(1,4)=8.13$, $p=0.046$, partial $\eta^2=0.67$ for 5αR2]. Similar reductions were observed in the NAc (Fig. S6F, GFP staining), with quantification showing significant decreases [Fig. S6G-H, one-way ANOVA,

$F(1,4)=9.58$, $p=0.04$, partial $\eta^2=0.71$ for 5αR1; Fig. S6I-J, $F(1,4)=15.07$, $p=0.02$, partial $\eta^2=0.79$ for 5αR2].

Behavioral analyses conducted 14 days post-surgery indicated that knockdown (KD) of 5αR1 in the PFC increased locomotor activity, as measured by distance covered over 15 minutes in the open field arena [Fig. 2A, one-way ANOVA with Brown-Forsythe correction, $F(1,9.548)=6.72$, $p=0.03$, partial $\eta^2=0.30$]. No differences were noted in the forced swim test [Fig. 2B, one-way ANOVA, $F(1,18)=0.14$, $p=0.71$, partial $\eta^2=0.01$] or in the defensive withdrawal paradigm [Fig. 2C, one-way ANOVA with Brown-Forsythe correction, $F(1,12.24)=0.87$, $p=0.40$, partial $\eta^2=0.05$].

We then tested the rats in the olfactory arousal test, a novel paradigm designed to assess exploration toward olfactory stimuli (Fig. S7A; see Methods for details). This test was validated using a reserpine regimen (1 mg/kg/day, SC, for six days), which is known to reduce novelty exploration without affecting locomotor activity (*39*). Analyses were conducted using a two-way repeated-measures ANOVA, which revealed a significant session × treatment interaction [$F(1,9)=14.59$, $p=0.004$, partial $\eta^2=0.62$]. Post-hoc analyses showed significant differences between the number of rears induced by the old scent compared to the new scent in vehicle-treated animals ($p=0.001$) and between the number of rears induced by the new scent in reserpine-treated animals compared to the vehicle-treated animals ($p=0.006$).

New odor exposure elicited similar arousal between animals injected with 5αR1-shRNA and scrambled-shRNA [Fig. 2D, repeated measures two-way ANOVA, main effect of odor, $F(1,14)=16.75$, $p=0.001$, partial $\eta^2=0.55$; main effect of condition, $F(1,14)=0.83$, $p=0.38$, partial $\eta^2=0.06$; interaction, $F(1,14)=0.06$, $p=0.81$, partial $\eta^2=0.004$]. 5αR1 KD rats interacted significantly less with foreign counterparts than control animals [Fig. 2E, one-way ANOVA, $F(1,18)=82.224$, $p<0.001$, partial $\eta 2=0.820$]. No differences were observed in the sucrose preference test [Fig. 2F, repeated measures two-way ANOVA, main effect of time, $F(7,126)=1.10$, $p=0.37$, partial $\eta^2=0.06$; main effect of condition, $F(1,18)=1.53$, $p=0.23$, partial $\eta^2=0.08$; interaction, $F(7,126)=1.69$, $p=0.12$, partial $\eta^2=0.09$]. KD of 5αR1 in the NAc did not impact open-field activity [Fig. 2G, one-way ANOVA, $F(1,18)=0.07$, $p=0.79$, partial $\eta^2=0.004$], forced swim performance [Fig. 2H, $F(1,18)=2.01$, $p=0.17$, partial $\eta^2=0.10$], or defensive withdrawal behavior [Fig. 2I, $F(1,18)=0.87$, $p=0.36$, partial $\eta^2=0.05$]. Responses to new odors were also similar between groups [Fig. 2J, repeated measures two-way ANOVA, main effect of odor, $F(1,15)=9.64$, $p=0.007$, partial $\eta^2=0.39$; main effect of condition, $F(1,15)=0.24$, $p=0.63$, partial $\eta^2=0.02$; interaction, $F(1,15)=0.09$, $p=0.77$, partial $\eta^2=0.01$]. Additionally, 5αR1 KD rats displayed significantly lower social interaction with foreign counterparts [Fig. 2K, one-way ANOVA with Brown-Forsythe correction, $F(1,13.071)=93.96$, $p<0.001$, partial $\eta^2=0.84$]. No differences were found in the sucrose preference test [Fig. 2L, repeated measures two-way ANOVA, main effect of time, $F(7,126)=1.83$, $p=0.09$, partial $\eta^2=0.09$; main effect of condition, $F(1,18)=0.86$, $p=0.37$, partial $\eta^2=0.05$; interaction, $F(7,126)=0.18$, $p=0.99$, partial $\eta^2=0.01$].

The same behavioral assessments were conducted following KD of 5αR2. Male rats injected with 5αR2-targeting shRNA into the PFC showed no changes in open-field activity [Fig. 2M, one-way ANOVA with Brown-Forsythe correction, $F(1,10.983)=0.55$, $p=0.47$, partial $\eta^2=0.03$]; however, 5αR2 KD rats exhibited significantly increased floating time in the forced swim test [Fig. 2N, one-way ANOVA, $F(1,18)=9.52$, $p=0.006$, partial $\eta^2=0.35$]. Additionally, these rats spent significantly less time outside the chamber in the defensive withdrawal paradigm [Fig. 2O, $F(1,17)=5.45$, $p=0.032$, partial $\eta^2=0.24$]. Reduced arousal in response to olfactory cues was noted in 5αR2 KD animals [Fig. 2P, repeated measures two-way ANOVA, main effect of odor, $F(1,12)=24.16$, $p=0.0004$, partial $\eta^2=0.67$; main effect of condition, $F(1,12)=2.46$, $p=0.14$, partial $\eta^2=0.17$; interaction, $F(1,12)=11.03$, $p=0.006$, partial $\eta^2=0.48$]. Furthermore, KD rats exhibited less social interaction compared to scrambled controls [Fig. 2Q, one-way ANOVA, $F(1,18)=4.88$, $p=0.040$, partial $\eta^2=0.21$]. 5αR2 KD rats also did not show a preference for sucrose over time [Fig. 2R, repeated- measure two-way ANOVA: main effect of condition, $F(1,18)=6.37$, $p=0.018$, partial $\eta^2=0.27$; main effect of time, $F(7,126)=0.15$, $p=0.99$, partial $\eta^2=0.01$; interaction, $F(7,126)=4.12$, $p<0.001$, partial $\eta^2=0.19$]. KD of 5αR2 in the NAc did not result in any significant differences in open-field locomotion [Fig. 2S, $F(1,14)=0.44$, $p=0.52$, partial $\eta^2=0.03$], forced swim test performance [Fig. 2T, $F(1,18)=1.18$, $p=0.29$, partial $\eta^2=0.06$], or defensive withdrawal behavior [Fig. 2U, $F(1,18)=0.02$, $p=0.89$, partial $\eta^2=0.001$]. Exposure to a new odor led to similar arousal between groups [Fig. 2V, repeated measures two-way ANOVA, main effect of odor, $F(1,9)=101.80$, $p<0.001$, partial $\eta^2=0.92$; main effect of condition, $F(1,9)=0.22$, $p=0.65$, partial $\eta^2=0.02$; interaction, $F(1,9)=0.67$, $p=0.43$, partial $\eta^2=0.07$]. 5αR2 KD rats interacted significantly less with a foreign conspecific [Fig. 2W, one-way ANOVA, $F(1,18)=16.38$, $p<0.001$, partial $\eta^2=0.48$]. The sucrose preference test revealed no preference on days 5 and 6 [Fig. 2X, repeated measures two-way ANOVA, main effect of condition, $F(1,18)=5.36$, $p=0.033$, partial $\eta^2=0.23$; main effect of time, $F(7,126)=0.66$, $p=0.71$, partial $\eta^2=0.04$; interaction, $F(7,126)=2.64$, $p=0.014$, partial $\eta^2=0.13$].

To explore potential sex differences, female rats were also stereotaxically injected in both areas, and their main behavioral phenotypes were characterized 14 days post-surgery. KD of 5αR1 in the PFC led to an increase in immobility time in the forced swim test [Fig. 3A, one-way ANOVA, $F(1,14)=5.24$, $p=0.038$, partial $\eta^2=0.27$] without affecting defensive withdrawal behavior [Fig. 3B, $F(1,14)=0.94$, $p=0.35$, partial $\eta^2=0.06$] or sucrose preference [Fig. 3C, two-way ANOVA: main effect of condition, $F(1,12)=0.08$, $p=0.78$, partial $\eta^2=0.007$; main effect of time, $F(7,84)=2.27$, $p=0.036$, partial $\eta^2=0.16$; interaction, $F(7,84)=0.04$, $p=1.000$, partial $\eta^2=0.004$]. KD of 5αR1 in the NAc did not lead to any significant changes in the assessed behaviors, including the forced swim test [Fig. 3D, one-way ANOVA with Brown-Forsythe correction, $F(1,8.393)=1.25$, $p=0.29$, partial $\eta^2=0.08$], defensive withdrawal paradigm [Fig. 3E, one-way ANOVA, $F(1,14)=0.23$, $p=0.64$, partial $\eta^2=0.02$], or the sucrose preference test [Fig. 3F, two-way ANOVA, main effect of condition, $F(1,10)=0.46$, $p=0.51$, partial

$\eta^2$=0.04; main effect of time, F(7,70)=1.43, p=0.21, partial $\eta^2$=0.13; interaction, F(7,70)=2.61, p=0.019, partial $\eta^2$=0.21].

KD of 5αR2 in female rats did not induce significant differences regardless of the targeted area (PFC, Fig. 3G-I, and NAc, Fig. 3J-L). For PFC-targeted animals, ANOVA analyses showed no significant effects in the forced swim test [Fig. 3G, one-way ANOVA, F(1,14)=0.18, p=0.68, partial $\eta^2$=0.01], defensive withdrawal paradigm [Fig. 3H, F(1,14)=0.74, p=0.40, partial $\eta^2$=0.05], or sucrose preference test [Fig. 3I, two-way ANOVA: main effect of condition, F(1,14)=0.99, p=0.34, partial $\eta^2$=0.07; main effect of time, F(7,98)=0.82, p=0.58, partial $\eta^2$=0.06; interaction, F(7,98)=1.15, p=0.34, partial $\eta^2$=0.08]. Similarly, targeting the NAc with 5αR2-shRNA did not result in any significant differences in the forced swim test [Fig. 3J, one-way ANOVA, F(1,14)=0.004, p=0.95, partial $\eta^2=3.04 \times 10^{-4}$], defensive withdrawal paradigm [Fig. 3K, F(1,14)=1.44, p=0.25, partial $\eta^2$=0.09], or sucrose consumption in the sucrose preference test [Fig. 3L, two-way ANOVA, main effect of condition, F(1,10)=0.40, p=0.54, partial $\eta^2$=0.04; main effect of time, F(7,70)=1.45, p=0.20, partial $\eta^2$=0.13; interaction, F(7,70)=2.60, p=0.019, partial $\eta^2$=0.21].

To further investigate the impact of 5αR2 in these altered phenotypes, we generated 5αR2 full-body knock-out (KO) rats using CRISPR/Cas9-based genome editing (Fig. 4A). Validation of the model was carried out through Western blot analysis of prostate tissue (Fig. 4B), confirming effective deletion and abolition of 5αR2 expression [one-way ANOVA, F(1,4)=99.56, p<0.001, partial $\eta^2$=0.96]. Consistent with previous findings in PFC 5αR2 KD males, male KO rats exhibited a significant increase in floating time in the forced swim test [Fig. 4C, one-way ANOVA, F(1,18)=10.42, p=0.005, partial $\eta^2$=0.37], decreased time spent outside the chamber in the defensive withdrawal paradigm [Fig. 4D, one-way ANOVA with Brown-Forsythe correction, F(1,11.72)=6.42, p=0.027, partial $\eta^2$=0.26], and reduced sucrose preference [Fig. 4E, repeated measures two-way ANOVA, main effect of time, F(3,57)=5.98, p=0.001, partial $\eta^2$=0.24; main effect of genotype, F(1,19)=18.15, p<0.001, partial $\eta^2$=0.49; interaction, F(3,57)=5.35, p=0.003, partial $\eta^2$=0.22].

KO female rats showed a less pronounced phenotype, with no significant differences in the forced swim test [Fig. 4F, one-way ANOVA, F(1,16)=1.52, p=0.24, partial $\eta^2$=0.09]. However, they did exhibit a significant decrease in the time spent interacting with a foreign counterpart [Fig. 4G, one-way ANOVA, F(1,16)=8.80, p=0.009, partial $\eta^2$=0.36], while no changes were observed in the sucrose preference test [Fig. 4H, repeated measures two-way ANOVA, main effect of time, F(3,48)=3.33, p=0.03, partial $\eta^2$=0.17; main effect of genotype, F(1,16)=0.28, p=0.60, partial $\eta^2$=0.02; interaction, F(3,48)=0.09, p=0.97, partial $\eta^2$=0.01].

Building on these observations, we aimed to profile progesterone and AP levels, as well as their relative ratio, in the PFC under basal (no stress) and stress conditions. To this end, male rats with KD of either 5αR1 or 5αR2 were sacrificed 30 minutes after forced swim test exposure (stress) or under basal conditions. At baseline, progesterone levels were significantly increased by 5αR1-shRNA [Fig. 5A, one-way ANOVA with Brown-Forsythe correction, $F(1,9.524)=5.41$, $p=0.04$, partial $\eta^2=0.28$], while 5αR2-shRNA led to significantly decreased levels [Fig. 5A, $F(1,15)=8.87$, $p=0.01$, partial $\eta^2=0.37$]. AP levels were significantly reduced by KD of either 5αR1 or 5αR2 [Fig. 5B, one-way ANOVA with Brown-Forsythe correction, $F(1,11.772)=5.37$, $p=0.04$, partial $\eta^2=0.23$ for 5αR1; $F(1,14.807)=4.90$, $p=0.04$, partial $\eta^2=0.21$ for 5αR2]. However, only the downregulation of 5αR1 decreased the ratio of AP to progesterone [Fig. 5C, one-way ANOVA with Brown-Forsythe correction, $F(1,8.53)=8.95$, $p=0.02$, partial $\eta^2=0.35$ for 5αR1; $F(1,14)=4.785\times10^{-4}$, $p=0.83$, partial $\eta^2=0.003$ for 5αR2].

After stress exposure, we observed a significant upregulation of progesterone and a downregulation of AP synthesis, as well as the AP/progesterone ratio, in 5αR2 KD animals. Stress exposure significantly increased progesterone levels in 5αR2 KD rats compared to scrambled controls [Fig. 5D, one-way ANOVA with Brown-Forsythe correction, $F(1,9.804)=6.58$, $p=0.03$, partial $\eta^2=0.32$], while no significant differences were observed in 5αR1 KD rats [Fig. 5D, one-way ANOVA with Brown-Forsythe correction, $F(1,15)=2.19$, $p=0.16$, partial $\eta^2=0.13$]. Stress exposure did not affect AP levels in 5αR1 KD males but led to reduced AP levels in 5αR2 KD males [Fig. 5E, one-way ANOVA, $F(1,12)=0.01$, $p=0.93$, partial $\eta^2=0.0006$ for 5αR1; $F(1,15)=15.38$, $p=0.001$, partial $\eta^2=0.51$ for 5αR2]. Correspondingly, the AP/progesterone ratio was significantly altered in 5αR2 KD animals [Fig. 5F, one-way ANOVA with Brown-Forsythe correction, $F(1,8.34)=7.92$, $p=0.02$, partial $\eta^2=0.33$], while no significant effects were detected in 5αR1 KD animals [Fig. 5F, one-way ANOVA, $F(1,13)=0.04$, $p=0.84$, partial $\eta^2=0.003$].

To assess whether AP treatment could counteract the behavioral abnormalities observed in 5αR2 KD rats, we treated the animals with AP (IP, 6 mg/kg, 15 minutes prior to testing). Analyses of the defensive withdrawal paradigm highlighted an increase in the time spent outside the chamber [Fig. 5G, one-way ANOVA, $F(1,14)=7.95$, $p=0.01$, partial $\eta^2=0.36$], and a reduction in the latency to exit the chamber [Fig. S9A, one-way ANOVA with Brown-Forsythe correction, $F(1,9.394)=6.74$, $p=0.03$, partial $\eta^2=0.33$]. However, no differences were detected in the number of head dips [Fig. S9B, one-way ANOVA, $F(1,14)=0.51$, $p=0.49$, partial $\eta^2=0.04$]. The forced swim test showed a decrease in immobility time [Fig. 5H, one-way ANOVA, $F(1,14)=8.67$, $p=0.01$, partial $\eta^2=0.38$] and an increase in the latency to immobility [Fig. S9C, one-way ANOVA, $F(1,14)=11.17$, $p=0.01$, partial $\eta^2=0.44$]. AP treatment also increased social interaction, as measured by the total duration of interaction [Fig. 5I, one-way ANOVA, $F(1,14)=5.98$, $p=0.03$, partial $\eta^2=0.30$]. However, AP treatment did not change the latency to interact [Fig. S9D, one-way ANOVA, $F(1,14)=2.70$, $p=0.12$, partial $\eta^2=0.16$] or the time spent in facial sniffing [Fig. S9E, one-way ANOVA, $F(1,14)=0.92$, $p=0.35$, partial $\eta^2=0.06$], while it increased the time spent in anogenital sniffing [Fig. S9F, one-way ANOVA, $F(1,14)=8.23$, $p=0.01$, partial $\eta^2=0.37$]. Finally, AP treatment enhanced odor arousal [Fig. 5J, repeated measures two-way ANOVA, main effect of treatment, $F(1,14)=6.94$, $p=0.02$, partial $\eta^2=0.33$; main effect of odor, $F(1,14)=51.40$, $p<0.001$, partial $\eta^2=0.79$; interaction, $F(1,14)=20.57$, $p<0.001$, partial $\eta^2=0.60$].

In order to define the underlying mechanisms of the phenotype driven by 5αR2 downregulation, we conducted single-nucleus RNA-sequencing of the mPFC from rats stereotaxically injected with either scrambled RNA or 5αR2-shRNA, both in baseline conditions and in the context of acute stress (forced swim). Leveraging Gamma-Poisson transformation, we labeled 13 distinct clusters in our data. Using established cell markers, we identified primary cell populations, namely pyramidal neurons (PN) ranging from layers II to VI, interneurons (VIP positive and VIP negative), oligodendrocyte progenitor cells (OPCs), oligodendrocytes, astrocytes, microglia, and stromal and endothelial cells (Fig. 6A-B, S10). No discernible differences in gene expression were observed in the interneuron, stromal, and endothelial cell clusters, as well as in most PN clusters. In contrast, PN layer V, microglia, astrocytes, OPCs, and oligodendrocytes exhibited substantial transcriptomic variations between groups, suggesting their heightened susceptibility to the effects of partial 5αR2 deficiency (Data S1-S3). The downregulation of 5alpha-reductase 2 (5αR2) leads to broad and coordinated biological changes across oligodendrocyte precursor cells (OPCs), oligodendrocytes, astrocytes, microglia, and pyramidal neurons in layer 5 (PN layer 5), particularly affecting processes related to protein synthesis, energy metabolism, cellular signaling, and adhesion (Fig. 6C, Fig. S11, and Data S1-S3).

In OPCs and oligodendrocytes, 5αR2 knockdown induces significant upregulation in biological processes related to protein synthesis, including cytoplasmic translation, macromolecule biosynthesis, and peptide biosynthesis (Fig. 6C and Data S2). This suggests that these cells are ramping up protein production, likely to support myelination or cellular growth. Additionally, both cell types show enhanced mitochondrial ATP synthesis and electron transport pathways, reflecting the high energy demands required for myelin production and maintenance. Astrocytes, too, exhibit similar upregulation in protein synthesis-related processes, such as cytoplasmic translation and macromolecule biosynthesis (Fig. 6C and Data S2). This upregulation points to increased astrocytic activity, including their essential roles in supporting neural function, maintaining the blood-brain barrier, regulating extracellular ion balance, and providing metabolic support to neurons. In contrast, 5αR2 knockdown leads to significant downregulation in key biological processes within microglia and PN layer 5 (Fig. 6C and Data S1). In microglia, pathways related to immune responses and phosphorylation activities are heavily affected. Processes such as the regulation of peptidyl-tyrosine phosphorylation and protein phosphorylation are reduced, signaling a decrease in immune signaling and regulation. Microglial functions involved in adaptive immune responses and cytokine production are downregulated, potentially dampening their ability to respond to injury or inflammation. Additionally, pathways related to endoplasmic reticulum (ER) stress and the unfolded protein response are impaired, suggesting decreased capacity to manage cellular stress in microglia. In PN layer 5, downregulated biological processes include neuron projection guidance, axon guidance, and synaptic transmission modulation. This reduction indicates a diminished capacity for neuronal connectivity and synaptic communication, impairing essential aspects of neural plasticity and network formation.

The downregulation of 5αR2 significantly alters cellular components related to synaptic and neuronal structures mainly in pyramidal neurons (Fig. S11 and Data S1). These include the neuron projection, dendrite, postsynaptic density membrane, and postsynaptic specialization membrane, reflecting potential impairments in synaptic organization and neuronal connectivity. This could result in diminished synaptic communication, affecting both the formation and maintenance of neural networks. In contrast, OPCs, oligodendrocytes, and astrocytes show significant upregulation in ribosomal components (Fig. S11 and Data S2). OPCs exhibit upregulation in the cytosolic small ribosomal subunit, ribosome, and rough endoplasmic reticulum membrane, indicating their preparation for differentiation and myelination (Fig. S11 and Data S2). Oligodendrocytes show enrichment in mitochondrial structures, such as the mitochondrial respiratory chain complex and inner mitochondrial membrane, highlighting their energy demands during myelin production (Fig. S11 and Data S2). Both OPCs and oligodendrocytes show enhanced activity in focal adhesions and cell-substrate junctions, suggesting stronger cellular interactions and structural support. Astrocytes demonstrate upregulation in ribosomal components and focal adhesion-related structures, underscoring their roles in maintaining structural integrity and facilitating cellular interactions (Fig. S11 and Data S2). Their involvement in maintaining the blood-brain barrier, neuronal support, and extracellular ion balance is further highlighted through these cellular component enrichments.

The molecular functions regulated by 5aR2 knockdown reveal distinct patterns of upregulation and downregulation across different cell types (Fig. S11 and Data S1-S3). In OPCs, the most notable upregulations involve zinc ion binding and transition metal ion binding, processes essential for enzymatic activities and signaling. OPCs also show increased RNA binding activities, reflecting an active regulation of gene expression and growth control at the post-transcriptional level. Additionally, OPCs exhibit increased activity in ubiquitin-protein transferase inhibitor activity, suggesting a role in maintaining protein homeostasis (Fig. S11 and Data S2). Oligodendrocytes show a strong emphasis on ribosome-related functions, particularly RNA and rRNA binding, which reflects the high protein synthesis demands necessary for myelination. These cells also exhibit enhanced activity in regulating protein degradation and turnover, with significant upregulation in ubiquitin ligase inhibitor and ubiquitin-protein transferase inhibitor activities. Calcium and zinc ion binding are also elevated, pointing to their role in enzymatic and structural functions during myelin formation and maintenance. Astrocytes, meanwhile, show heightened molecular functions related to ion binding, including zinc, calcium, and potassium ion binding, essential for maintaining ion homeostasis and neurotransmission (Fig. S11 and Data S2). Conversely, molecular functions in PN layer 5 neurons are significantly downregulated following 5aR2 knockdown, particularly those involved in cell adhesion and receptor activities (Fig. S11 and Data S1). These changes could impair excitatory neurotransmission and neural development, potentially affecting synaptic plasticity and communication between neurons.

The downregulation of 5αR2 results in significant changes across various biological processes, molecular functions, and cellular components in OPCs, oligodendrocytes, astrocytes, microglia, and PN layer 5. These alterations impact protein synthesis, energy metabolism, immune responses, synaptic organization, and cellular signaling, with distinct effects across different cell types. While OPCs, oligodendrocytes, and astrocytes show enhanced protein synthesis and energy production, critical for maintaining neural function and myelination, PN layer 5 neurons exhibit significant impairments in synaptic connectivity, and neural signaling. These findings highlight the essential role of 5αR2 in maintaining cellular homeostasis and neural function in the central nervous system.

To investigate gene-gene interactions among differentially expressed genes (DEGs) in the comparison between 5αR2 KD rats and their controls, we employed high-dimensional Weighted Gene Co-expression Network Analysis (hdWGCNA). This approach organized the DEGs into distinct modules according to their transcriptional expression profiles. For oligodendrocyte clusters, three co-expression modules were identified (Figure S12A-B and Data S4). Each module was characterized by specific eigengenes, such as *Rps23* and *Rp32* in M1, *Zfp536* and *Ank3* in M2, and *Hibch* and *Dock9* in M3. Notably, Modules 1 and 3 were downregulated in the knockdown rats (Fig. S12C) and showed enrichment in biological processes like “Proton-driven mitochondrial ATP synthesis”, “Bleb assembly”, “Monoacylglycerol metabolic processes”, and, interestingly, “Cytoplasmic translation” (Figure S12D). Conversely, Module 2 was upregulated in knockdown rats, with enrichment in processes such as “Microtubule nucleation regulation” and ‘Protein acetylation” (Figure S12D). These findings imply that 5αR2 KD in oligodendrocytes influences various biological pathways, notably affecting mitochondrial function, cytoskeletal regulation, and metabolic activities in the mPFC.

A similar analysis of astrocyte clusters revealed four co-expression modules (Figure S13A-B and Data S4), each represented by distinct eigengenes, including *Rps29* and *Rps21* in M1, *Prex2,* and *Slc1a2* in M2, *Cnmm1* and *Emid1* in M3, and *Dgki* and *Pbx1* in M4. Modules 1 and 2 were downregulated in knockdown rats (Fig. 13C), with enrichment in “Proton-driven mitochondrial ATP synthesis”, “Cytoplasmic translation”, “L-aspartate import”, and “Purine ribonucleoside monophosphate catabolic processes” (Figure S13D). In contrast, Modules 3 and 4 were upregulated in the knockdown rats, showing enrichment in pathways related to “Blood vessel endothelial cell proliferation”, “Inhibitory synapse assembly”, and “Positive regulation of CREB transcription factor activity” (Figure S13D). This indicates that 5αR2 KD in astrocytes affects a distinct set of biological processes, impacting mitochondrial function, synaptic regulation, and various metabolic pathways in the mPFC.

Lastly, analysis of pyramidal neuron clusters in layer V revealed nine co-expression modules (Figure S14A-B and Data S4), defined by eigengenes such as *Rpl35* and *Rps29* in M1, *Ddx39b* and *Cltnn3* in M2, and *Lrrc4c* and *Col23a1* in M3. Modules 1, 2, 3, 6, and 8 were downregulated in knockdown rats (Fig. S14C), with enrichment in biological processes like “Proton-driven mitochondrial ATP synthesis”, “Cytoplasmic translation”, “Establishment of mitochondrial localization”, “Galactose metabolism”, and “Dendrite arborization” (Figure S14D). Meanwhile, Modules 4, 5, 7, and 9 were upregulated and enriched for processes involving “Calcium ion transmembrane transport”, “Dendrite morphogenesis”, “Glutamate receptor signaling”, and “Regulation of pseudopodium assembly” (Figure S14D). These results suggest that 5αR2 KD in pyramidal neurons of layer V affects an extensive range of biological processes, including mitochondrial function, synaptic signaling, and structural plasticity within the mPFC. These findings collectively highlight how 5αR2 KD impacts mitochondrial function and distinct cellular processes in several cell types, likely underscoring cell-specific roles of the neurosteroids produced by this enzyme across the mPFC.

Considering that our analyses revealed that the effects of 5αR2 are most pronounced in relation to acute stress response, we performed a detailed examination of snRNA-seq data to evaluate the effects of forced swim stress in scrambled- and 5αR2-shRNA rats. We highlighted the transcriptomic changes due to stress by ranking GO analyses related to biological process, cellular component, and molecular function across control groups (baseline and after stress exposure) and then comparing the same processes in the 5αR2-shRNA groups (Fig. 7A-B, Fig. S15, Data S7-12).

The response to stress in animals knocked down for 5αR2 differs notably from that observed in scramble-shRNA animals, with the most striking feature being a widespread downregulation of biological processes, cellular components, and molecular functions. In control animals, stress exposure results in a mixed response, with significant upregulation of processes related to protein synthesis, immune response, and metabolic activity. In neurons, particularly pyramidal neurons, there is an increase in peptide biosynthesis, ribosomal activity, and macromolecule synthesis, suggesting a robust adaptive response aimed at enhancing protein production and cellular growth (Fig. 7A, Data S8). Furthermore, glial cells, essential for maintaining neural homeostasis, also show upregulation in processes related to immune responses and metabolic support, which is crucial in helping neurons cope with stress-induced changes. In contrast, 5αR2 KD animals exhibit a different stress response profile, characterized primarily by downregulation in several critical processes. In these animals, neurons across cortical layers II-III, IV, V, and VI show a significant reduction in cytoplasmic translation, ribosome biogenesis, and gene expression, indicating a suppressed capacity for protein synthesis (Fig. 7A, Data S10). This downregulation could reflect a diminished ability to adapt to stress, impairing cellular maintenance and growth. Interneurons, both VIP-positive and VIP-negative, also exhibit reduced activity in translation-related processes, further highlighting the broad reduction in biosynthetic capacity across neuronal populations. Glial cells, including astrocytes, oligodendrocytes, OPCs, and microglia, mirror this trend with downregulated energy metabolism and ribosomal biogenesis, which suggests a decreased ability to support neurons under stress.

At the cellular component level, scramble-shRNA animals subjected to stress show an upregulation in ribosomal subunits and polysomes, consistent with their increased protein synthesis activity (Fig. 7B, Data S8). However, in KD animals, cellular components associated with protein synthesis, including the cytosolic small and large ribosomal subunits, are significantly downregulated in neurons and glial cells (Fig. 7B, Data S10). Additionally, components related to mitochondrial function, such as the mitochondrial inner and outer membranes, and key complexes involved in the respiratory chain and ATP synthesis, are similarly downregulated. This reduction in mitochondrial activity indicates a disruption in energy production, which is essential for sustaining neural and glial activity under stress. Furthermore, focal adhesions and cell-substrate junctions, crucial for cellular adhesion and signaling, are also downregulated, leading to impaired cell-to-cell communication. The downregulation of vesicle-related components in microglia and astrocytes further suggests impaired cellular communication and reduced capacity for neuroinflammatory regulation in response to stress.

In terms of molecular functions, control animals exposed to stress show an upregulation of transporter and ribosomal activities, which supports increased protein synthesis and cellular metabolism in response to stress (Fig. S15, Data S8). However, in KD animals, molecular functions related to translation, such as RNA binding, rRNA binding, and mRNA binding, are markedly downregulated across both neurons and interneurons (Fig. S15, Data S10). Additionally, glial cells in knockdown animals exhibit reduced activity in oxidative phosphorylation pathways and functions related to energy metabolism, such as oxidoreduction-driven transmembrane transporter activity and proton transmembrane transporter activity. These reductions in molecular functions are critical, as they reflect a widespread suppression of energy production and protein synthesis, which are essential for maintaining cellular function, particularly under stressful conditions.

Despite the overwhelming trend of stress-induced downregulation in 5αR2 KD animals, a few upregulated signals suggest that some adaptive processes are still active. Specifically, in neurons, there is upregulation in synaptic transmission, neuroplasticity, and neurogenesis, particularly in processes related to synaptic plasticity and excitatory transmission (Data S11). Key molecular functions, such as glutamate receptor activity, voltage-gated calcium channel activity, and GABA receptor activity, are also upregulated in neurons and interneurons. These upregulated signals suggest an adaptive attempt to maintain synaptic communication and excitatory-inhibitory balance despite the broader suppression of protein synthesis and metabolic functions. In scrambled-shRNA animals, the stress response is more balanced, with both upregulation of adaptive processes like protein synthesis and immune response and some downregulation in processes related to synaptic plasticity and neurotransmitter regulation (Data S7). However, in 5αR2 KD animals, the stress-induced downregulation dominates, affecting critical pathways involved in protein synthesis, energy production, and cellular maintenance. This stark contrast highlights the vulnerability of KD animals to stress, as their ability to upregulate adaptive processes is severely compromised. While some synaptic plasticity and neurotransmission processes remain upregulated, the overwhelming downregulation of biosynthetic and metabolic functions suggests that these animals have a reduced capacity to cope with stress, potentially leading to long-term impairments in neural function and connectivity.

In summary, stress exposure in normal animals induces a mixed response, with significant upregulation of adaptive processes, while in 5α-reductase 2 knockdown animals, stress leads to a more pronounced downregulation of key biological processes, cellular components, and molecular functions. The downregulation of protein synthesis, energy metabolism, and cellular maintenance pathways dominates the response in knockdown animals, with only a few upregulated signals related to synaptic plasticity and neurotransmission standing out. This comparison highlights the critical role of 5αR2 in enabling a robust adaptive response to stress and underscores the detrimental effects of its absence.

The hdWGCNA was performed to further assess the coexpression modules affected by stress in key cell populations. For pyramidal neuron clusters in control animals, under both stress and unstressed conditions, nine co-expression modules were identified in *Layer IV* (Fig. S16A-B and Data S13). These modules had specific eigengenes, such as *S100b* and *Rpl35* in M1, *RGD1564053* and *Dmd* in M2, and *Galnt17* and *Zfp804b* in M3. Modules 1, 3, 4, 5, 7, and 9 were downregulated in stressed rats and enriched in processes including “Oxidative phosphorylation”, “Glomerular epithelial cell development”, “Podocyte development”, “Nuclear pore complex assembly”, and “7-methylguanosine RNA capping” (Fig. S16C-D). Meanwhile, Modules 2, 6, and 8 were upregulated in stressed rats, enriched for processes such as “Proton motive force-driven ATP synthesis”, “Embryonic appendage morphogenesis”, and “Viral mRNA export” (Fig. S16C-D). These findings suggest that stress exposure in pyramidal neurons of Layer IV affects distinct biological processes, impacting mitochondrial function, cellular differentiation, and synaptic regulation in the mPFC.

The same analysis in unstressed and stressed knockdown rats revealed seven co-expression modules in pyramidal neurons of Layer IV (Fig. S17A-B and Data S14). The modules were characterized by eigengenes such as *Rps29* and *S100b* in M1, *Zfp804a* and *Sorcs3* in M2, and *Camk4* and *Cpne4* in M3. Modules 1, 3, 5, and 7 were downregulated in stressed rats, showing enrichment in pathways related to “Proton-driven mitochondrial ATP synthesis”, “Presynaptic active zone organization”, “Cardiolipin biosynthesis”, and “Cellular response to histamine” (Fig. S17C-D). On the other hand, Modules 2, 4, and 6 were upregulated, associated with processes such as “Synaptic signaling”, “Actin nucleation regulation”, “Tubular network organization”, and “Amino sugar metabolic processes” (Fig. S17C-D). These results suggest that 5αR2 knockdown, combined with stress exposure, influences biological processes related to mitochondrial function, synaptic organization, and cellular signaling in the mPFC.

Similarly, pyramidal neuron clusters in control animals under stress or unstressed conditions in Layer V revealed ten co-expression modules (Fig. S18A-B and Data S15). These modules were defined by eigengenes such as *Mt3* and *Rpl35* in M1, *Pdzdz2* and *Ryr1* in M2, and *Clasp* and *Trim39* in M3. Modules 1, 3, 4, 6, 8, 9, and 10 were downregulated in stressed rats and were enriched in processes like “Proton motive force-driven ATP synthesis”, “RNA processing”, “Axon guidance”, “Protein modification”, and “cAMP biosynthesis” (Fig. S18C-D). Conversely, Modules 2, 5, and 7 were upregulated in stressed rats, enriched in processes involving “p53-class mediator signaling”, “Retinal ganglion cell axon guidance”, and “Regulation of lipoprotein lipase activity” (Fig. S18C-D). These results suggest that stress exposure in Layer V pyramidal neurons affects biological processes impacting mitochondrial function, cellular signaling, and structural plasticity.

The same analysis for 5αR2 knockdown rats in Layer V pyramidal neurons revealed eight co-expression modules (Fig. S19A-B and Data S16). Modules had eigengenes such as *Zc3h10* and *Rps27a* in M1, *Prkcg* and *Tmem132b* in M2, and *Flrt3* and *Kcnq1* in M3. Modules 1, 3, 4, and 5 were downregulated, with enrichment in “Cytoplasmic translation”, “Spliceosomal complex assembly”, “T-circle formation”, and “Regulation of glial apoptosis” (Fig. S19C-D). Modules 2, 6, 7, and 8 were upregulated and were enriched in pathways related to synaptic signaling, presynaptic organization, growth factor signaling, and nucleotide metabolism (Fig. S19C-D). These findings indicate that 5αR2 knockdown, combined with stress, affects processes linked to mitochondrial function, synaptic organization, and cellular signaling.

For the first cluster of pyramidal neurons in Layer VI, hdWGCNA revealed six co-expression modules in control animals (Fig. S20A-B and Data S17). Modules were characterized by eigengenes such as *Rps21* and *Rpl35* in M1, *Slc24a2* and *Kcnd2* in M2, and *Ppp2r3a* and *Igf1r* in M3. Modules 1, 2, 3, and 4 were downregulated in stressed rats, enriched for “Oxidative phosphorylation”, “Lipid kinase activity regulation”, “Mitochondrial protein processing”, and “Synaptic transmission” (Fig. S20C-D). Modules 5 and 6 were upregulated in stressed conditions and enriched in “Synaptic vesicle uncoating”, “Adherens junction maintenance”, and “Plasma membrane transport” (Fig. S20C-D). These results suggest that stress in Layer VI pyramidal neurons affects processes related to mitochondrial function, synaptic regulation, and intracellular transport.

In knockdown rats, the first cluster of pyramidal neurons in Layer VI revealed seven co-expression modules (Figure S21A-B and Data S18). Eigengenes included *Mt3* and *S100b* in M1, *Adgrb2* and *Ngef* in M2, and *Fgf14* and *Frm4p4* in M3. Modules 1, 2, 3, 5, and 6 were downregulated in stressed rats, and enriched for processes like “ATP synthesis”, “X chromosome inactivation regulation”, “Histone acetylation”, and “Calcium ion transport” (Fig. S21C-D). Modules 4 and 7 were upregulated and enriched for processes related to “Muscle cell differentiation”, “Synaptic membrane organization”, and “CREB transcription factor activity regulation“ (Fig. S21C-D). These findings indicate that 5αR2 knockdown in Layer VI pyramidal neurons, combined with stress exposure, affects mitochondrial function, gene regulation, and synaptic signaling.

The analysis of the second cluster of pyramidal neurons in *Layer VI* of control rats revealed eight co-expression modules (Fig. S22A-B and Data S19), with eigengenes like *Rpl35* and *Rps23* in M1, *Pak5* and *Nfkm* in M2, and *Man1a1* and *Trim33* in M3. Modules 1, 2, 4, 5, 6, and 8 were downregulated, enriched in “Oxidative phosphorylation”, “GDP-mannose metabolism”, “Histone acetylation”, “Protein tyrosine kinase activity”, and “Autophagosome docking” (Fig. S22C-D). Modules 3 and 7 were upregulated and enriched in “Mitochondrial autophagy”, “rRNA processing”, and “Presynaptic active zone organization” (Fig. S22C-D). These findings imply that stress in Layer VI pyramidal neurons impacts mitochondrial function, autophagy, and synaptic organization.

For the second cluster of pyramidal neurons in Layer VI, nine co-expression modules were identified (Fig. S23A-B and Data S20). Specific eigengenes included *Tmsb4x* and *Cst3* in M1, *Ccser1* and *Adgrb3* in M2, and *Akap8l* and *Ogt* in M3. Modules 1, 3, 5, 6, 7, and 9 were downregulated in stressed rats, with enrichment in pathways like “Oxidative phosphorylation”, “pre-mRNA processing”, “Response to sterol”, “Ganglioside biosynthesis”, and “Regulation of synaptic vesicle exocytosis” (Fig. S23C-D). Conversely, Modules 2, 4, and 8 were upregulated, with enrichment in processes related to “Voltage-gated sodium channel activity” and “Mitotic spindle establishment” (Fig. S23C-D). These results suggest that 5αR2 KD in pyramidal neurons of Layer VI, coupled with acute stress, affects biological processes related to mitochondrial function, synaptic regulation, and cellular differentiation within the mPFC.

A similar hdWGCNA was conducted for the VIP negative interneuron clusters under control and stressed conditions, revealing six co-expression modules (Fig. S24A-B and Data S21). Modules were characterized by eigengenes such as *Rpl35* and *Cst3* in M1*, Lrrc4c* and *Fgf12* in M2, and *Schip1* and *Grid2* in M3. Modules 1, 3, 4, and 5 were downregulated in stressed rats, enriched for processes like “Proton motive force-driven ATP synthesis”, “Chondroitin sulfate proteoglycan metabolism”, and “Positive regulation of protein localization to the telomere” (Fig. S24C-D). Conversely, Modules 2 and 6 were upregulated in stressed conditions, with enrichment in pathways involving “Cellular response to leucine”, “Protein localization to cytoplasmic stress granules”, and “Regulation of glial cell differentiation” (Fig. S24C-D). These findings indicate that stress exposure in VIP-negative interneurons primarily impacts mitochondrial function, cellular differentiation, and metabolic processes.

The same analysis for VIP-negative interneurons in 5αR2 KD rats identified four co-expression modules (Fig. S25A-B and Data S22). Specific eigengenes included *Kcnh7* and *Lrrc4c* in M1, *Grid2* and *Nrxn3* in M2, and *Cst3* and *S100b* in M3. Modules 1 and 3 were downregulated in stressed rats, enriched in biological processes such as “Ketone biosynthetic regulation”, “Gamma-aminobutyric acid metabolism”, “Proton-driven ATP synthesis”, and “Cytoplasmic translation” (Fig. S25C-D). Modules 2 and 4 were upregulated, showing enrichment in pathways related to “Postsynaptic density organization”, “Sodium ion transport regulation”, and “Regulation of cardiac muscle cell action potential” (Fig. S25C-D). These findings indicate that 5αR2 KD in VIP negative interneurons, combined with acute stress, influences mitochondrial activity, synaptic regulation, and ion transport processes.

For VIP positive interneurons, hdWGCNA identified five co-expression modules under control and stressed conditions (Fig. S26A-B and Data S23). Modules were characterized by eigengenes including *Rps23* and *Rpl35* in M1, *Gm7* and *Dscam* in M2, and *Cyct* and *Rims1* in M3. Modules 1, 3, and 4 were downregulated, enriched for pathways involving “ATP synthesis”, “Blood-brain barrier regulation”, and “Collateral sprouting” (Fig. S26C-D). Modules 2 and 5 were upregulated and enriched in processes related to “Neurotransmitter receptor activity regulation”, “Cardiac muscle hypertrophy”, and “Synaptic adhesion” (Fig. S26C-D). These results imply that stress in VIP positive interneurons mainly affects mitochondrial function, neurotransmitter signaling, and structural processes.

In 5αR2 KD VIP positive interneurons, six co-expression modules were identified (Fig. S27A-B and Data S24). Eigengenes included *Zc3h10* and *Rps29* in M1, *Pde4d* and *Gpn3* in M2, and *Atf1* and *Phc4* in M3. Modules 1, 3, and 4 were downregulated, with enrichment in processes like “ATP synthesis”, “RNA secondary structure unwinding”, and “Synapse assembly” (Fig. S27C-D). Modules 2, 5, and 6 were upregulated, enriched in pathways involving “Histone deacetylase activity regulation”, “Sensory perception of pain”, and “Mitochondrial autophagy” (Fig. S27C-D). This indicates that 5αR2 KD, combined with stress, primarily affects mitochondrial function, synaptic plasticity, and cellular homeostasis.

The analysis of oligodendrocyte clusters in control rats under stress revealed three co-expression modules (Fig. S28A-B and Data S25). Specific eigengenes included *Rpl32* and *Rps23* in M1, *Igf1r* and *Grid4* in M2, and *Atosa* and *Ptpro* in M3. Modules 1 and 2 were downregulated and enriched in pathways like “Cytoplasmic translation”, “ATP synthesis”, and “Dendritic spine maintenance” (Fig. S28C-D). Module 3 was upregulated and involved in “TORC2 signaling”, “Calcineurin-NFAT signaling”, and “Calcineurin-mediated signaling” (Fig. S28C-D). These findings indicate that stress in oligodendrocytes affects cellular signaling, mitochondrial function, and structural processes.

In 5αR2 KD oligodendrocytes, four co-expression modules were identified (Fig. S29A-B and Data S26). Specific eigengenes included *Uba52* and *Rps29* in M1, *Cac39l* and *ApoD* in M2, and *Zip536* and *Creb3* in M3. Modules 1, 2, and 3 were downregulated, enriched in pathways like “ATP synthesis” and “ncRNA export” (Fig. S29C-D). Module 4 was upregulated and enriched for processes involving “Ion transport”, “Cation transport”, and “Sister chromatid cohesion” (Fig. S29C-D). This indicates that 5αR2 KD, combined with stress, influences ion transport, mitochondrial function, and cellular differentiation.

Lastly, comparing the oligodendrocyte precursor cell (OPC) clusters in control rats under stress revealed four co-expression modules in the mPFC (Fig. S30A-B and Data S27). Each module was characterized by specific eigengenes, such as *Rps27a* and *Rpl32* in M1, *Tns3* and *Arhgap24* in M2, and *Dlg2* and *Nxph1* in M3. Modules 1 and 2 were downregulated and enriched in processes including “Cytoplasmic translation”, “ATP synthesis”, “miRNA metabolic process’, and “Adherens junction maintenance” (Fig. S30C-D). Modules 3 and 4 were upregulated and enriched for biological processes such as “Regulation of trans-synaptic signaling”, “Synaptic transmission (GABAergic)”, and “Regulation of chemotaxis” (Fig. S30C-D). These findings suggest that acute stress in OPCs impacts mitochondrial function, synaptic signaling, and cellular migration.

In 5αR2 KD OPCs, five co-expression modules were identified (Fig. S31A-B and Data S28). Modules included eigengenes such as *Rpl32* and *Rps29* in M1, *Epb42f2* and *Tns3* in M2, and *Csmd3* and *Marchf1* in M3. Modules 1 and 3 were downregulated in stressed rats, enriched for pathways like “ATP synthesis”, “Oxidative phosphorylation”, “ncRNA metabolic process”, and “Synaptic transmission” (Fig. S31C-D). Modules 2, 4, and 5 were upregulated and involved in processes such as “Golgi to endosome transport”, “Adherens junction assembly”, and “Podocyte cell migration” (Fig. S31C-D). These findings indicate that 5αR2 KD in OPCs, combined with stress, influences mitochondrial function, synaptic plasticity, and cellular adhesion.

### Supplementary Figures

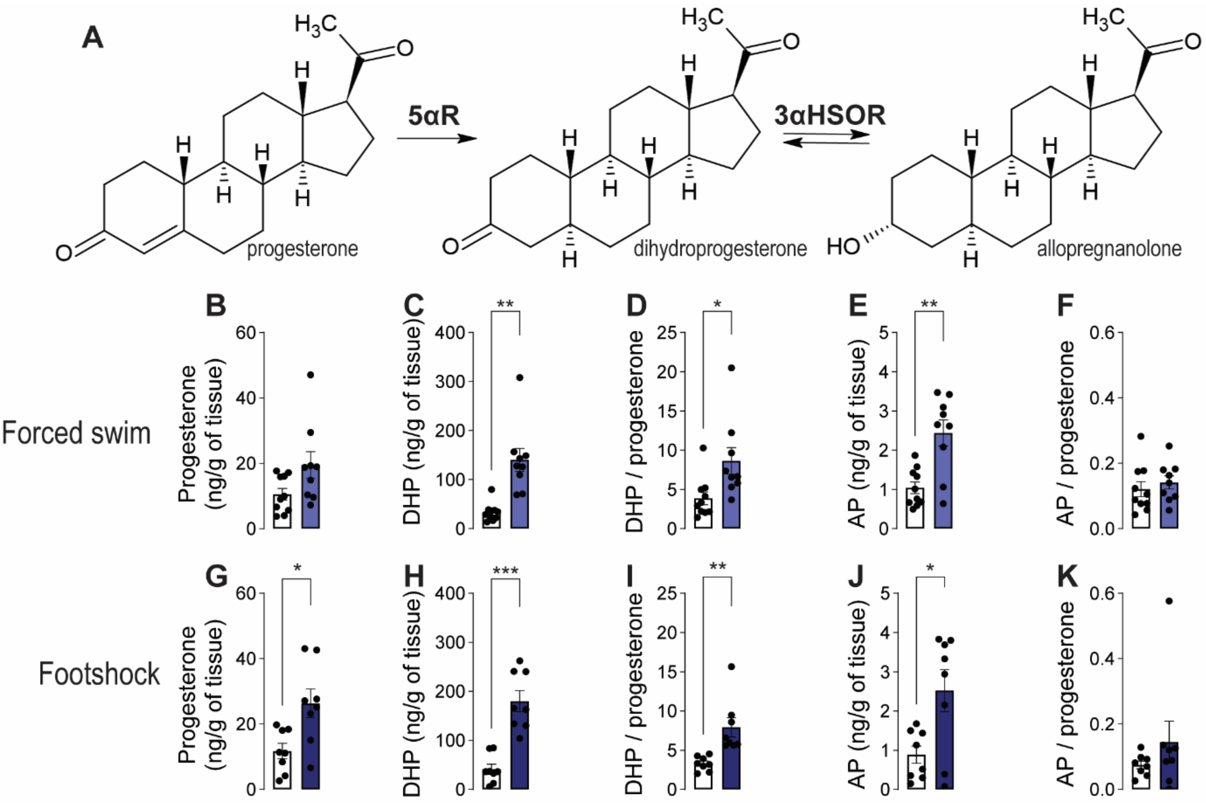


**Figure S1. Acute stressors upregulate allopregnanolone (AP) levels in the medial prefrontal cortex (mPFC) of male rats**. (A) The conversion of progesterone to AP occurs through a two-step enzymatic pathway: 5α-reductase (5αR) transforms progesterone into dihydroprogesterone (DHP), and 3α-hydroxysteroid oxidoreductase (3α-HSOR) subsequently converts DHP into AP. The process catalyzed by 5αR is unidirectional and serves as the rate-limiting step of this pathway. We measured the content of these steroids in male rats subjected to either forced swim (B-F; light blue columns) or footshock (G-K; dark blue columns) at 30 minutes of their completion. Control rats (represented by white columns) remained undisturbed in their home cages. Forced swim exposure resulted in a marginal increase in progesterone levels (B, p=0.06), along with significant elevations in DHP (C) and the DHP/progesterone ratio (D). As expected, this stressor also led to an increase in AP levels (E), although it did not significantly affect the AP/progesterone ratio (F). Footshock similarly caused a significant rise in progesterone (G), DHP (H), and AP levels (J), as well as a significant increase in the DHP/progesterone ratio (I). However, the AP/progesterone ratio (K) was not significantly affected. For further details, refer to Supplementary Results. Data are presented as mean ± SEM (n=8-10/group). Light blue indicates forced swim, dark blue indicates footshock. *, $p<0.05$; **, $p<0.01$; ***, $p<0.001$ for comparisons indicated by brackets.

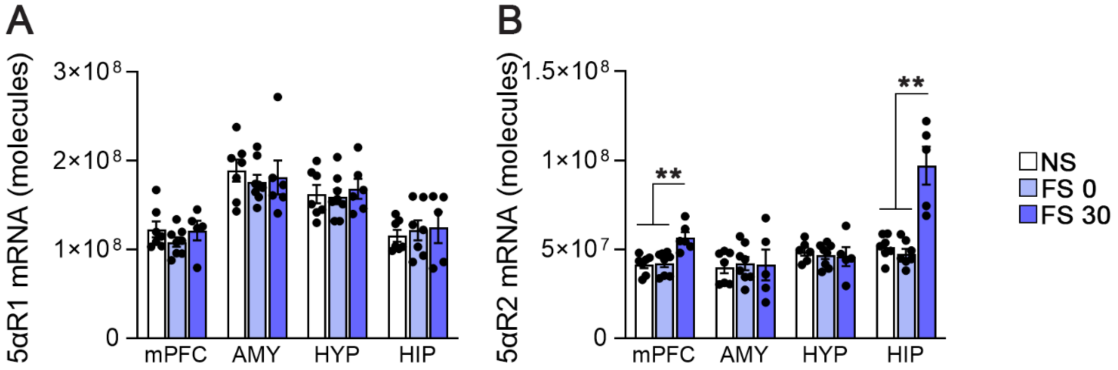


**Figure S2: Acute forced swim stress upregulates cortical and hippocampal 5αR2 mRNA levels.** Male rats were exposed to forced swim (FS) and were then sacrificed immediately after the task (FS 0) or 30 minutes later (FS 30). The control group (NS) was left undisturbed in the home cage. (A-B) 5αR1 (A) and 5αR2 (B) mRNA levels were evaluated by qRT-PCR in extracts from: medial prefrontal cortex (mPFC), amygdala (AMY), hypothalamus (HYP), and hippocampus (HIP). 30 minutes after stress exposure, 5αR2 mRNA levels were significantly upregulated in mPFC and HIP. Data are presented as mean ± SEM (n=6-8/group). **, $p<0.01$ for comparisons indicated by brackets.

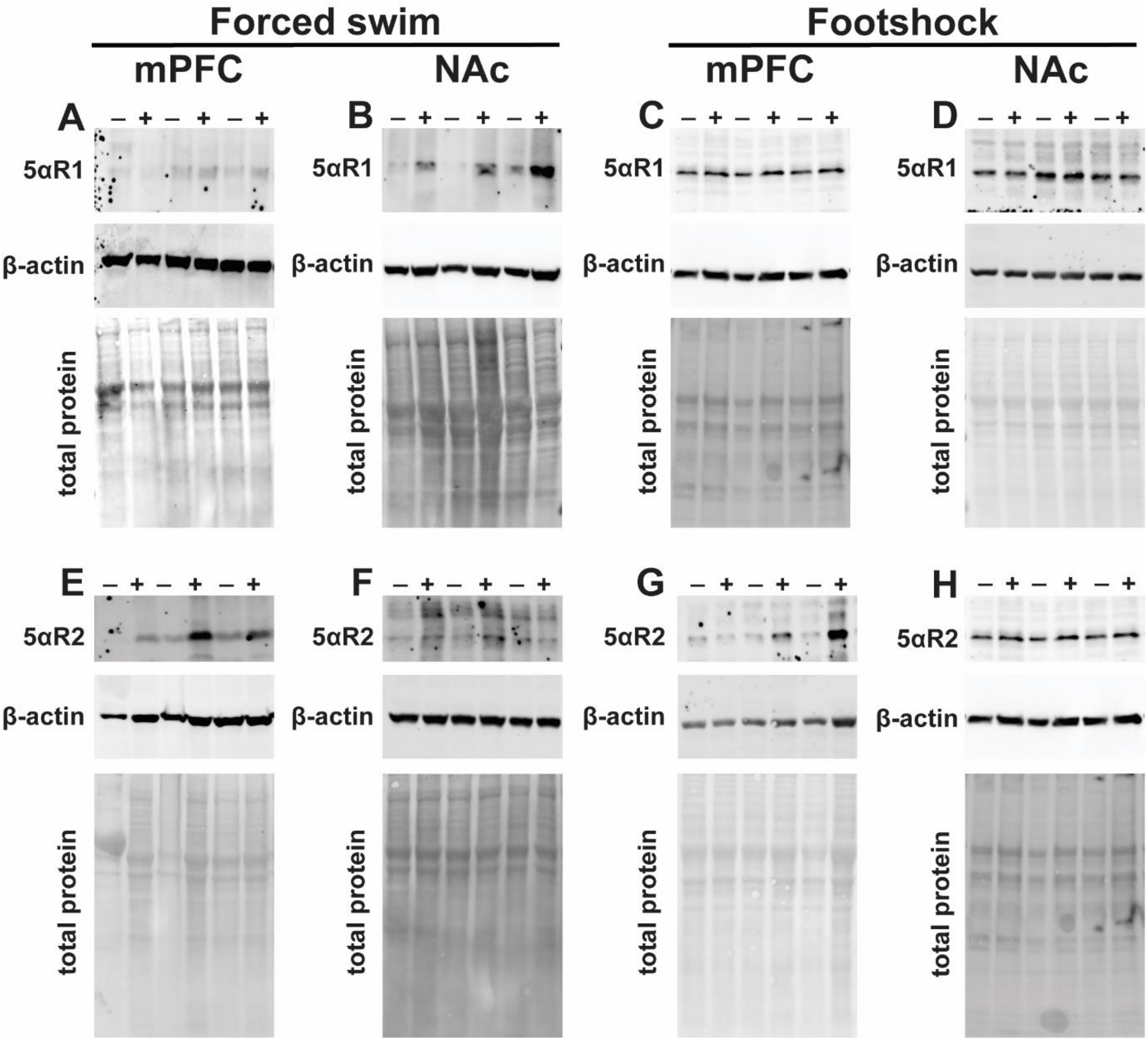


**Fig. S3: Representative Western blot images** used for the quantification shown in Fig. 1. (A-D) Representative images used for the quantification of 5αR1 in the medial prefrontal cortex (mPFC, A and C) and nucleus accumbens (NAc, B and D) of rats exposed to either forced swim (A-B) or footshock (C-D). (E-H) Representative images used for the quantification of 5αR2 in the mPFC (E and G) and NAc (F and H) of rats exposed to either forced swim (E-F) or footshock (G-H). For every panel, shown the protein of interest (5αR1 and 5αR2), β-actin, and the total loaded proteins (obtained through stain-free or ponceau-red).

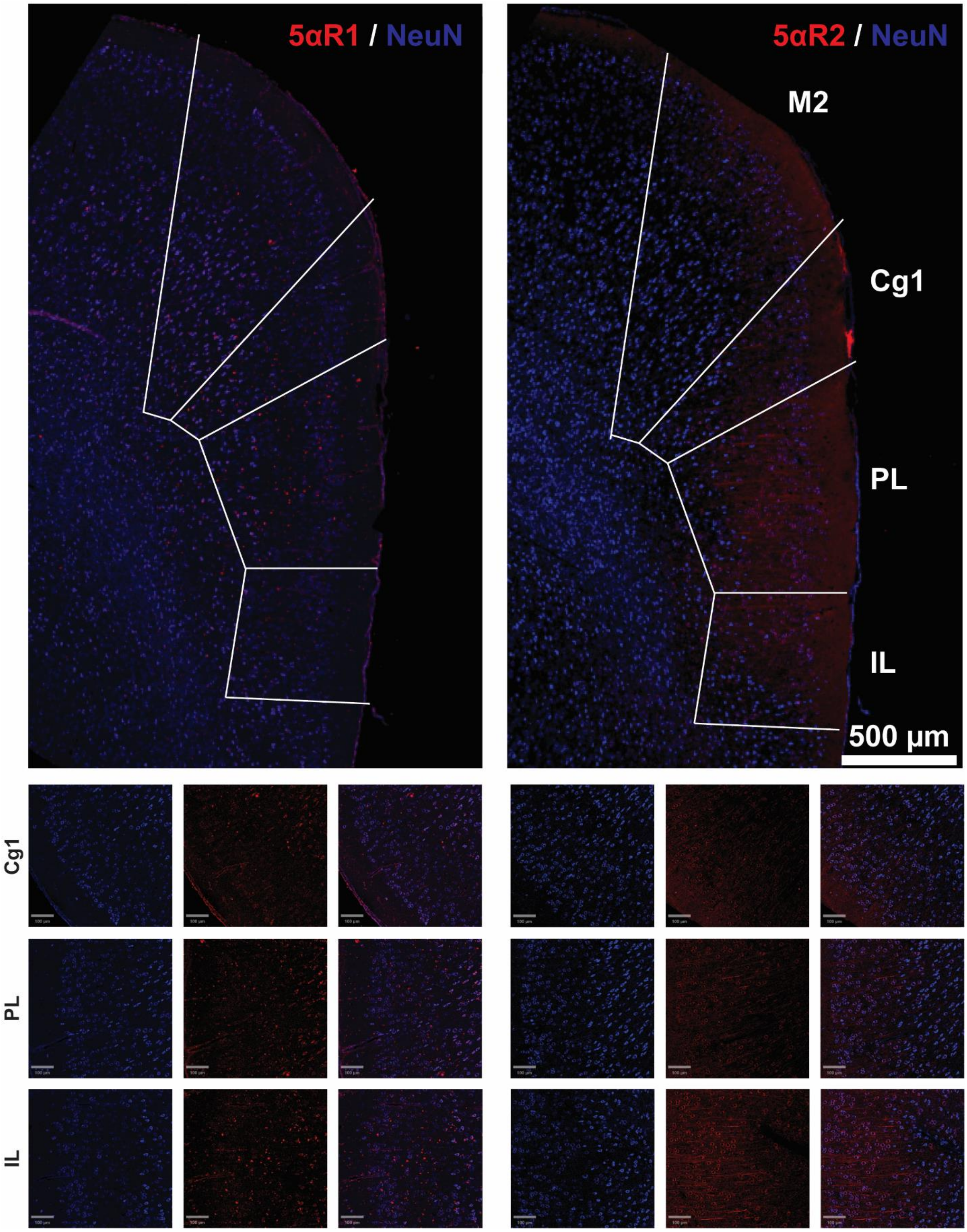


**Figure S4: Localization and expression of 5aR isoforms.** Representative overview images (above) and magnification (below) of the immunofluorescence co-staining of NeuN and 5αR1 (left) or 5αR2 (right). Abbreviations: M2, secondary motor cortex; Cg1, cingulate cortex; PL, prelimbic cortex; IL, infralimbic cortex.

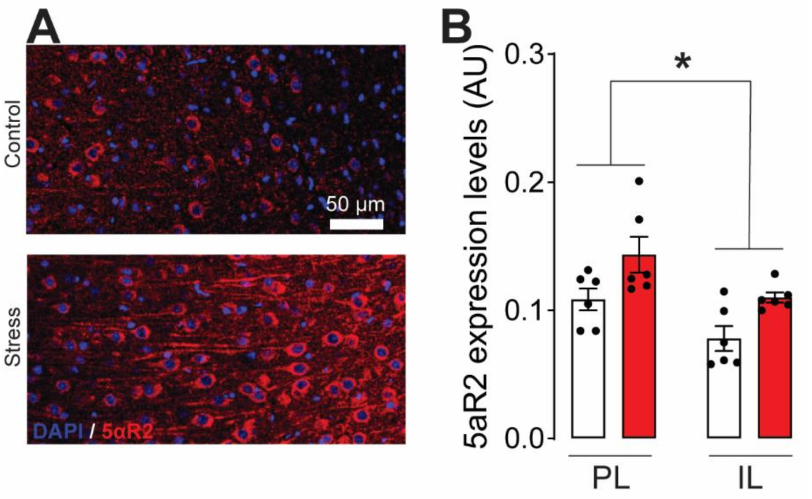


**Figure S5: Acute forced swim stress upregulates cortical 5αR2 protein levels.** Male rats were exposed to forced swim and were then sacrificed 30 minutes after the task (stress, red bars). The control group (white bars) was left undisturbed in the home cage. (A) Representative images of 5αR2 staining in the cortex of the experimental animals. (B) While the infralimbic cortex (IL) expressed lower levels of 5αR2 than the prelimbic cortex (PL), stress exposure significantly upregulated 5αR2 levels irrespective of the considered area. Data are presented as mean ± SEM (n=6/group). **, $p<0.01$ for comparisons indicated by brackets.

.

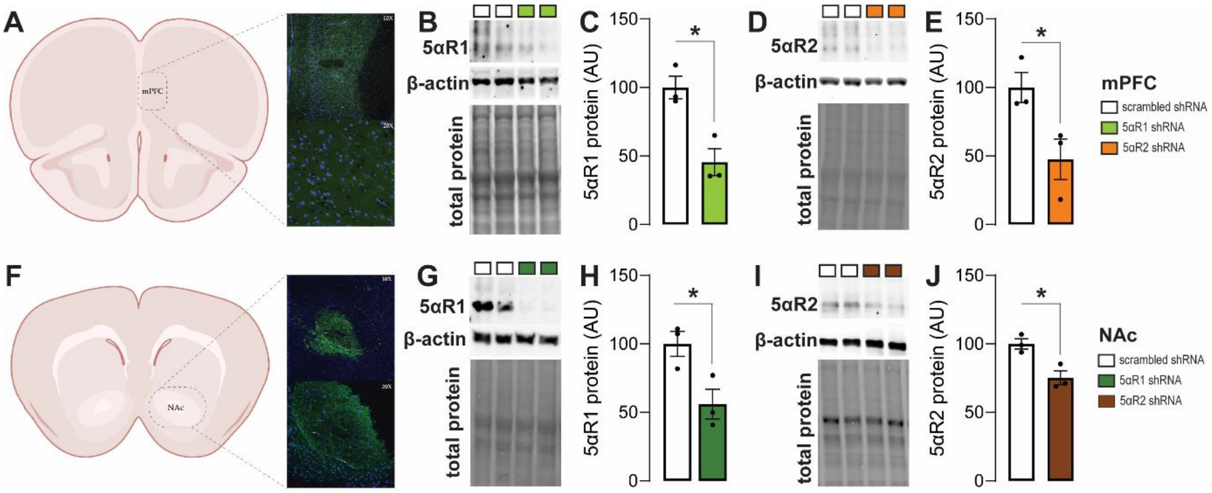


**Figure S6: Targeted downregulation of 5αR1 and 5αR2 in medial prefrontal cortex (mPFC) and nucleus accumbens (NAc).** (A-E) Targeting the mPFC, as shown by representative images of the AAV5 spreading (A, visualized as GFP staining), successfully reduced 5αR1 (B-C) or 5αR2 (D-e) expression in PFC. (F-J) Targeting the NAc, as shown by representative images of the GFP staining (D), successfully reduced 5αR1 (G-H) or 5αR2 (I-J) expression in the NAc Data in C, E, H, and J were obtained by quantification of protein levels by Western blot analyses (representative images are reported in panels B, D, G, and I). Data are reported as mean ± SEM (n=3/group). *, $p<0.05$.

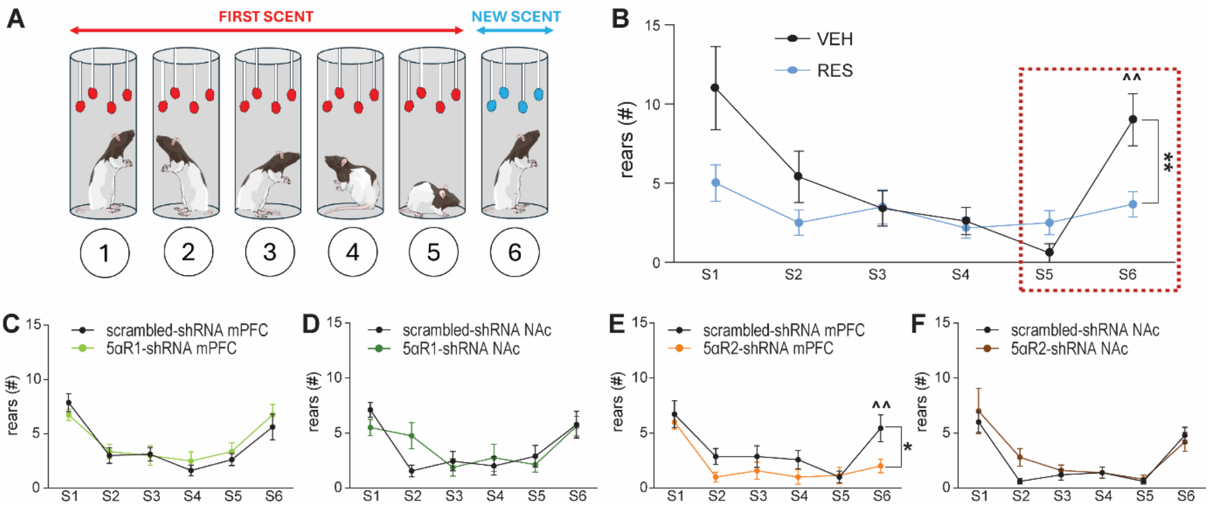


**Figure S7. Description and validation of the olfactory arousal test.** (A) Rats were placed in a cylindrical enclosure containing four long cotton swabs imbibed with a scent. Each rat remained in the cylinder for 2 minutes before being removed for a 30-second interval, after which it was placed back in the cylinder with four new swabs impregnated with the same odor. This procedure was repeated over five sessions, resulting in a gradual decline in the number of rears. For the sixth and final session, the swabs were imbibed with a new odor, leading to a renewed arousal response. The sequence of odors was counterbalanced across rats to avoid any bias due to intrinsic preferences for specific odors. The number of rears in the fifth and sixth sessions was compared using a two-way repeated-measures ANOVA to examine the arousal evoked by the novel olfactory stimulus. (B) To validate the protocol, we administered a low-dose reserpine regimen (1 mg/kg/day, SC, for six days), which was previously shown to elicit no change in locomotor activity (*39*). This drug was chosen for its ability to reduce dopamine release, thereby decreasing motivational drive. Indeed, reserpine produced a significant decrease in the number of rears induced by the exposure to the novel scent. (C-D) Knockdown of 5αR1 in the medial prefrontal cortex (mPFC) and nucleus accumbens (NAc) did not modify olfactory arousal; (E) Conversely, knockdown of 5αR2 in the mPFC produced a significant reduction in olfactory arousal. (F) Finally, no change was observed in olfactory arousal in rats subjected to 5αR2 knockdown in the NAc. Data are presented as mean ± SEM (n=8-12/group). *, $p<0.05$; **, $p<0.01$ for comparisons indicated by brackets. ^^, $p<0.01$ for comparisons between old (S5) and new (S6) odor for vehicle animals.

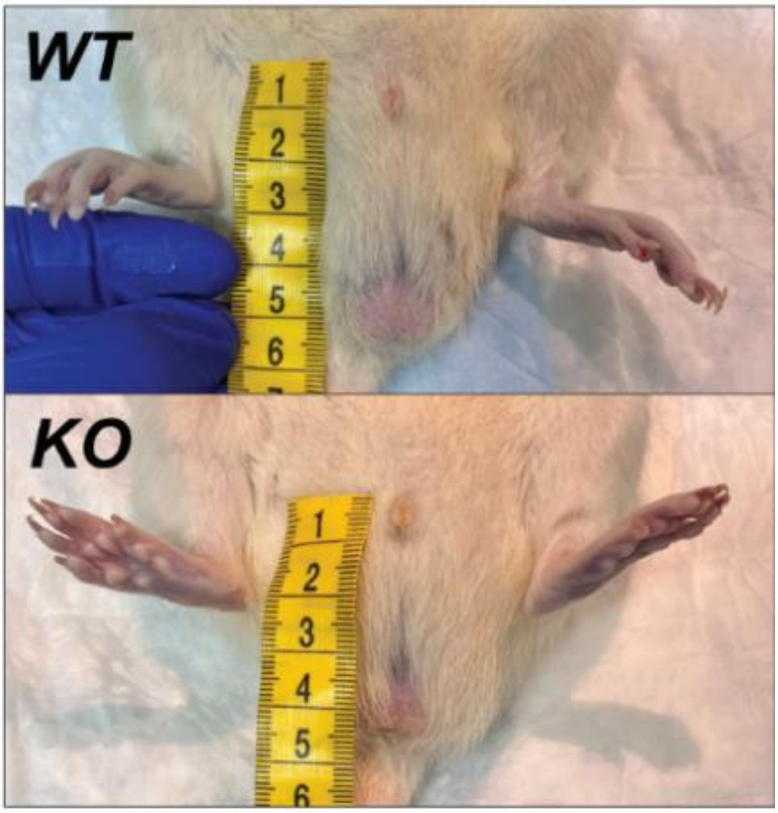


**Figure S8**. Anogenital distance comparison between WT (wild-type) and KO (5αR2 knockout) male rats. The KO male rats exhibit a significantly shorter anogenital distance compared to WT, indicative of impaired masculinization.

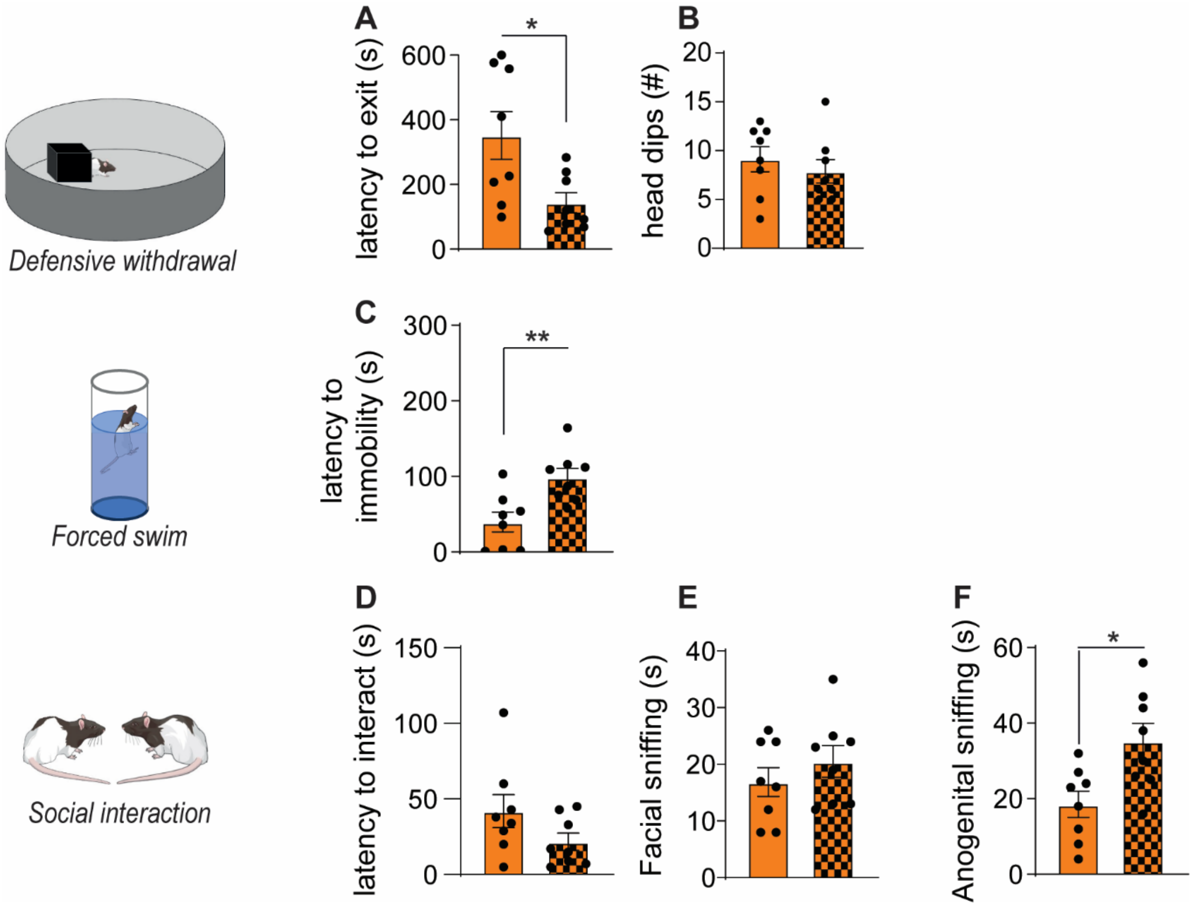


**Fig. S9. Effects of systemic administration of allopregnanolone in 5αR2 deficient rats.** To explore the effects of 5αR2 downregulation, we tested whether allopregnanolone (AP) treatment could rescue the observed phenotype. AP was administered 15 minutes before testing by IP injection at a dose of 6mg/kg. (A-B) Treatment with AP decreased the latency to exit the chamber (A) in the defensive withdrawal test, while no differences were detected in the number of head dips (B). (C) AP treatment increased the latency to immobility in the forced swim test- (D-F) AP treatment did not affect the latency to interact (D) or the time spent in facial sniffing (E), while it increased the time spent in anogenital sniffing (F) during the social interaction test. Data are presented as mean ± SEM (n=8/group) *, $p<0.05$; **, $p<0.01$ for comparisons indicated by brackets.

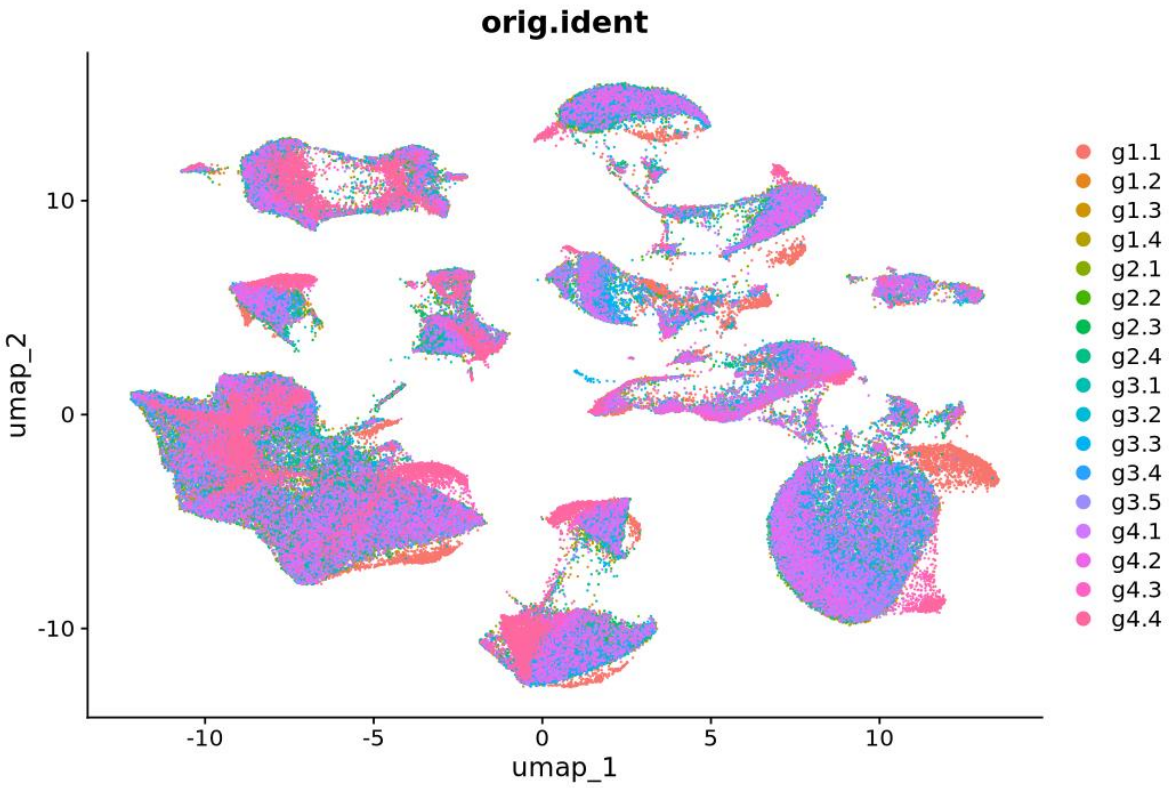


**Figure S10. Uniform Manifold Approximation and Projection (UMAP) of all cell types in all four groups.** Overlay of the UMAP of all the individual samples used in this analysis. Abbreviations: g1, non-stressed scrambled-shRNA; g2, stressed scrambled-shRNA; g3, non-stressed 5αR2-shRNA; g4, stressed 5αR2-shRNA.

### cellular component

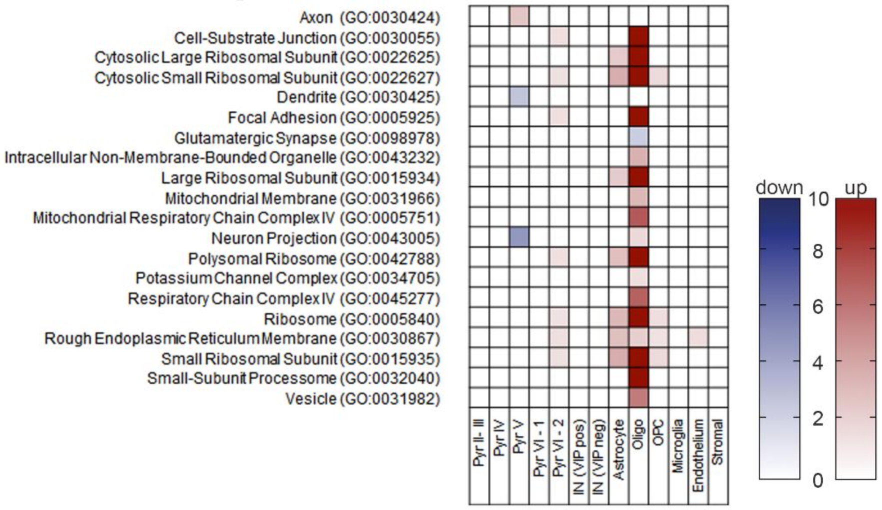


### molecular function

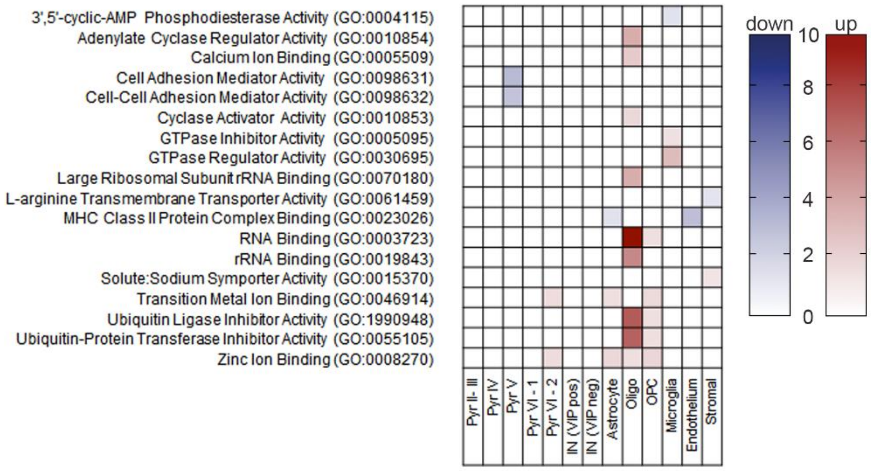


**Figure S11: Gene Ontology cellular component and molecular function analyses of cortical tissues of 5αR2 knock-down and scrambled control rats.** Gene ontology analysis for cellular component (top) and molecular function (bottom) highlighting the most differential ontologies among these cell clusters in presence or in absence of 5αR2. Abbreviations: Pyr, pyramidal neurons; IN, interneurons; Oligo, oligodendrocytes; OPC, oligodendrocyte precursor cell.

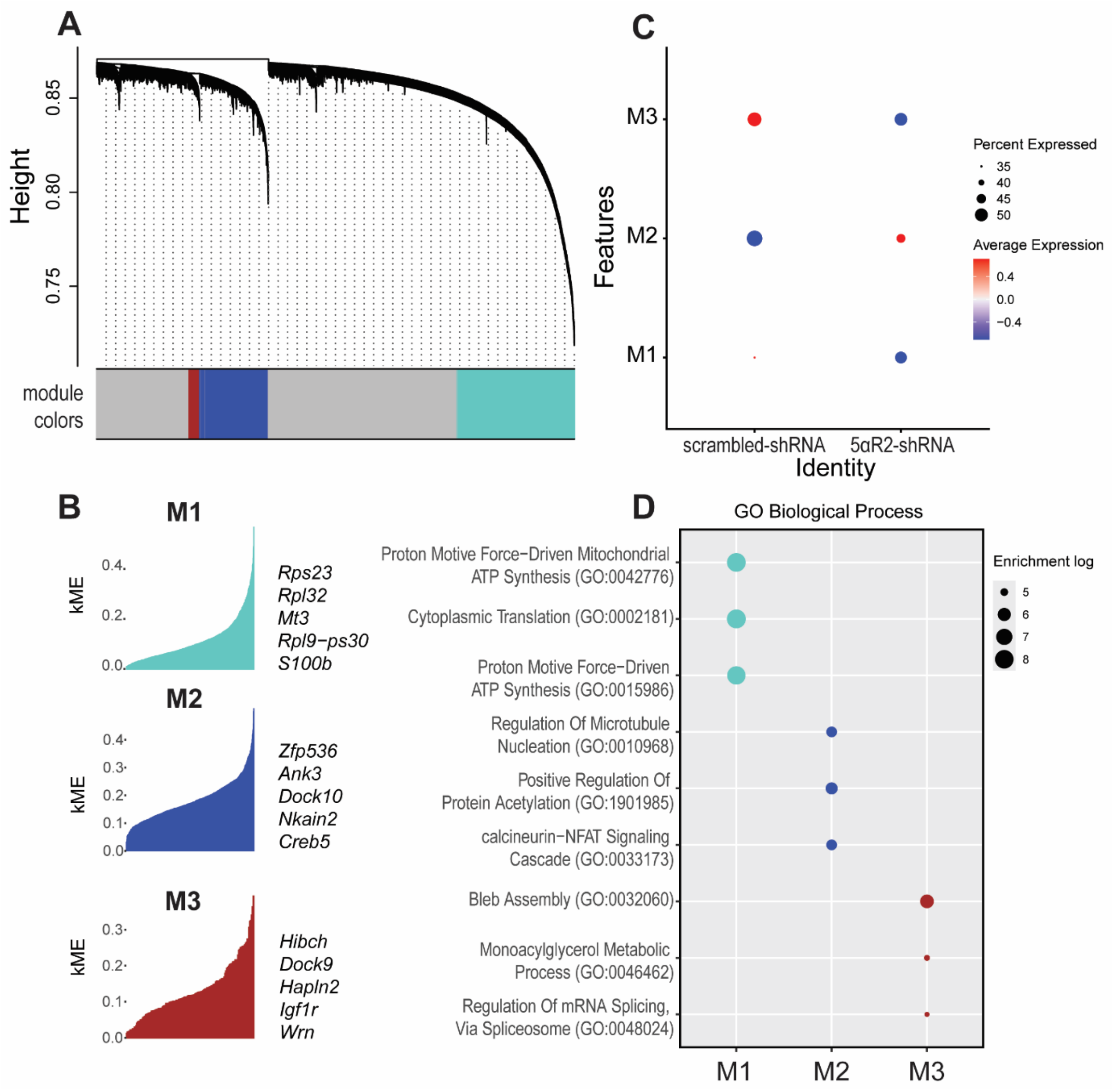


**Figure S12: Gene coexpression analysis of oligodendrocytes in the mPFC of male rats treated with either 5αR2-shRNA (knockdown) or scrambled-shRNA (controls).** (A) The dendrogram clusters differentially expressed genes into distinct modules, representing the network modular organization. The lower part of the panel shows the module colors that identify three distinct modules, designated as M1 (cyan), M2 (blue), and M3 (red). (B) The three modules are shown with the top five eigengenes for each module. (C) Module expression profiles with comparisons between knockdown and control samples, with red and blue representing up- and downregulation, respectively. (D) GO Biological Process pathway enrichment analysis of each coexpression module as calculated by hdWGCNA. The top three processes are indicated in each module.

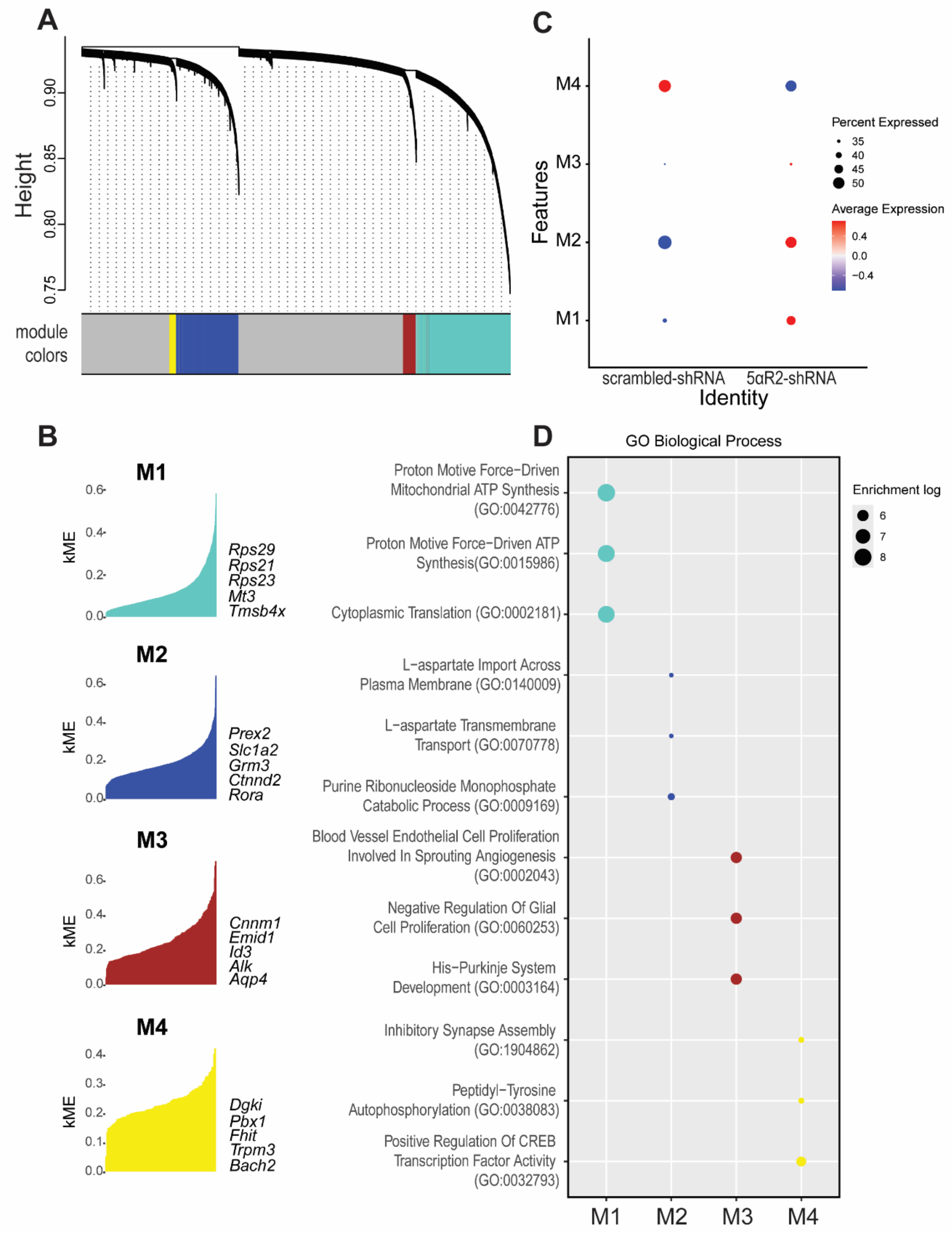


**Figure S13: Gene coexpression analysis of astrocytes in the mPFC of male rats treated with either 5αR2-shRNA (knockdown) or scrambled-shRNA (controls).** (A) The dendrogram illustrates the network modular organization of differentially expressed genes comparing mPFC samples from knockdown and control rats. (B) Four modules were identified, with the top 5 eigengenes reported. (C) Module expression profiles with comparisons between knockdown and control samples, with red and blue representing up- and downregulation, respectively. (D) GO Biological Process pathway enrichment analysis of each coexpression module as calculated by hdWGCNA. The top three processes are indicated in each module.

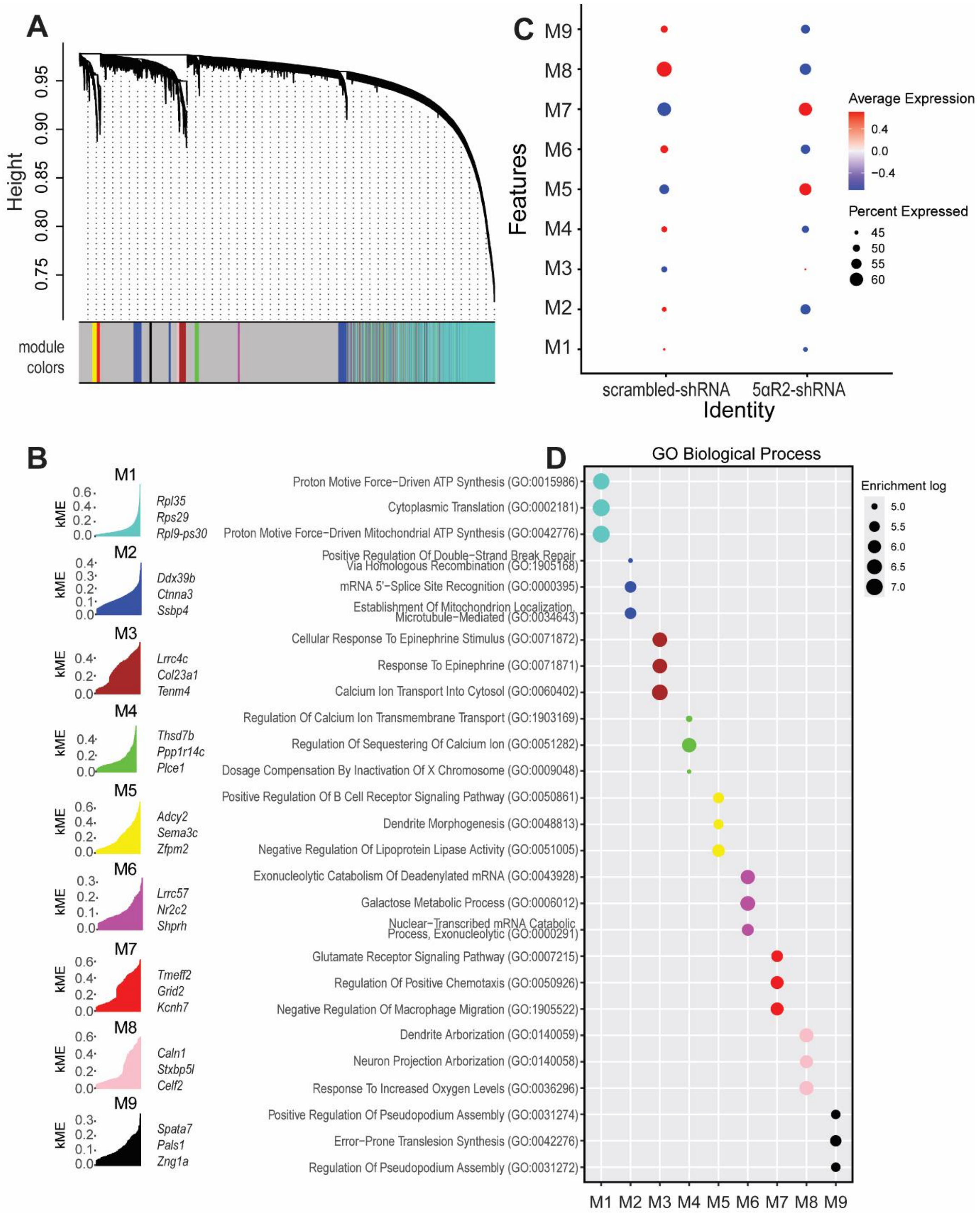


**Figure S14: Gene coexpression analysis of pyramidal neurons in layer V in the mPFC of male rats treated with either 5αR2-shRNA (knockdown) or scrambled-shRNA (controls).** (A) The dendrogram illustrates the network modular organization of differentially expressed genes comparing mPFC samples from knockdown and control rats. (B) Nine modules were identified, with the top 3 eigengenes reported. (C) Module expression profiles with comparisons between knockdown and control samples, with red and blue representing up- and downregulation, respectively. (D) GO Biological Process pathway enrichment analysis of each coexpression module as calculated by hdWGCNA. The top three processes are indicated in each module.

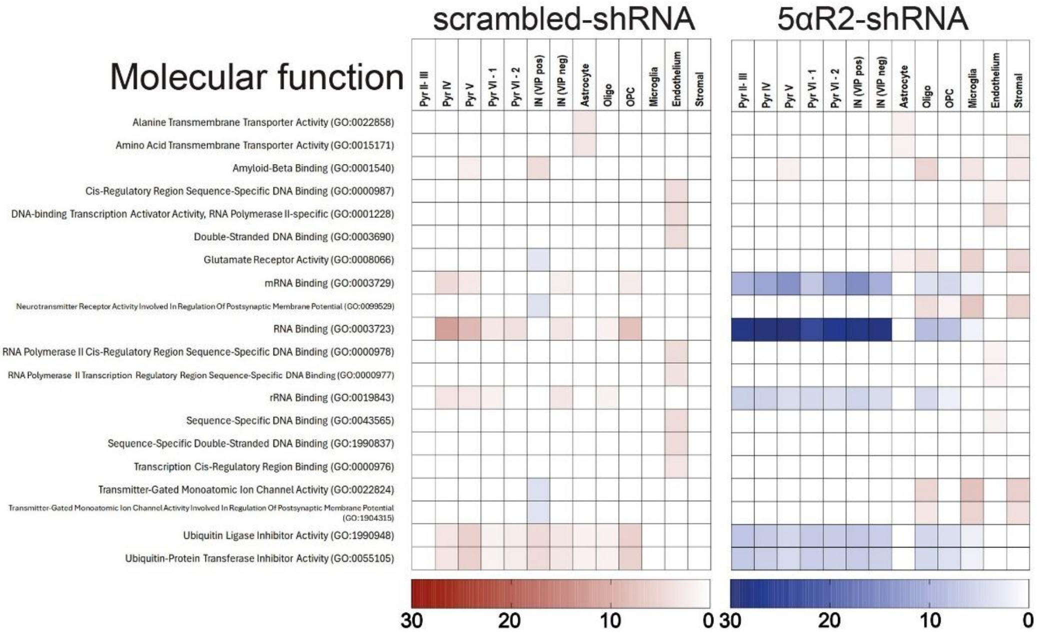


**Figure S15: Gene Ontology molecular function analyses of cortical tissues of 5αR2 knock-down and scrambled control rats exposed to acute stress.** Gene ontology analysis for molecular function (bottom) highlighting the most differential ontologies among these cell clusters. Two comparisons are shown side by side, on the left differences induced by stress exposure in prefrontal cortex (PFC) of control rats, on the right the same behavioral paradigm applied to rats knocked-down for 5αR2 in the PFC. Abbreviations: Pyr, pyramidal neurons; IN, interneurons; Oligo, oligodendrocytes; OPC, oligodendrocyte precursor cell.

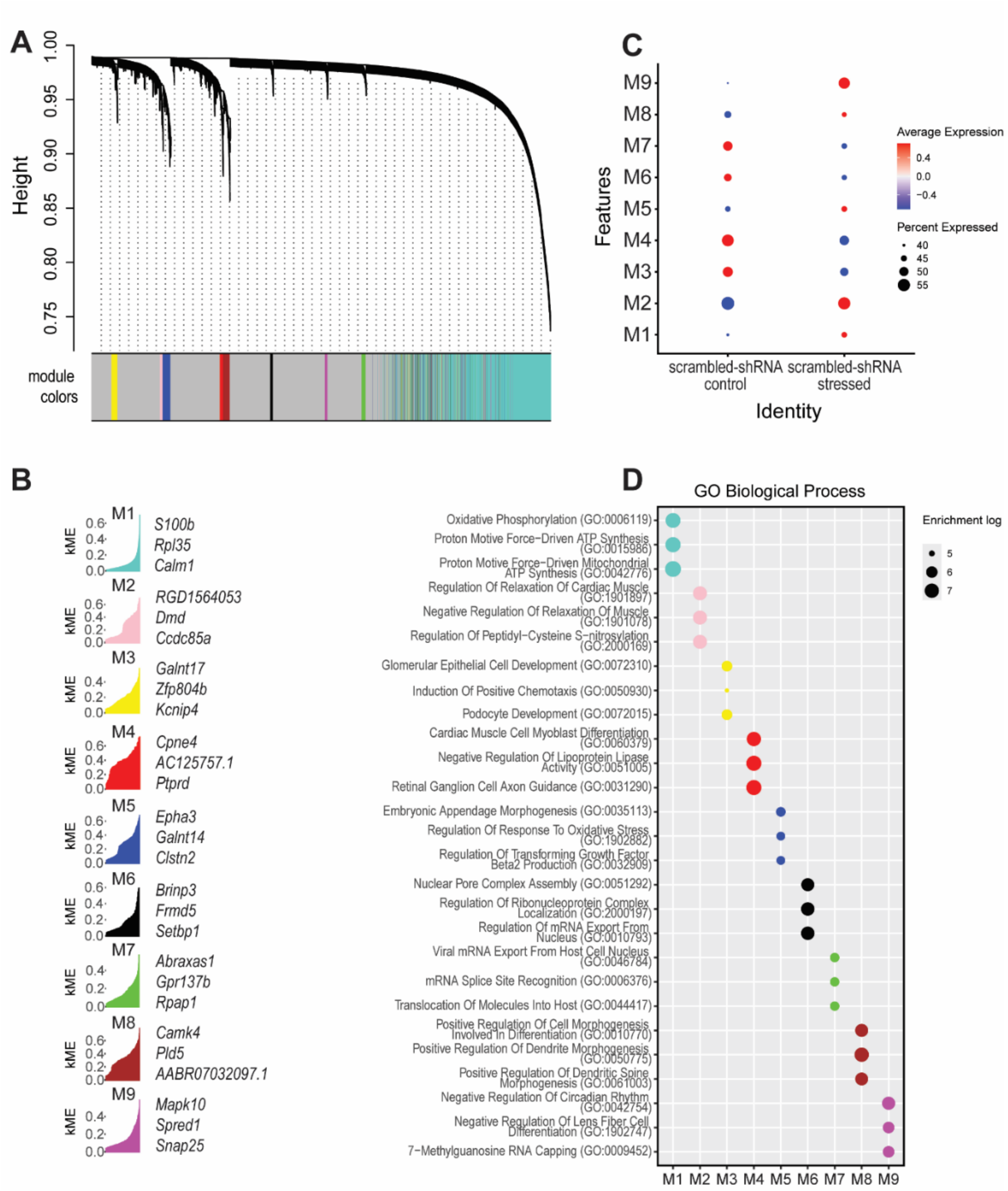


**Figure S16: Gene coexpression analysis of pyramidal neurons in Layer IV in the mPFC of scrambled-shRNA male rats exposed to acute stress.** (A) The dendrogram illustrates the network modular organization of differentially expressed genes comparing mPFC samples from stressed and control rats. (B) Nine modules were identified, with the top 3 eigengenes reported. (C) Module expression profiles with comparisons between knockdown and control samples, with red and blue representing up- and downregulation, respectively. (D) GO Biological Process pathway enrichment analysis of each coexpression module as calculated by hdWGCNA. The top three processes are indicated in each module.

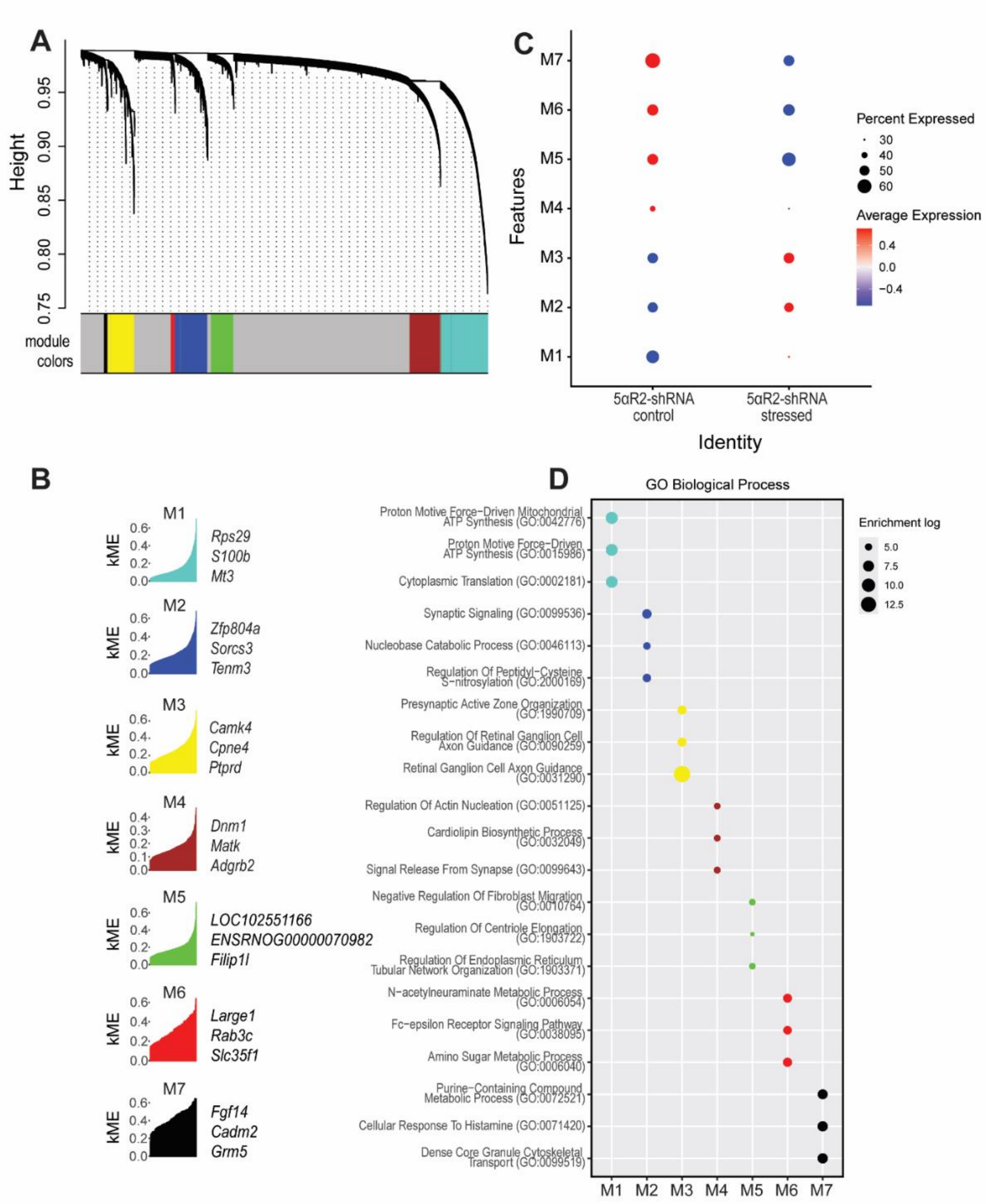


**Figure S17: Gene coexpression analysis of pyramidal neurons in Layer IV in the mPFC of 5αR2-shRNA male rats exposed to acute stress.** (A) The dendrogram illustrates the network modular organization of differentially expressed genes comparing mPFC samples from stressed and control rats. (B) Seven modules were identified, with the top 3 eigengenes reported. (C) Module expression profiles with comparisons between knockdown and control samples, with red and blue representing up- and downregulation, respectively. (D) GO Biological Process pathway enrichment analysis of each coexpression module as calculated by hdWGCNA. The top three processes are indicated in each module.

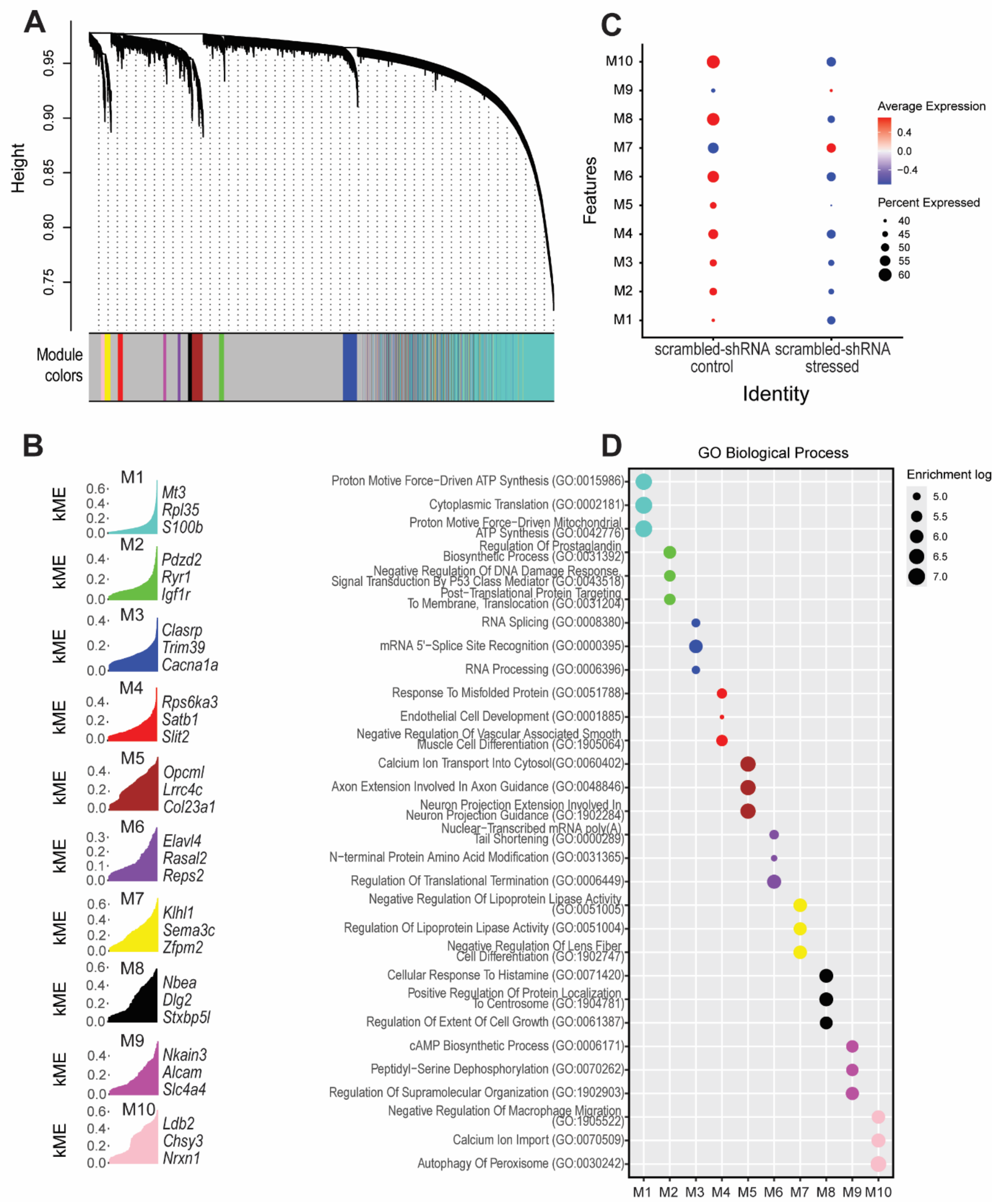


**Figure S18: Gene coexpression analysis of pyramidal neurons in Layer V in the mPFC of scrambled-shRNA male rats exposed to acute stress.** (A) The dendrogram illustrates the network modular organization of differentially expressed genes comparing mPFC samples from stressed and control rats. (B) Ten modules were identified, with the top 3 eigengenes reported. (C) Module expression profiles with comparisons between knockdown and control samples, with red and blue representing up- and downregulation, respectively. (D) GO Biological Process pathway enrichment analysis of each coexpression module as calculated by hdWGCNA. The top three processes are indicated in each module.

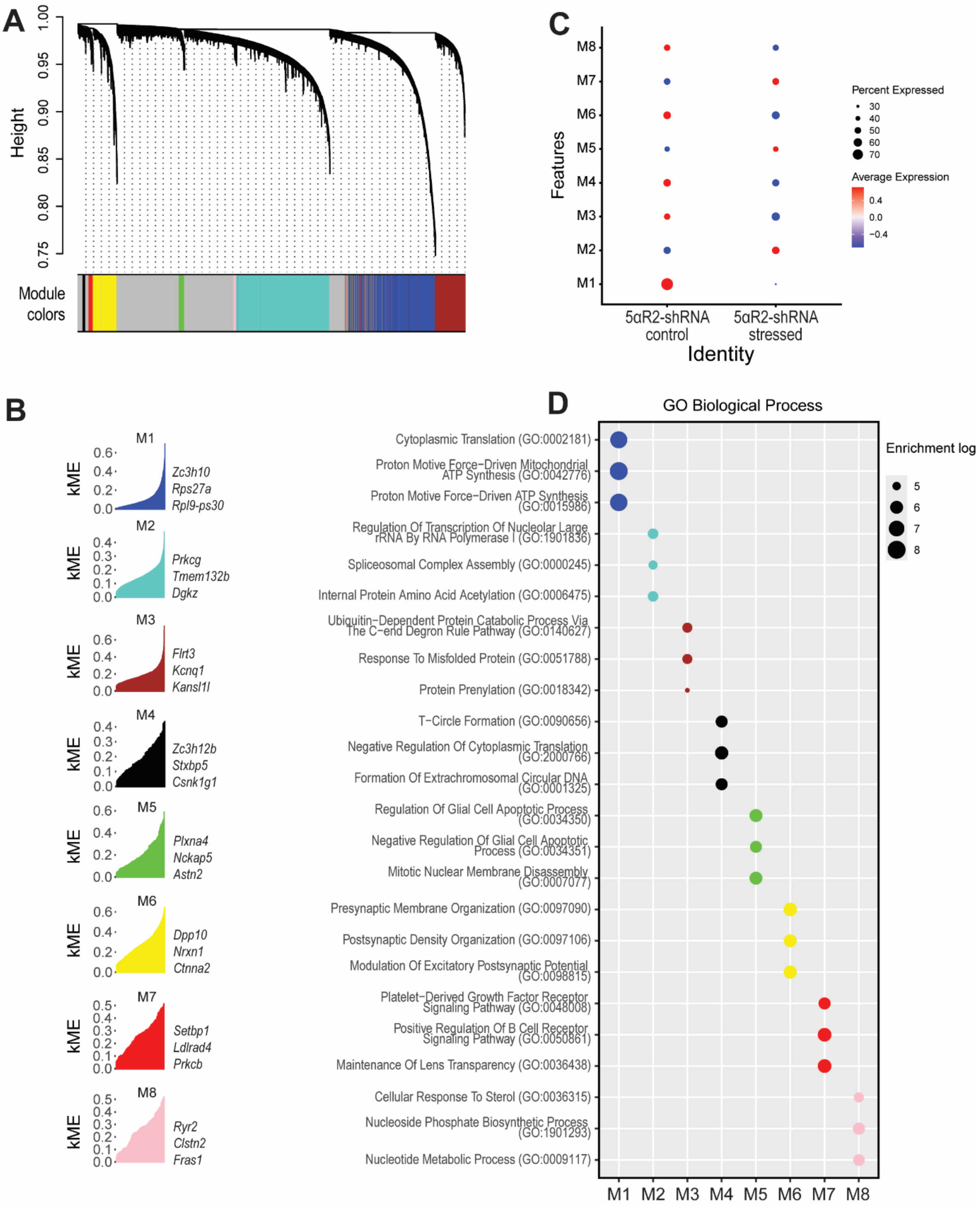


**Figure S19: Gene coexpression analysis of pyramidal neurons in Layer V in the mPFC of 5αR2-shRNA male rats exposed to acute stress.** (A) The dendrogram illustrates the network modular organization of differentially expressed genes comparing mPFC samples from stressed and control rats. (B) Eight modules were identified, with the top 3 eigengenes reported. (C) Module expression profiles with comparisons between knockdown and control samples, with red and blue representing up- and downregulation, respectively. (D) GO Biological Process pathway enrichment analysis of each coexpression module as calculated by hdWGCNA. The top three processes are indicated in each module.

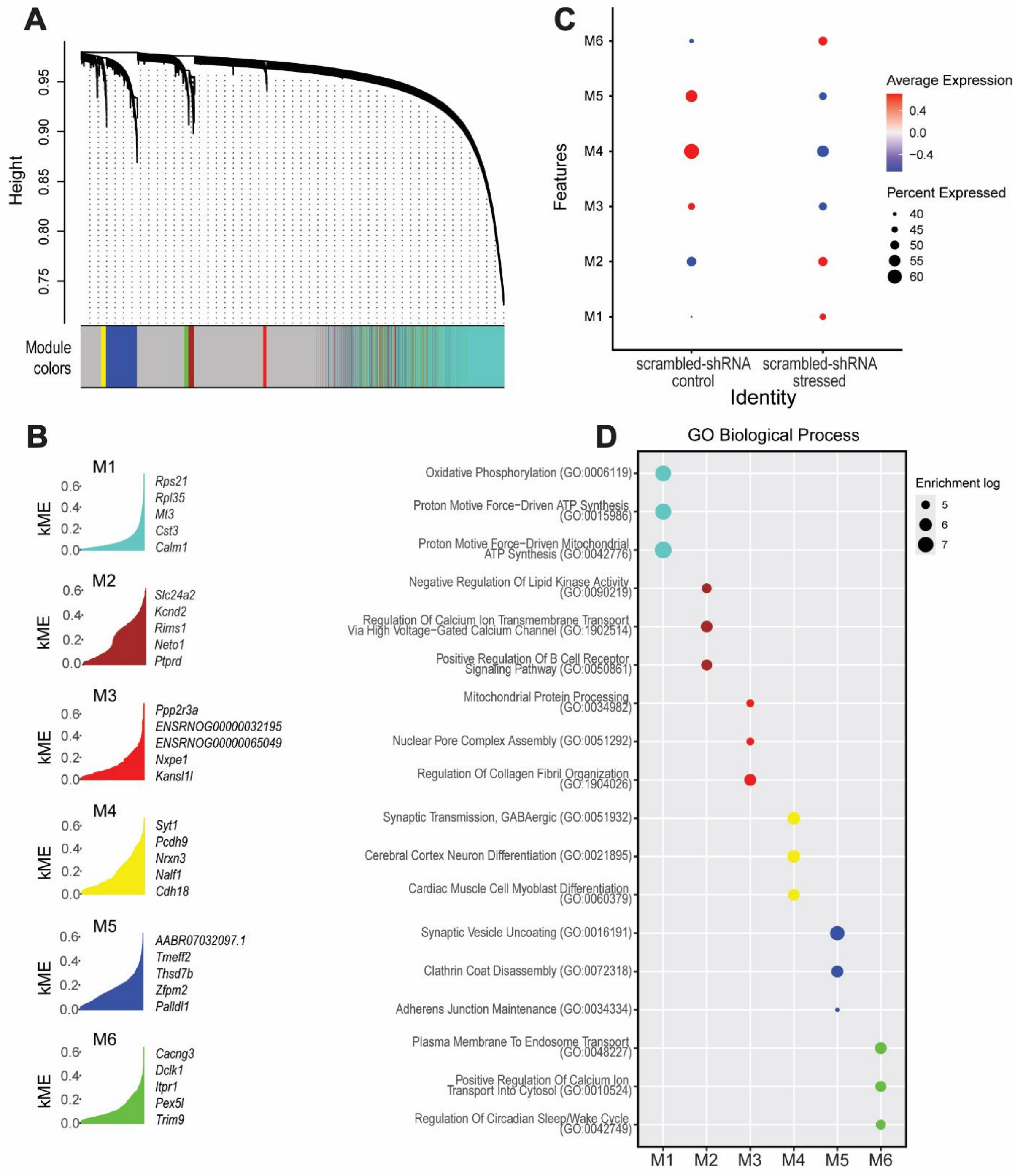


**Figure S20: Gene coexpression analysis of the first cluster of pyramidal neurons in Layer VI in the mPFC of scrambled-shRNA male rats exposed to acute stress.** (A) The dendrogram illustrates the network modular organization of differentially expressed genes comparing mPFC samples from stressed and control rats. (B) Six modules were identified, with the top 5 eigengenes reported. (C) Module expression profiles with comparisons between knockdown and control samples, with red and blue representing up- and downregulation, respectively. (D) GO Biological Process pathway enrichment analysis of each coexpression module as calculated by hdWGCNA. The top three processes are indicated in each module.

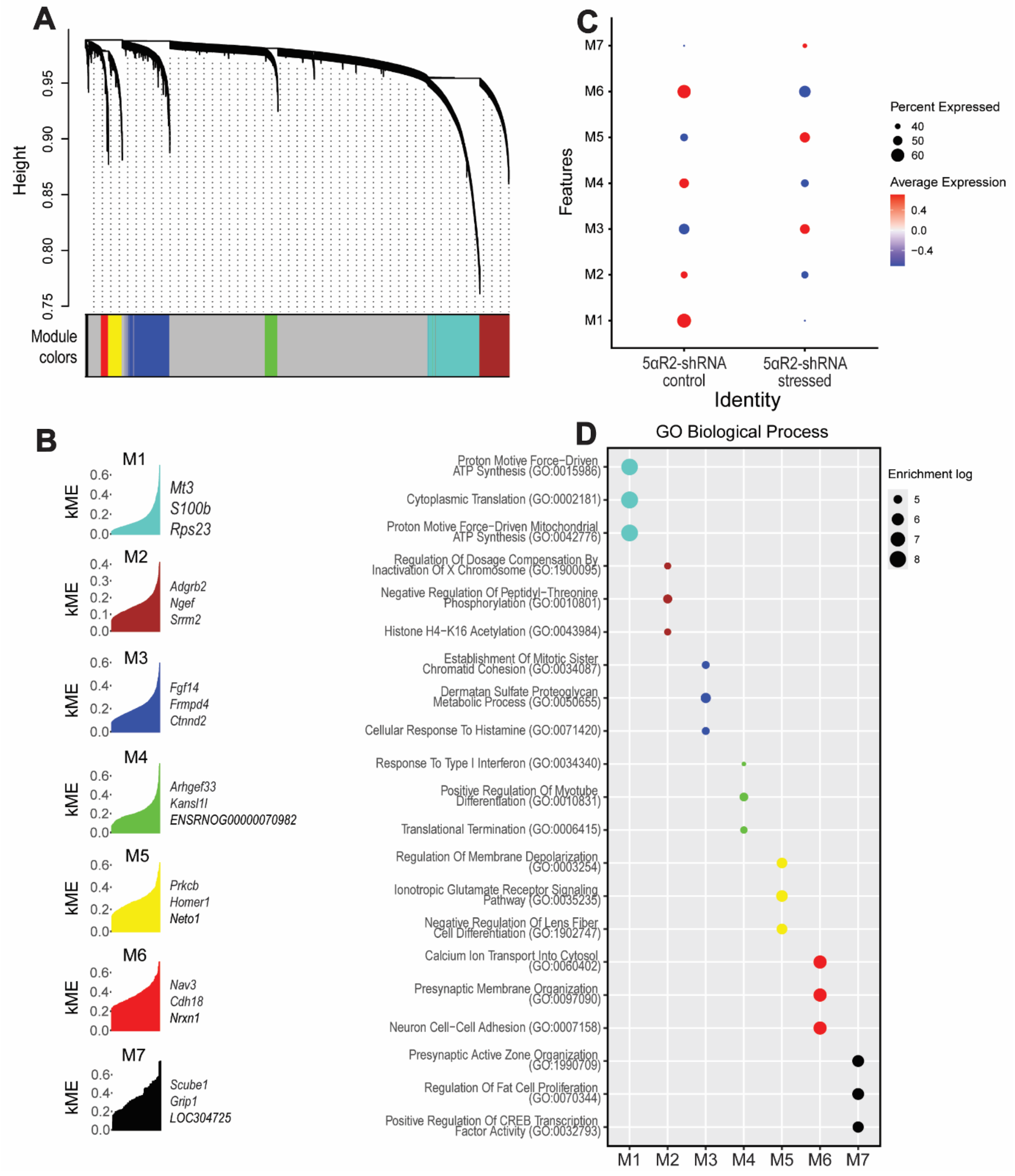


**Figure S21: Gene coexpression analysis of the first cluster of pyramidal neurons in Layer VI in the mPFC of 5αR2-shRNA male rats exposed to acute stress.** (A) The dendrogram illustrates the network modular organization of differentially expressed genes comparing mPFC samples from stressed and control rats. (B) Seven modules were identified, with the top 3 eigengenes reported. (C) Module expression profiles with comparisons between knockdown and control samples, with red and blue representing up- and downregulation, respectively. (D) GO Biological Process pathway enrichment analysis of each coexpression module as calculated by hdWGCNA. The top three processes are indicated in each module.

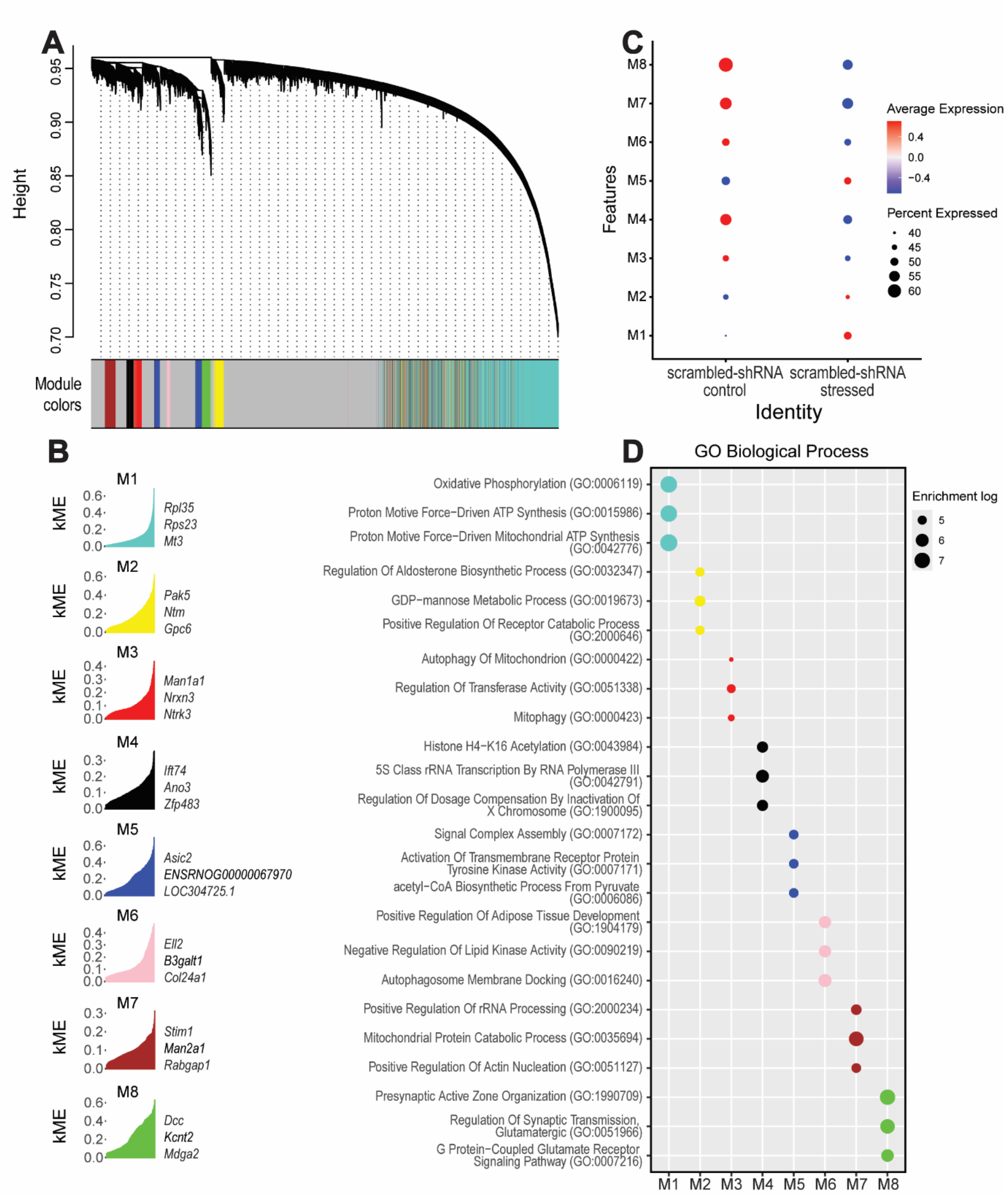


**Figure S22: Gene coexpression analysis of the second cluster of pyramidal neurons in Layer VI in the mPFC of scrambled-shRNA male rats exposed to acute stress.** (A) The dendrogram illustrates the network modular organization of differentially expressed genes comparing mPFC samples from stressed and control rats. (B) Eight modules were identified, with the top 3 eigengenes reported. (C) Module expression profiles with comparisons between knockdown and control samples, with red and blue representing up- and downregulation, respectively. (D) GO Biological Process pathway enrichment analysis of each coexpression module as calculated by hdWGCNA. The top three processes are indicated in each module.

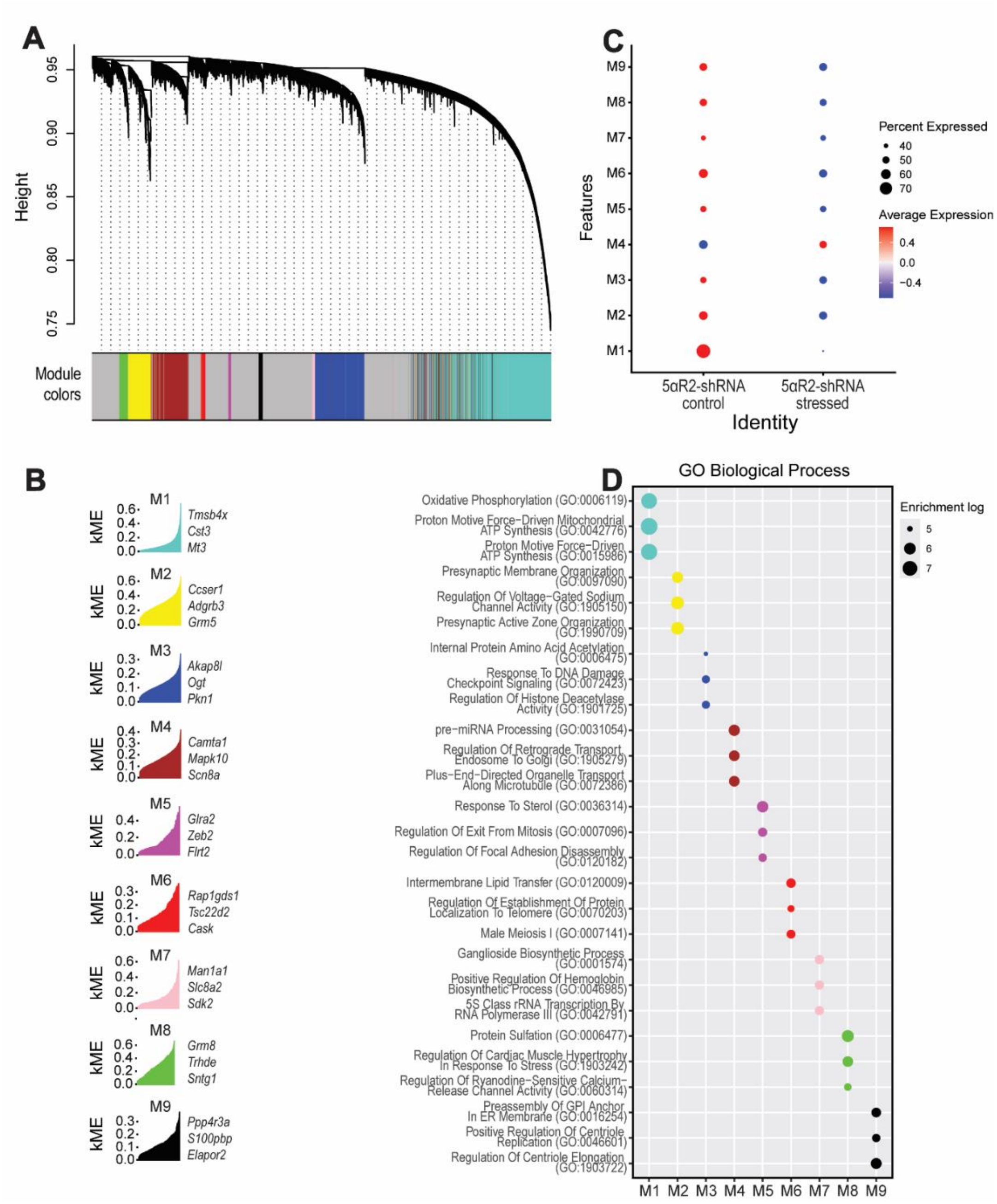


**Figure S23: Gene coexpression analysis of the second cluster of pyramidal neurons in Layer VI in the mPFC of 5αR2-shRNA male rats exposed to acute stress.** (A) The dendrogram illustrates the network modular organization of differentially expressed genes comparing mPFC samples from stressed and control rats. (B) Nind modules were identified, with the top 3 eigengenes reported. (C) Module expression profiles with comparisons between knockdown and control samples, with red and blue representing up- and downregulation, respectively. (D) GO Biological Process pathway enrichment analysis of each coexpression module as calculated by hdWGCNA. The top three processes are indicated in each module.

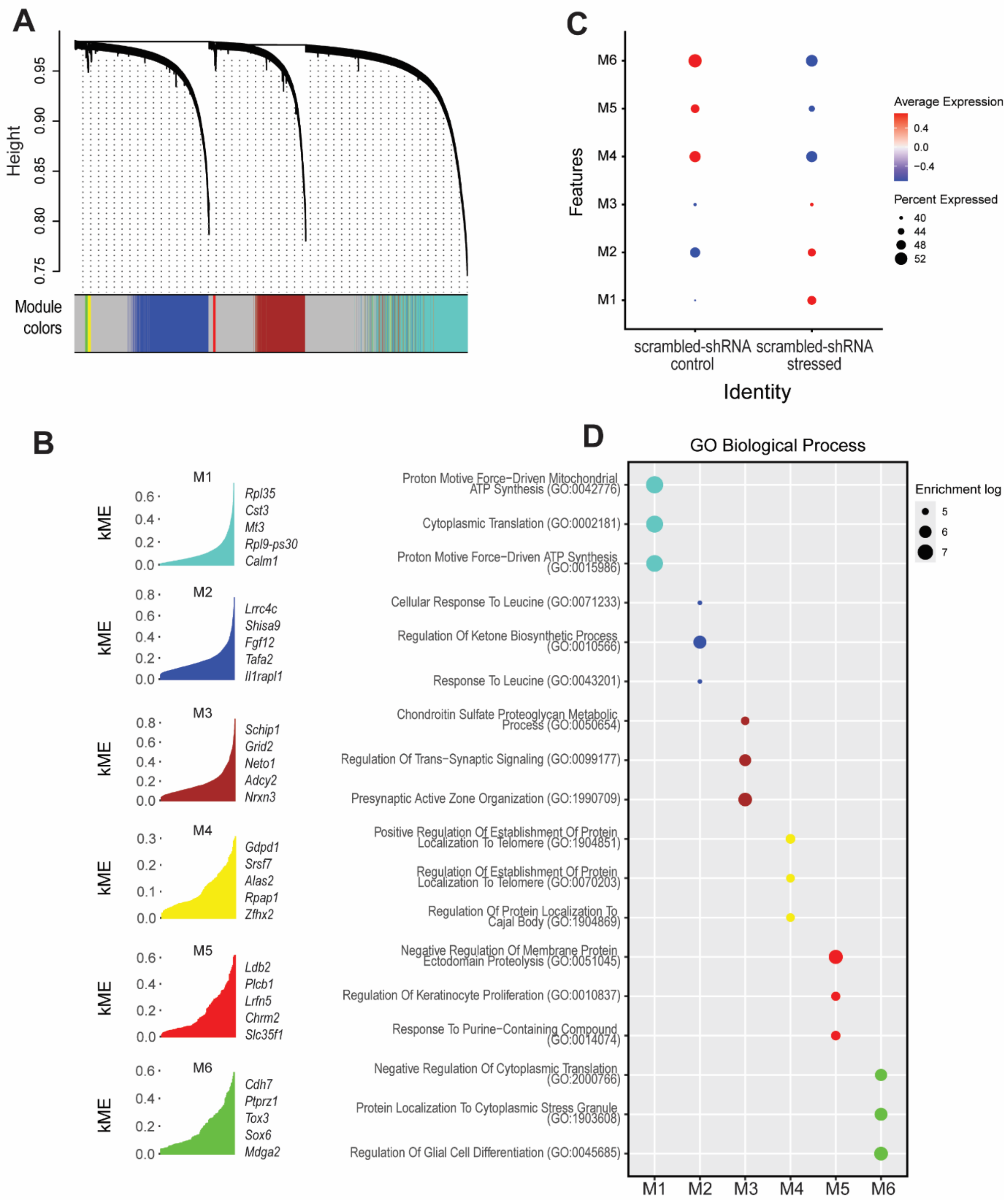


**Figure S24: Gene coexpression analysis of the VIP negative interneurons in the mPFC of scrambled-shRNA male rats exposed to acute stress.** (A) The dendrogram illustrates the network modular organization of differentially expressed genes comparing mPFC samples from stressed and control rats. (B) Six modules were identified, with the top 5 eigengenes reported. (C) Module expression profiles with comparisons between knockdown and control samples, with red and blue representing up- and downregulation, respectively. (D) GO Biological Process pathway enrichment analysis of each coexpression module as calculated by hdWGCNA. The top three processes are indicated in each module.

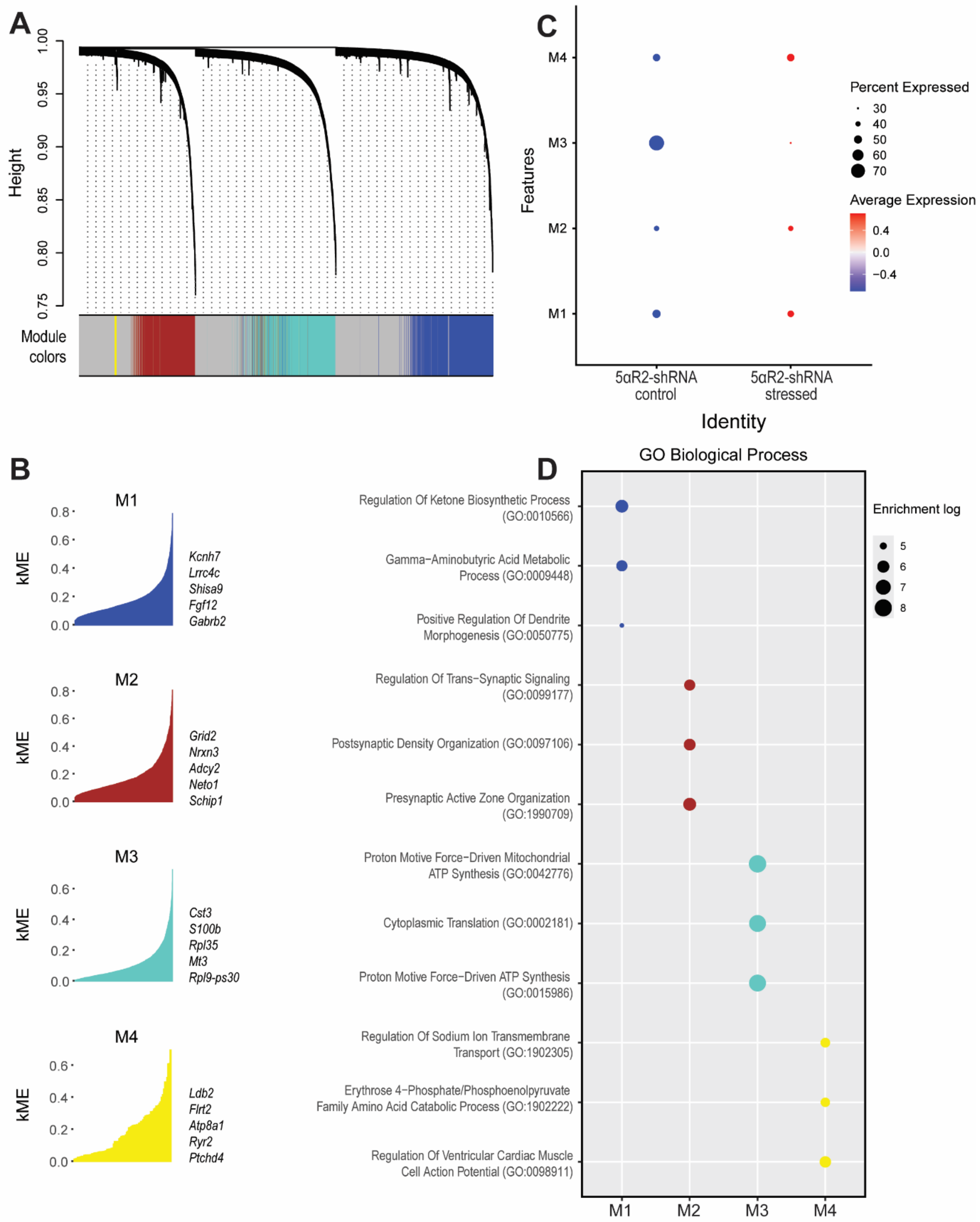


**Figure S25: Gene coexpression analysis of the VIP negative interneurons in the mPFC of 5αR2-shRNA male rats exposed to acute stress.** (A) The dendrogram illustrates the network modular organization of differentially expressed genes comparing mPFC samples from stressed and control rats. (B) Four modules were identified, with the top 5 eigengenes reported. (C) Module expression profiles with comparisons between knockdown and control samples, with red and blue representing up- and downregulation, respectively. (D) GO Biological Process pathway enrichment analysis of each coexpression module as calculated by hdWGCNA. The top three processes are indicated in each module.

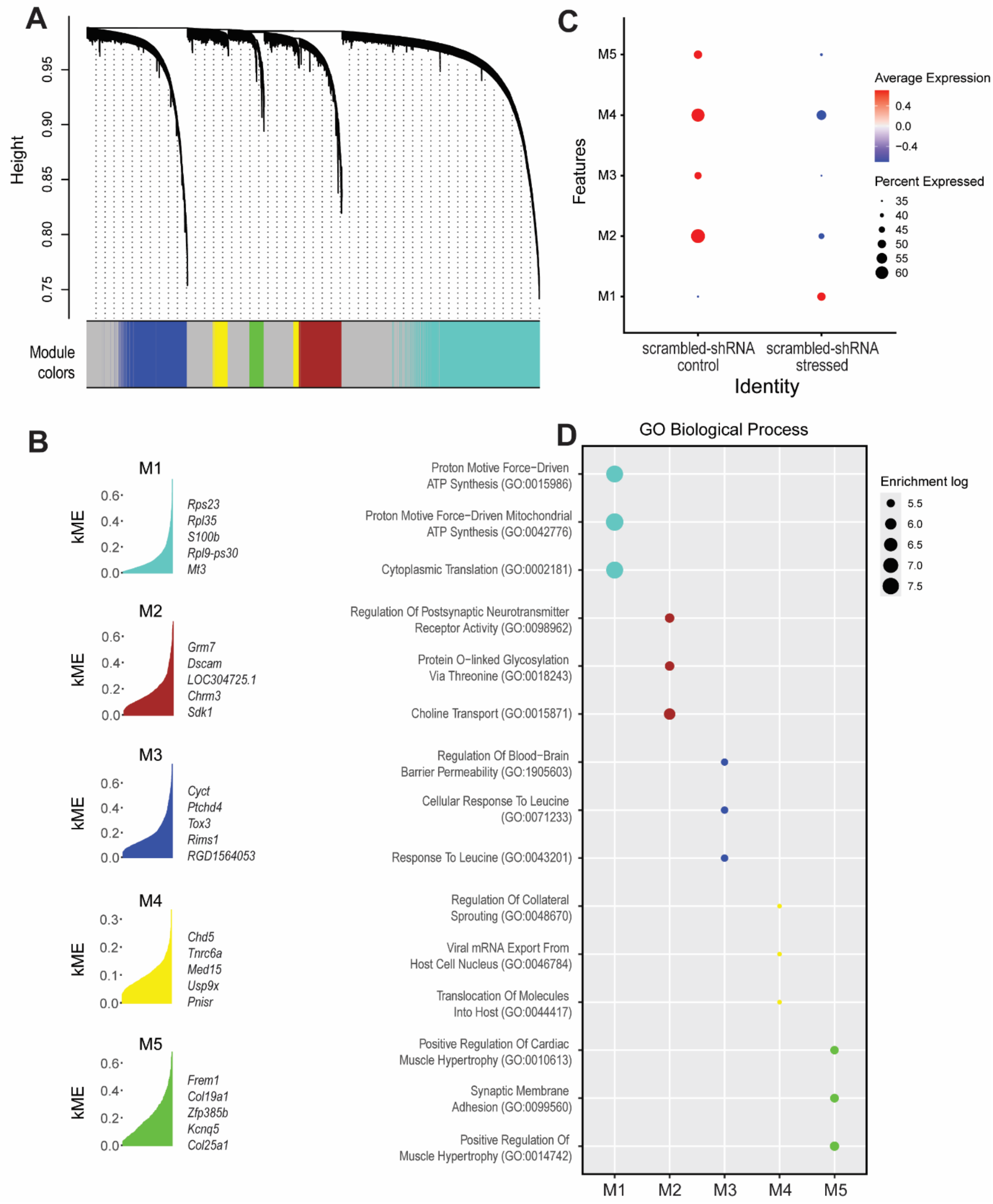


**Figure S26: Gene coexpression analysis of the VIP positive interneurons in the mPFC of scrambled-shRNA male rats exposed to acute stress.** (A) The dendrogram illustrates the network modular organization of differentially expressed genes comparing mPFC samples from stressed and control rats. (B) Five modules were identified, with the top 5 eigengenes reported. (C) Module expression profiles with comparisons between knockdown and control samples, with red and blue representing up- and downregulation, respectively. (D) GO Biological Process pathway enrichment analysis of each coexpression module as calculated by hdWGCNA. The top three processes are indicated in each module.

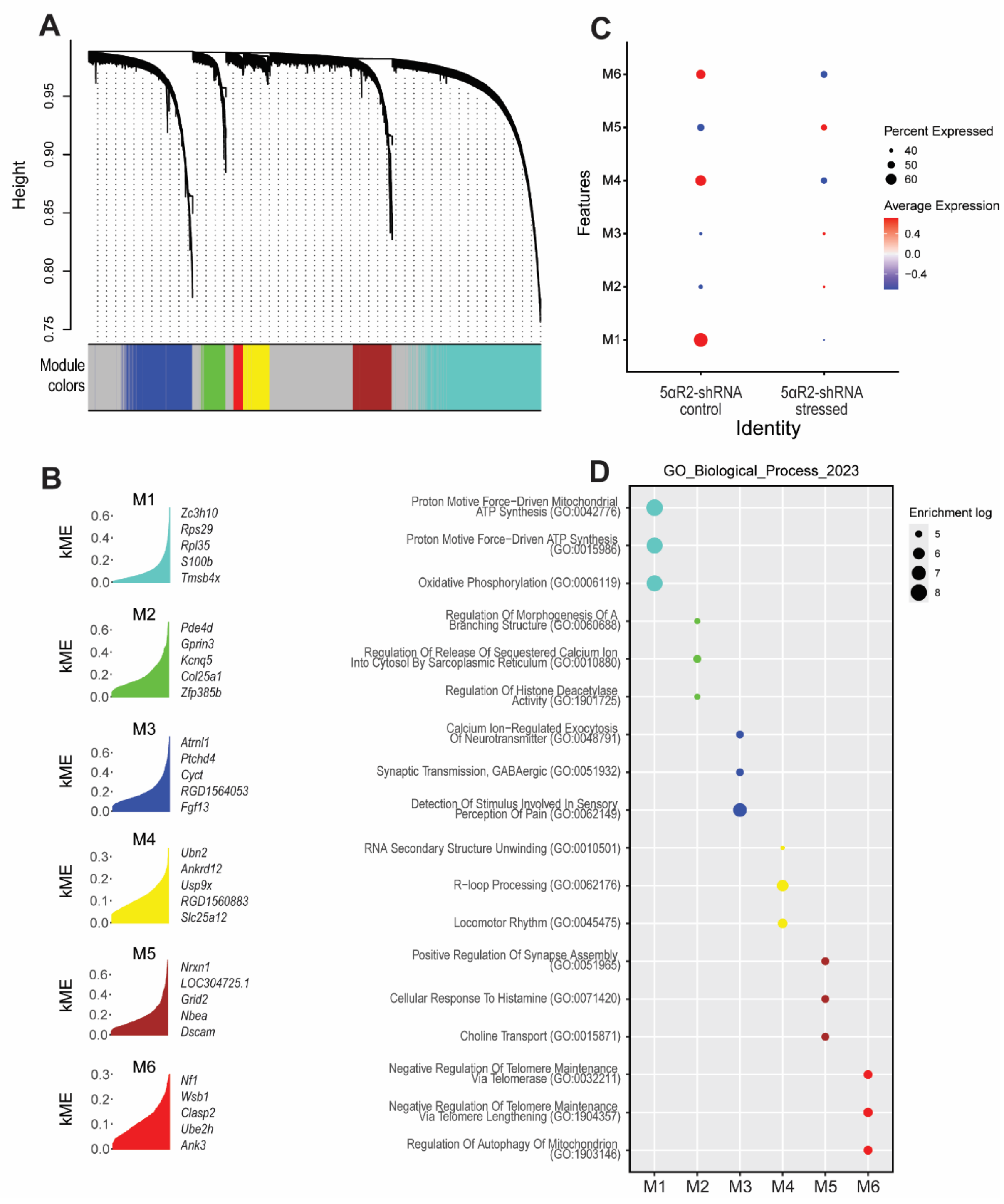


**Figure S27: Gene coexpression analysis of the VIP positive interneurons in the mPFC of 5αR2-shRNA male rats exposed to acute stress.** (A) The dendrogram illustrates the network modular organization of differentially expressed genes comparing mPFC samples from stressed and control rats. (B) Six modules were identified, with the top 5 eigengenes reported. (C) Module expression profiles with comparisons between knockdown and control samples, with red and blue representing up- and downregulation, respectively. (D) GO Biological Process pathway enrichment analysis of each coexpression module as calculated by hdWGCNA. The top three processes are indicated in each module.

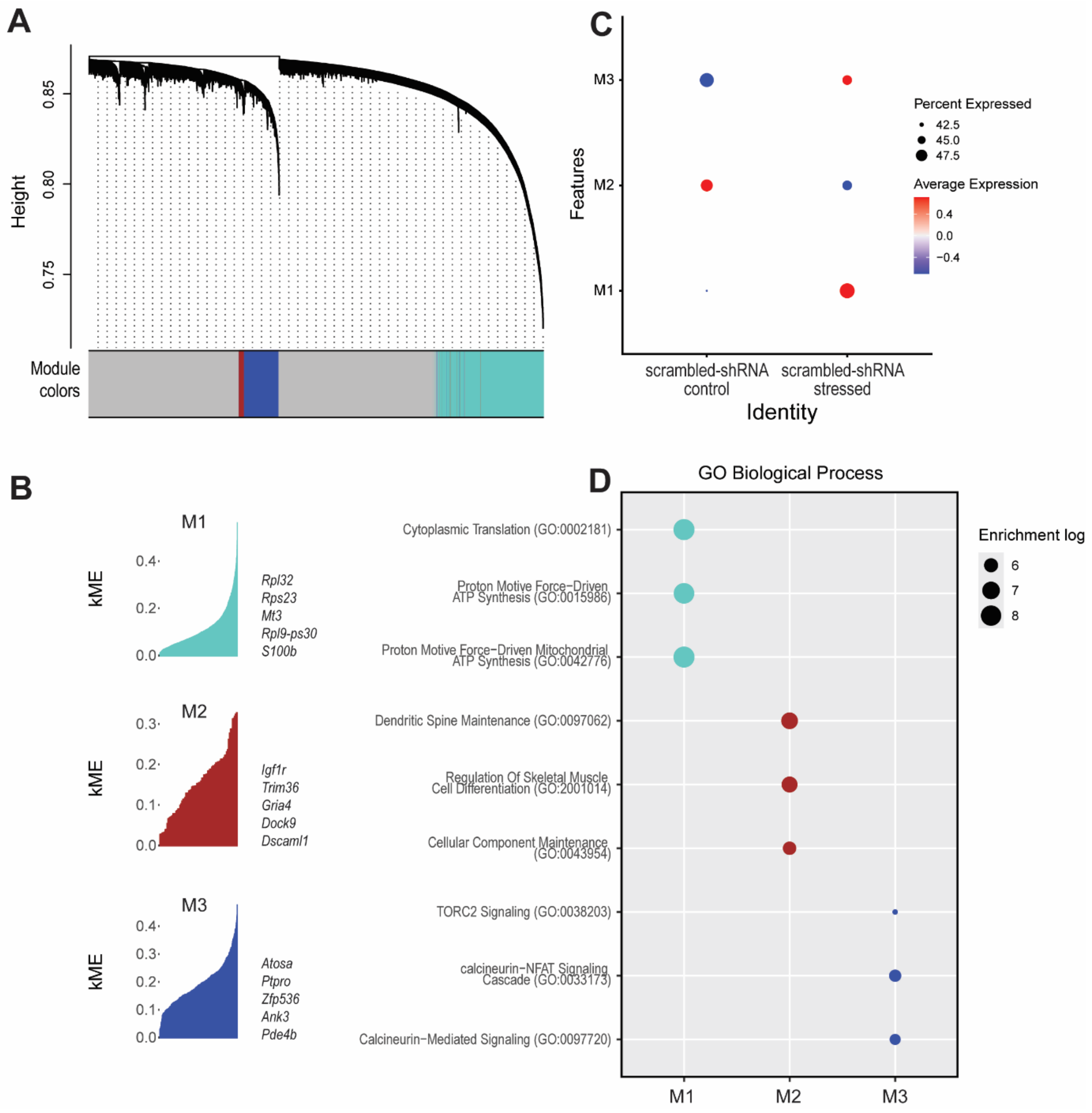


**Figure S28: Gene coexpression analysis of the oligodendendrocytes in the mPFC of scrambled-shRNA male rats exposed to acute stress.** (A) The dendrogram illustrates the network modular organization of differentially expressed genes comparing mPFC samples from stressed and control rats. (B) Three modules were identified, with the top 5 eigengenes reported. (C) Module expression profiles with comparisons between knockdown and control samples, with red and blue representing up- and downregulation, respectively. (D) GO Biological Process pathway enrichment analysis of each coexpression module as calculated by hdWGCNA. The top three processes are indicated in each module.

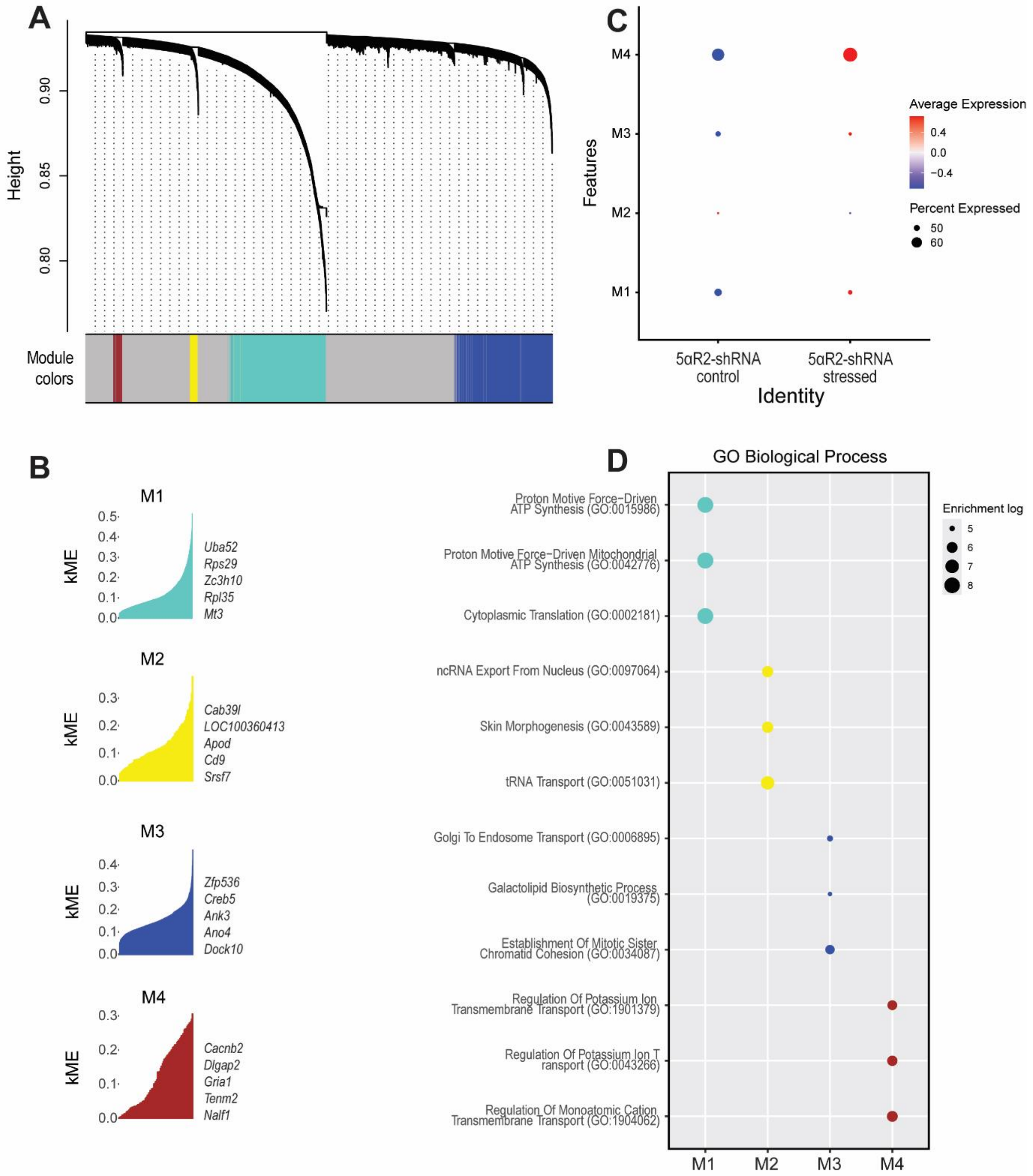


**Figure S29: Gene coexpression analysis of the oligodendendrocytes in the mPFC of 5αR2-shRNA male rats exposed to acute stress.** (A) The dendrogram illustrates the network modular organization of differentially expressed genes comparing mPFC samples from stressed and control rats. (B) Four modules were identified, with the top 5 eigengenes reported. (C) Module expression profiles with comparisons between knockdown and control samples, with red and blue representing up- and downregulation, respectively. (D) GO Biological Process pathway enrichment analysis of each coexpression module as calculated by hdWGCNA. The top three processes are indicated in each module.

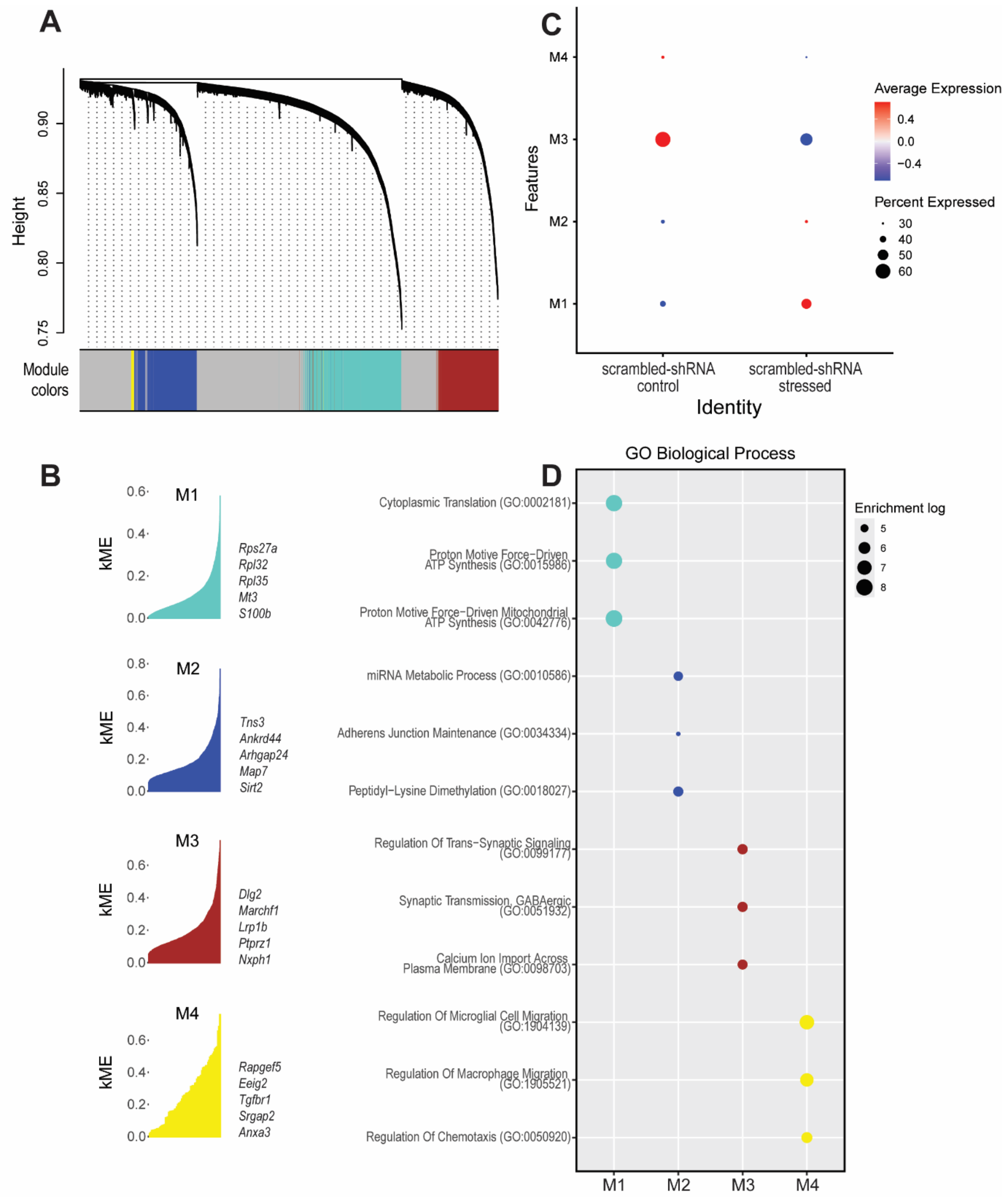


**Figure S30: Gene coexpression analysis of the OPC in the mPFC of scrambled-shRNA male rats exposed to acute stress.** (A) The dendrogram illustrates the network modular organization of differentially expressed genes comparing mPFC samples from stressed and control rats. (B) Four modules were identified, with the top 5 eigengenes reported. (C) Module expression profiles with comparisons between knockdown and control samples, with red and blue representing up- and downregulation, respectively. (D) GO Biological Process pathway enrichment analysis of each coexpression module as calculated by hdWGCNA. The top three processes are indicated in each module.

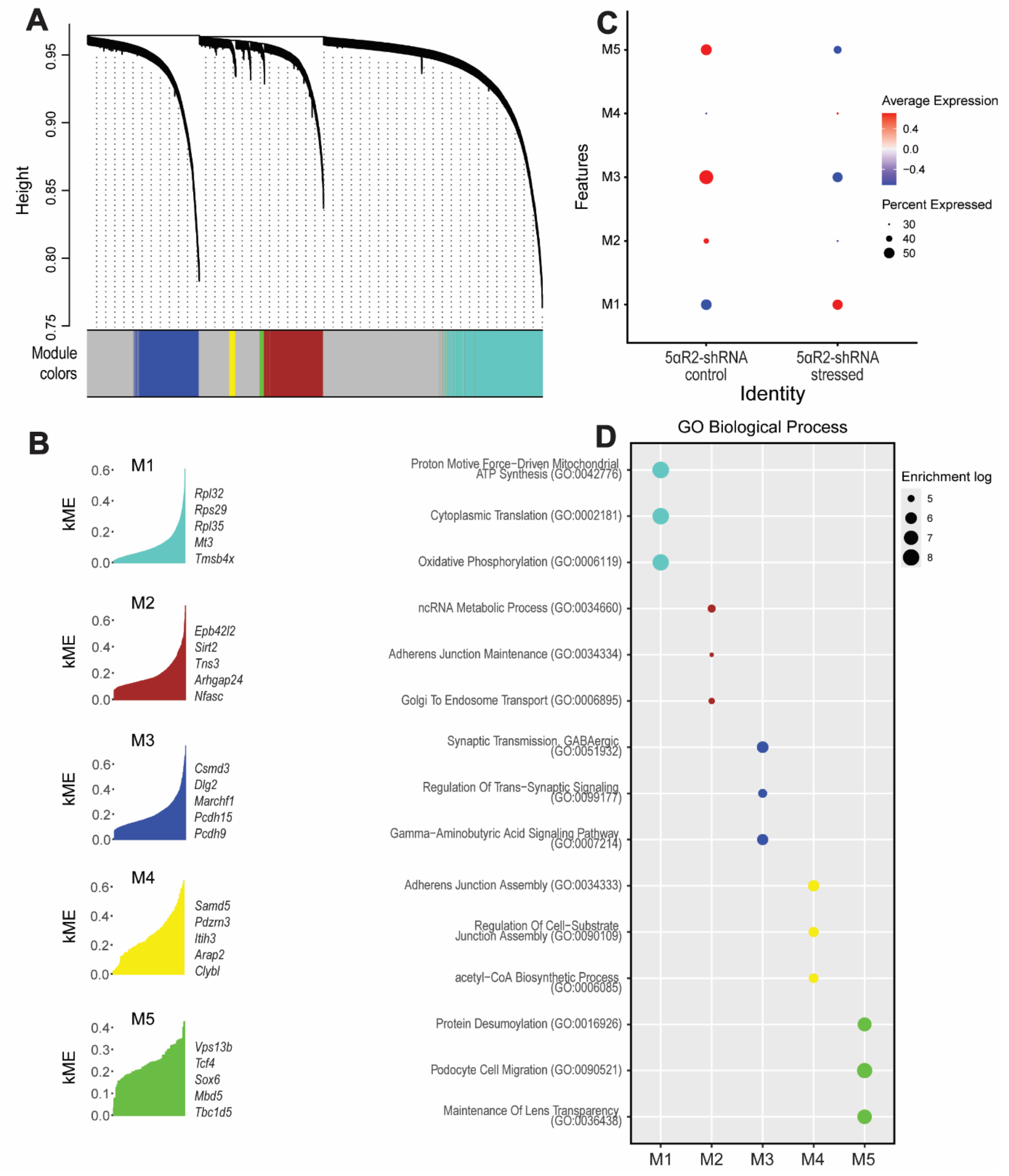


**Figure S31: Gene coexpression analysis of the OPC in the mPFC of 5αR2-shRNA male rats exposed to acute stress.** (A) The dendrogram illustrates the network modular organization of differentially expressed genes comparing mPFC samples from stressed and control rats. (B) Five modules were identified, with the top 5 eigengenes reported. (C) Module expression profiles with comparisons between knockdown and control samples, with red and blue representing up- and downregulation, respectively. (D) GO Biological Process pathway enrichment analysis of each coexpression module as calculated by hdWGCNA. The top three processes are indicated in each module.

**Supplementary Data**

Data S1: Data from single-nucleus (sn) transcriptomic studies of prefrontal cortices, depicting the list of downregulated genes in the comparison of scrambled-shRNA control vs 5αR2-shRNA control animals across all the different identified clusters and the GO analyses.

Data S2: Data from single-nucleus (sn) transcriptomic studies of prefrontal cortices, depicting the list of upregulated genes in the comparison of scrambled-shRNA control vs 5αR2-shRNA control animals across all the different identified clusters and the GO analyses.

Data S3: Data from single-nucleus (sn) transcriptomic studies of prefrontal cortices, depicting the list of differentially expressed genes (DEGs) in the comparison of scrambled-shRNA control vs 5αR2-shRNA control animals across all the different identified clusters and the GO analyses.

Data S4: Data from hdWGCNA defining the modules and the GO analyses of the oligodendrocyte cluster in the comparison of scrambled-shRNA control vs 5αR2-shRNA control animals.

Data S5: Data from hdWGCNA defining the modules and the GO analyses of the astrocyte cluster in the comparison of scrambled-shRNA control vs 5αR2-shRNA control animals.

Data S6: Data from hdWGCNA defining the modules and the GO analyses of the pyramidal neuron Layer V cluster in the comparison of scrambled-shRNA control vs 5αR2-shRNA control animals.

Data S7: Data from single-nucleus (sn) transcriptomic studies of prefrontal cortices, depicting the list of downregulated genes in the comparison of scrambled-shRNA control vs scrambled-shRNA stressed animals across all the different identified clusters and the GO analyses.

Data S8: Data from single-nucleus (sn) transcriptomic studies of prefrontal cortices, depicting the list of upregulated genes in the comparison of scrambled-shRNA control vs scrambled-shRNA stressed animals across all the different identified clusters and the GO analyses.

Data S9: Data from single-nucleus (sn) transcriptomic studies of prefrontal cortices, depicting the list of differentially expressed genes (DEGs) in the comparison of scrambled-shRNA control vs scrambled-shRNA stressed animals across all the different identified clusters and the GO analyses.

Data S10: Data from single-nucleus (sn) transcriptomic studies of prefrontal cortices, depicting the list of downregulated genes in the comparison of 5αR2-shRNA control vs 5αR2-shRNA stressed animals across all the different identified clusters and the GO analyses.

Data S11: Data from single-nucleus (sn) transcriptomic studies of prefrontal cortices, depicting the list of upregulated genes in the comparison of 5αR2-shRNA control vs 5αR2-shRNA stressed animals across all the different identified clusters and the GO analyses.

Data S12: Data from single-nucleus (sn) transcriptomic studies of prefrontal cortices, depicting the list of differentially expressed genes (DEGs) in the comparison of 5αR2-shRNA control vs 5αR2-shRNA stressed animals across all the different identified clusters and the GO analyses.

Data S13: Data from hdWGCNA defining the modules and the GO analyses of the pyramidal neuron Layer IV cluster in the comparison of scrambled-shRNA control vs scrambled-shRNA stressed animals.

Data S14: Data from hdWGCNA defining the modules and the GO analyses of the pyramidal neuron Layer IV cluster in the comparison of 5αR2-shRNA control vs 5αR2-shRNA stressed animals.

Data S15: Data from hdWGCNA defining the modules and the GO analyses of the pyramidal neuron Layer V cluster in the comparison of scrambled-shRNA control vs scrambled-shRNA stressed animals.

Data S16: Data from hdWGCNA defining the modules and the GO analyses of the pyramidal neuron Layer V cluster in the comparison of 5αR2-shRNA control vs 5αR2-shRNA stressed animals.

Data S17: Data from hdWGCNA defining the modules and the GO analyses of the pyramidal neuron Layer VIa cluster in the comparison of scrambled-shRNA control vs scrambled-shRNA stressed animals.

Data S18: Data from hdWGCNA defining the modules and the GO analyses of the pyramidal neuron Layer VIa cluster in the comparison of 5αR2-shRNA control vs 5αR2-shRNA stressed animals.

Data S19: Data from hdWGCNA defining the modules and the GO analyses of the pyramidal neuron Layer VIb cluster in the comparison of scrambled-shRNA control vs scrambled-shRNA stressed animals.

Data S20: Data from hdWGCNA defining the modules and the GO analyses of the pyramidal neuron Layer Vib cluster in the comparison of 5αR2-shRNA control vs 5αR2-shRNA stressed animals.

Data S21: Data from hdWGCNA defining the modules and the GO analyses of the VIP negative interneuron cluster in the comparison of scrambled-shRNA control vs scrambled-shRNA stressed animals.

Data S22: Data from hdWGCNA defining the modules and the GO analyses of the VIP negative interneuron cluster in the comparison of 5αR2-shRNA control vs 5αR2-shRNA stressed animals.

Data S23: Data from hdWGCNA defining the modules and the GO analyses of the VIP positive interneuron cluster in the comparison of scrambled-shRNA control vs scrambled-shRNA stressed animals.

Data S24: Data from hdWGCNA defining the modules and the GO analyses of the VIP positive interneuron cluster in the comparison of 5αR2-shRNA control vs 5αR2-shRNA stressed animals.

Data S25: Data from hdWGCNA defining the modules and the GO analyses of the oligodendrocyte cluster in the comparison of scrambled-shRNA control vs scrambled-shRNA stressed animals.

Data S26: Data from hdWGCNA defining the modules and the GO analyses of the oligodendrocyte cluster in the comparison of 5αR2-shRNA control vs 5αR2-shRNA stressed animals.

Data S27: Data from hdWGCNA defining the modules and the GO analyses of the OPC cluster in the comparison of scrambled-shRNA control vs scrambled-shRNA stressed animals.

Data S28: Data from hdWGCNA defining the modules and the GO analyses of the OPC cluster in the comparison of 5αR2-shRNA control vs 5αR2-shRNA stressed animals.
